# The not-so-great speciator: Systematics and species limits in a rapid radiation, the Asiatic white-eye complex (*Zosterops spp.*)

**DOI:** 10.64898/2026.02.25.708059

**Authors:** Herman L. Mays, Bailey D. McKay, Isao Nishiumi, Cheng-Te Yao, Fa-Sheng Zou, Madeline Boyd, Devon DeRaad, Ruey-Shing Lin, Kazuto Kawakami, Chang-Hoe Kim, Laura Kubatko, Robert Moyle

## Abstract

Here we untangle the systematics of the Asiatic white-eye complex (*Zosterops* spp.) to better understand the early stages of a recent island radiation. We adopt an integrative approach involving allelic data, genome-scale single nucleotide polymorphisms (SNPs), and museum-based morphometrics coupled with a comprehensive sampling to provide the most holistic understanding of the group to date. The island lineages of Asiatic white-eyes across Indonesia, the Philippines, East Asia, the adjacent oceanic islands of the Western Pacific underwent a deep split separating *Zosterops everetti* and *Z. nigrorum* in the Phillippines from a very rapid radiation including *Z. japonicus*, *Z. meyeni*, and *Z. montanus* in the Philippines, Japan, and Indonesia. *Z. nigrorum catarmanensis* on Camiguin South in the Philippines was found to be nested within *Z. montanus* and a species limit between *Z. nigrorum* populations on Panay and Luzon was strongly supported. Phylogenetic splits and population structure were detected within the clade containing *Z. japonicus*, *Z. meyeni*, and *Z. montanus*. A well-supported split separates a northern group including Northern Philippines *Z. montanus* subspecies, *Z. meyeni*, and *Z. japonicus* from the southerly *Z. montanus* taxa. This creates a paraphyletic *Z. montanus*. However, based on speciation rates within the broader Asiatic white-eye clade this break likely does not yet represent evolutionarily independent species lineages. Morphological evolution is taking place within the Asiatic white-eyes especially within the robust, large-billed subspecies of *Z. japonicus* on the oceanic islands of Japan and in the newly identified yellow-morph of *Z. montanus* on Camiguin South.

## 1. Introduction

### 1.1 Synthesis

Species radiations across island archipelagos are powerful exemplars of numerous evolutionary processes including speciation, reproductive isolation, genetic drift, introgression, and natural selection (Whittaker et al., 2017). Oceanic archipelagos, such as the Galapagos and Hawaiian Islands, have earned a central role in our understanding of evolutionary radiations (Losos & Ricklefs 2009; Cerca et al., 2023). Continental islands too have contributed to our knowledge of evolutionary radiations. For example, Anolis lizards across the Greater Antilles have revealed a story of parallel evolution and convergent species assemblies arising independently *in situ* on different islands (e.g., Patton et al., 2021). Many of these examples of island radiations are in a comparatively late stage and the processes responsible for their origins may be obscured by time. What’s more, many island radiations are spread across a mix of continental and oceanic islands and may be largely non-adaptive exhibiting little phenotypic divergence and ecological specialization (Czekanski-Moir & Rundell 2019).

Speciation is nascent in the earliest stages of rapid, non-adaptive radiations and systematists are therefore forced to deal with an ambiguous boundary between populations and species. This intersection is fraught with complications (Carstens et al., 2013; Huang 2020). This is especially true in cases where fluctuating, strong selection creates local ecomorphs that may be easily confused with species (McKay & Zink 2015; Hill & Powers 2021), when differentiation in a handful of genes is buried in a genomic background with little or no phylogenetic signal (Helmkampf et al., 2025), or where continuous phenotypic clines and isolation-by-distance make fractioning into discrete lineages appear arbitrary (Hillis 2019; Turbek et al., 2023). Distinguishing species from structured populations may be extraordinarily difficult (Reeve et al., 2023; Kearns et al., 2024) but is the first step in understanding the beginnings of an evolutionary radiation. Studying taxa originating at different times across continental and oceanic islands through dense geographic sampling is important if we are to understand those processes responsible for island radiations.

### 1.2 Focal system

One group that has long confounded systematists are the white-eyes (*Zosterops spp.*). Of the 150 recognized species of white-eyes, yuhinas, and their allies (Aves: Zosteropidae), distributed throughout Africa, continental Asia, Indonesia, Australasia, the Philippines, Oceania, and the Indian Ocean, 111 belong to the genus *Zosterops* (Winkler et al., 2020). This diversity has arisen since the Pliocene-Pleistocene boundary, placing the genus among the highest known per-lineage rates of vertebrate diversification. The diversification rate of *Zosterops* white-eyes exceeds that of Hawaiian honeycreepers and Galapagos finches and rivals that found in African Rift Valley cichlids (Moyle et al., 2009; Melo et al., 2011; Jetz et al., 2012).

In their classic study of island speciation Diamond *et al*. (1976) included the *Zosterops* [*griseotincta*] superspecies complex among a collection of birds in the Solomon Islands dubbed the “great speciators”. These species presented a paradox. They disperse well enough to colonize islands but not so well as to prevent isolation. Since, the “great speciator” moniker has been used consistently to refer to *Zosterops* white-eyes as a whole (Moyle et al., 2009; Cornetti et al., 2015; Cowles & Uy 2019; Lim et al., 2019; Gwee et al., 2020; Manthey et al., 2020; Oliveros et al., 2021; Vinciguerra et al., 2023). Because the island white-eyes so clearly embody this “great speciator” paradox, it is understandable that they are natural subjects for the study of evolutionary radiations.

This rapid radiation has generated no shortage of taxonomic challenges. Mayr (1965) and others (Moyle et al., 2009; Melo et al., 2011; Round et al., 2017; Wells 2017; Wells 2017) have noted that morphological similarity does not necessarily coincide with evolutionary relatedness within *Zosterops*. Distantly related and geographically separate and widespread species may be more phenotypically similar than adjacent, closely related, and narrowly distributed species. Genetic evidence has dashed many an avian systematist’s hypotheses regarding *Zosterops* and a series of recent studies have resulted in significant rearrangement (Melo et al., 2011; Wells 2017; Wells 2017; Lim et al., 2019; Clements et al., 2024).

### 1.3 Background

The *Zosterops* species distributed across continental East Asia and the extensive island archipelagos of the region including Taiwan, the Philippines, the Ryukyus, the main Japanese islands of Honshu, Hokkaido, Shikoku, and Kyushu, and the oceanic island chains of the Daito, Izu, Ogasawara, and Volcano Islands are among the most recent radiations within the genus. *Z. japonicus* once encompassed nine subspecies, including *Z. japonicus simplex* found throughout South and Southeast Mainland China, Northern Indochina, and Taiwan, and *Z. japonicus hainanus* on the island of Hainan, with the remaining subspecies distributed throughout the islands of Japan and the Korean Peninsula (Clements et al., 2007). Once again molecular data disrupted the taxonomic status quo and subspecies in Mainland China, Taiwan, and Hainan were found to be genetically distinct from the rest of *Z. japonicus*. Despite their phenotypic similarity and geographic proximity, these taxa are not sister to *Z. japonicus* in Japan and Korea and their status was elevated to *Z. simplex*. Populations on the Thai-Malay Peninsula, Enggano Island, Borneo, and Sumatra that were taxonomically shuffled around between species are now included in a more broadly defined *Z. simplex* (Moyle et al., 2009; Round et al., 2017; Lim et al., 2019; Clements et al., 2024).

The Philippines hosts four species of *Zosterops* that have not been extensively studied. The more southerly distributed *Z. everetti* and the more northern *Z. nigrorum* are the most divergent among the Philippine white-eyes while *Z. meyeni* and *Z. montanus* are variable in appearance but resemble *Z. japonicus* in Japan. While the species status of *Z. everetti* and *Z. nigrorum* have seldom been questioned, outside of whether to include other populations outside the Philippines with *Z. everetti* (Wells 2017), doubts about the species status of *Z. montanus* and *Z. meyeni* persist. Jones and Kennedy’s (2008) analysis of the mitochondrial DNA (mtDNA) ND2 locus discovered that *Z. montanus* is paraphyletic with respect to *Z. japonicus* with the northern *Z. montanus* subspecies of *whiteheadi* and *halconensis* more closely related to *Z. japonicus* than to the other *Z. montanus* subspecies in the Visayas and on Mindanao. Later work incorporating mtDNA sequences also revealed a paraphyletic *Z. montanus* with respect to *Z. japonicus* confirming the same close affiliation between *Z. montanus whiteheadi* on Luzon and *Z. japonicus* in Japan (Lim et al., 2019). Illustrative of this ongoing issue, Lim *et al*. (2019) called *Z. montanus*, together with *Z. japonicus* and *Z. palpebrosus*, a taxonomic wastebasket and used these data to successfully argue for merging *Z. montanus* and *Z. japonicus* into a now broadly distributed *Z. japonicus* ranging from Java to Hokkaido. This lumped *Z. japonicus* taxon à la Lim *et al*. (2019) has since been formally recognized (Van Riper & van Balen 2020; Clements et al., 2021; Clements et al., 2024).

Considering the recent lumping of *Z. montanus* and *Z. japonicus* across the island archipelagos of Indonesia, the Philippines, and Japan one might say the group falls short of their reputation as “great speciators”. However, sampling the Asiatic white-eyes, as is true for any geographically widespread group, is difficult. For this reason, prior studies often relied on limited geographic representation. For instance, Jones and Kennedy’s (2008) study focused on *Z. montanus* in the Philippines and utilized only a single sequence from *Z. japonicus*. Price *et al*. (2014) found a relatively deep sister relationship between *Z. japonicus* and *Z. montanus* but their *Z. japonicus* sampling was from Mainland China later shown to be highly divergent from those birds in the Japanese Archipelago. Lim *et al*. (2019) analyzed nine samples of *Z. japonicus*, all from the nominate subspecies, and 15 samples from *Z. montanus*, representing five of the nine subspecies. Additionally, this study did not include samples from *Z. meyeni*. Gwee *et al*. (2020) employed a larger molecular dataset to tackle the *Zosterops* genus as a whole. However, the study was limited to a single sample from *Z. meyeni*, two *Z. japonicus* samples, one in Japan and the other from the introduced population in Hawaii, and five *Z. montanus*, including four from Indonesia and a single sample from the Philippines. Together these samples represented only three of the subspecies in the combined *Z. japonicus/montanus* clade. Oliveros *et al*. (2021) used ultra-conserved elements (UCEs) to produce a family-level phylogeny for *Zosteropidae* and as such sampled broadly but not deeply with only a single sample from each of the Asiatic white-eye species, excluding any samples from Japan. Because both Gwee et al. (2020) and Oliveros *et al*. (2021) were focused on the deeper relationships within *Zosterops* the authors were able to draw few conclusions about species limits within the Asiatic white-eye group.

Thus, tackling the systematics of the Asiatic white-eyes has been akin to the Buddha’s parable of the blind men and the elephant in that different studies have sampled sparsely from different populations each revealing only part of the picture. Prior work was based on a limited number of loci (Jones & Kennedy 2008; Round et al., 2017; Lim et al., 2019) or employed large genetic datasets but with limited sampling (Gwee et al., 2020; Oliveros et al., 2021). None of these prior studies have combined genome-wide genetic data with broad sampling and phenotypic data. With an integrative approach and increased sampling this study reveals a little more of the whole elephant. Here we studied the Asiatic white-eye complex based on comprehensive geographic sampling and a combined dataset of Sanger sequenced loci including two mtDNA loci (COI and ND2) and six nuclear protein coding genes, genome-wide single nucleotide polymorphism (SNP) data derived from restriction site associated DNA sequencing (RADseq), and linear morphometric data from museum study skins. The results of this integrative study represent the most holistic understanding to date of the systematics of Asiatic white-eyes and reveal the patterns of diversification in a very recent evolutionary radiation at the intersection of population structure and species.

## 2. Methods

### 2.1 Genetic sampling

Sampling included specimen vouchered tissue samples and museum archived blood sampling from individual banded birds in the field. Samples were sourced from the American Museum of Natural History (New York, NY, USA), Cincinnati Museum Center (Cincinnati, OH, USA), Field Museum of Natural History (Chicago, IL, USA), Institute of Zoology of the Guangdong Academy of Sciences (Guangzhou, People’s Republic of China), University of Kansas Biodiversity Institute and Natural History Museum (Lawrence, KS, USA), Natural History Museum of Los Angeles County (Los Angeles, CA, USA), National Museum of Nature and Science (Tokyo, Japan), Smithsonian National Museum of Natural History (Washington D.C., USA), and the University of Washington Burke Museum (Seattle, WA, USA). For the sake of clarity, we treat *Z. japonicus* and *Z. montanus* according to the former taxonomic arrangement (i.e., as separate species) as we test various species delimitation hypotheses. Our sampling includes multiple individuals within each of five out of the six recognized subspecies for *Z. japonicus* and seven out of the nine subspecies within *Z. montanus*, including sampling from all the recognized *Z. montanus* subspecies within the Philippines (Van Riper & van Balen 2020). A total of 208 genomic DNA (gDNA) samples were used in this study with 159 represented in the Sanger sequencing dataset, 124 represented in the filtered SNP dataset, and 75 samples overlapping both genetic datasets. A map showing all sampling locations may be found in Figure 1 and detailed sampling locations for the Ryukyu Islands and the Philippines may be found in Supplementary figure 1. A complete list of samples used in both genetic datasets may be found in Supplementary table 1 and additional details are available in the “Supplementary: Sampling maps” document (https://hermmays.github.io/Zosterops1/ZosteropsMaps.html).

**Figure 1:**
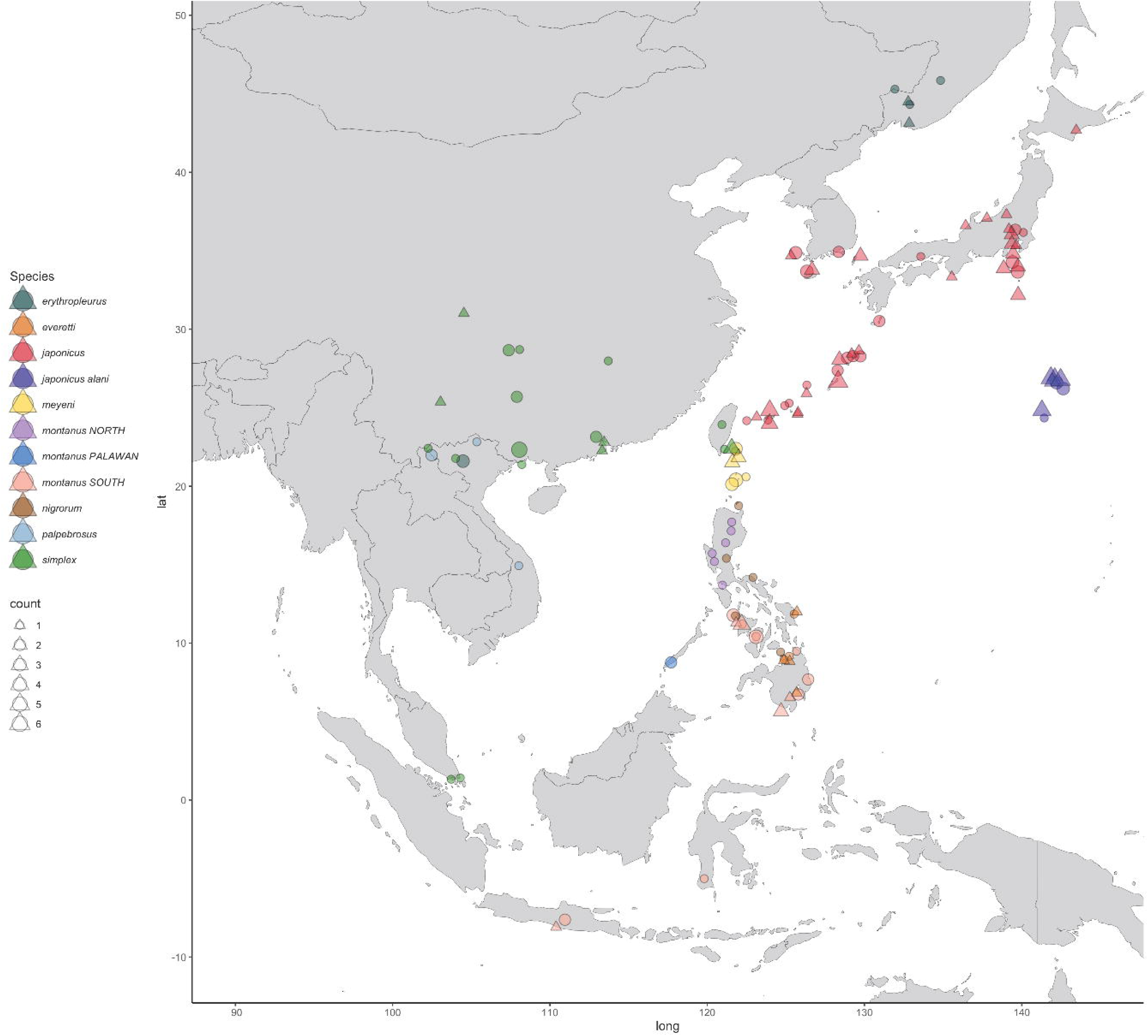
Localities for 208 Zosterops samples in this this study. 159 samples are represented by Sanger sequence data and 124 samples are represented by SNP data. 75 samples are represented in both the Sanger sequence and SNP datasets. Circles denote samples with SNP data alone or both SNP data and Sanger sequencing data. Triangles indicate those samples only with Sanger sequencing data. Sampling is color coded according to the legend by a combination of taxon and geography. The ‘montanus NORTH’ group includes those subspecies of Z. montanus on the islands of Luzon (Z. montanus whiteheadi) and Mindoro (Z. montanus halconensis) in the northern Philippines. The ‘montanus SOUTH’ group includes subspecies and geographic locales for Z. montanus including found in the Visayas in the central Philippines, Mindanao and adjacent islands in the southern Philippines, and Indonesia. The ‘montanus PALAWAN’ group denotes the Z. montanus parkesi subspecies on the Philippine island of Palawan. Z. japonicus is divided into the far-flung Z. japonicus alani subspecies in the Ogasawara and Volcano Islands (dark purple} and all other subspecies of Z. japonicus combined (red). Some jitter has been added to each point so as to prevent to much overlap so the reader is cautioned to not take localities as being too accurate. For a full list of samples, their source, their representation in each dataset, and more accurate locality information see supplementary table 1.

### 2.2 gDNA extraction

Whole genomic DNA (gDNA) was isolated from either blood or tissue samples using a magnetic bead based protocol (University of Kansas; Rohland & Reich 2012) or standard phenol-chloroform-isoamyl alcohol (PCI) protocol followed by ethanol precipitation (Cincinnati Museum Center and Marshall University) and quantified by a Qubit dsDNA Assay.

### 2.3 PCR and Sanger sequencing

159 individual samples underwent polymerase chain reactions (PCR) and Sanger sequencing across eight loci including two mtDNA and six nuclear protein-coding exons (nuDNA). *Z. atrifrons* was included as an outgroup. Common sequence tags were added to all the oligonucleotide sequences in this study to facilitate sequencing across all loci (Moonsamy et al., 2013). The NADH dehydrogenase subunit 2 (ND2) and cytochrome c oxidase subunit 1 (COI) were amplified using the primers L5216 and H6313 (Sorenson et al., 1999) and BIRDF1 and BIRDR2 (Hebert et al., 2004), respectively. Six nuDNA coding loci (B3GNT9, BIRC2, CCDC, GPRC6, LACTBL, and USP38) were amplified using primers from Liu *et al*. (2018). A complete list of the oligonucleotide sequences used as PCR primers in this study and the loci they amplify may be found in Supplementary table 2.

PCR and Sanger sequencing conditions are available in the supplementary material. Sequence quality control editing, assembly, and alignment were done using *Geneious* v8.1.4 (Biomatters, available from http://www.geneious.com; Kearse et al., 2012).

Chromatograms were inspected and every variation was checked for authenticity. The phase of nuclear alleles was determined statistically using *Seqphase* (Flot 2010) and *Phase* v2.1 (Stephens et al., 2001) and phased nuclear haplotypes aligned using *Geneious* (Biomatters). General descriptive statistics for Sanger sequenced loci including nucleotide diversity, haplotype diversity, segregating sites, θ, Tajima’s D (Tajima 1989), and the R_2_ test (Ramos-Onsins & Rozas 2002) as well as substitution model tests and trees and networks (distance-based and maximum likelihood) for individual loci were generated using the R packages *ape* (Paradis et al., 2004), *pegas* (Paradis 2010), and *phangorn* (Schliep 2011). Additional details and results are available in the “Supplementary: Sanger sequencing summary statistics and trees” document (https://hermmays.github.io/Zosterops1/Summary_Sanger.html).

### 2.4 StarBEAST2

A multilocus species tree was generated from the combined mtDNA and nuDNA Sanger sequencing data using *StarBEAST2* (Ogilvie et al., 2017) implemented in *BEAUTi2/BEAST2* v2.7.4 (Drummond et al., 2012; Bouckaert et al., 2014). We used *bModelTest* in *StarBEAST2* to select the substitution model for each partition/locus (Bouckaert & Drummond 2017). A strict molecular clock approach was used for each locus. The prior lognormal distribution for ND2 and COI was based on a mean of 0.0105 substitutions/site/lineage/Ma (Fleischer et al., 1998; Pereira & Baker 2006; Weir & Schluter 2008; Eo & DeWoody 2010) and a standard deviation of 1 (5–95% quantile range of 0.00123 - 0.033 and a median clock rate of 0.00637). For the intron loci we based our prior substitution rate distributions on a mean 1/10th that of the ND2 and COI rates or 0.00105 substitutions/site/lineage/Ma and a standard deviation of 1 (5–95% quantile range is 0.000123 - 0.0033 and the median clock rate is 0.000637). Three analyses in *StarBEAST2*, labeled A, B, and C, were conducted with these parameters. The taxon set for the A and B runs included information on taxon (species and subspecies) and geography. The A runs included partitions for all eight loci and the B runs include only the mtDNA loci. Another set of runs (C runs) incorporated all partitions (mtDNA and nuDNA) but was based on only taxonomic assignments (genus, species, subspecies) for each sample and no additional geographic information. 20 independent runs were conducted for each analysis with 10M length Markov chain Monte Carlo (MCMC) chains, 1M burnin lengths, and results stored every 20,000 iterations for each individual run. Additional details and results are available in the “Supplementary: StarBEAST2” document (https://hermmays.github.io/Zosterops1/Zosterops_StarBEAST2.html).

### 2.5 BPP and GDI

We tested hypotheses regarding species limits based on the eight locus Sanger sequencing dataset and the *Bayesian Phylogenetic and Phylogeography* (*BP&P* or hereafter *BPP*) program v4.7.0 (Yang 2015; Flouri et al., 2018) and estimates of the genealogical diversity index (GDI; Jackson et al., 2017; Leaché et al., 2019). BPP allows for multiple analyses including species delimitation from a fixed species tree (analysis A10 in BPP) and estimation of the population scaled mutation rate (θ) and speciation or split times (τ) from assigned species lineages on a fixed tree (analysis A00 in BPP).

Four files were created to map individual samples to taxa on a fixed species tree. The ‘tip’ schema included five putative lineages; *Z. japonicus*, *Z. meyeni*, *Z. montanus* from the Northern Philippines, *Z. montanus* from the Southern Philippines and Indonesia, and *Z. simplex*. The ‘tip’ species tree was defined such that *Z. japonicus*, *Z. meyeni*, and the northern *Z. montanus* formed a single clade sister to the southern *Z. montanus* and within that clade *Z. japonicus* and *Z. meyeni* are sister. *Z. simplex* formed an outgroup to a clade with *Z. japonicus*, *Z. meyeni*, and both lineages of *Z montanus*. Three other files were constructed such that each pair of lineages was successively collapsed starting with *Z. japonicus* and *Z. meyeni* merged together (the ‘mid1’ tree), followed by the collapse of *Z. japonicus*, *Z. meyeni*, and the northern *Z. montanus* (the ‘mid2’ tree), and finally the collapse of *Z. japonicus*, *Z. meyeni*, and both *Z. montanus* lineages into a sister group to *Z. simplex* (the ‘root’ tree). Replicate BPP runs of the A00 analysis for each of these four fixed species trees provided estimates of θ and τ along each branch which could then be used to calculate GDI for each lineage.

For the A10 runs various combinations of β_θ_ (0.001 – 0.1) and β_τ_ (0.0005 – 0.004) were used across runs corresponding to means of 0.0005 – 0.5 for θ and 0.00025 – 0.002 for τ prior distributions. These values were chosen based on what would be reasonable for a relatively recent set of lineages (< 1mya) and in consultation with the BPP manual (https://bpp.github.io/bpp-manual/bpp-4-manual/). For all the A10 and A00 runs the general time reversible (GTR) substitution model was used with a burnin of 10,000 and a sample size of 100,000. Species delimitation algorithms 1 and 0 were tested in the A10 runs.

Three replicate runs were done for the A00 analyses for each of four combinations of prior parameters (β_θ_ and β_τ_) across four fixed species trees (‘tip’, ‘mid1’, ‘mid2, and ‘root’) making a total of 48 A00 runs. Results from each set of three replicate runs under the same parameters were combined and GDI was calculated from estimates of θ and τ. θ and GDI were plotted in R as density plots for each branch on the fixed species tree. A full list of run parameters for the A10 runs may be found in Supplementary table 3 and parameters for each A00 run may be found in Supplementary table 4. BPP control files are included with the Supplementary data.

This analysis was confined to those taxa with the greatest taxonomic uncertainty (*Z. japonicus*, *Z. meyeni*, and *Z. montanus*, with *Z. simplex* as an outgroup). Additional details and results are available in the “Supplementary: BPP and GDI based species delimitation” document (https://hermmays.github.io/Zosterops1/ZosteropsBPP.html).

### 2.6 RADseq-derived SNP dataset

The RADseq-derived SNP dataset in this study is described in DeRaad *et al*. (2024). Briefly, restriction site associated DNA libraries were constructed by digesting gDNA using the enzyme *NdeI* and performing size selection for fragments between 495 – 605 base-pairs (bp). The complete library preparation protocol is available at https://github.com/DevonDeRaad/zosterops.rad/blob/main/lab.protocols/MSG-NdeI_2plates-150samples_DAD-Moyle_230530.doc. Pooled, bar-coded libraries were sequenced on the Illumina NextSeq2000 at the University of Kansas Genomic Sequencing Core using a P2 flow cell to generate 414,215,817 single-end 100 bp reads (β 41Gb raw sequence data). Raw genomic sequencing data is publicly available via the SRA BioProject PRJNA1079333 (https://www.ncbi.nlm.nih.gov/sra/PRJNA1079333).

### 2.7 Read mapping, variant calls, and filtering

Mapping, variant calling, and filtering were performed as in DeRaad *et al*. (2023) and DeRaad *et al*. (2024). Raw data was demultiplexed and reads of low quality and with uncalled bases were removed using *Stacks* v2.41 (Rochette et al., 2019). Reads were then mapped to a publicly available *Z. japonicus* genome assembly available at https://www.ncbi.nlm.nih.gov/assembly/GCA_017612475.1 (Venkatraman et al., 2021) using the *Burrows-Wheeler Alignment tool* (*BWA*) v0.7.17 (Li & Durbin 2009). *SAMtools* v1.3.1 (Li et al., 2009) was used to convert files to a sorted bam format and RAD loci were identified in *Stacks* using the ‘gstacks’ module and exported as a variant call format (vcf) file resulting in an unfiltered SNP dataset containing 236,767 SNPs shared among 155 individual samples with 65.5% missing genotypes. The R packages *vcfR* v1.14.0 (Knaus & Grünwald 2017) and *SNPfiltR* v1.0.1 (DeRaad 2022) were used to visualize parameter distributions and filter the SNP data for quality. The final filtered dataset included 15,704 SNPs across 124 individual samples with 5.3% missing data. A more detailed description of the filtering protocol may be found in DeRaad et al. (2024) and at: https://devonderaad.github.io/zosterops.rad/zost.radseq.filtering.html.

A pairwise distance matrix based on Nei’s D (1972) was generated from the filtered SNP dataset using the R package *StAMPP* v1.6.3 (Pembleton et al., 2013) and used to construct a neighbor-net phylogenetic network using *SplitsTree4* v4.15.1 (Huson & Bryant 2006). This allowed us to visualize the data and, in some cases, validate the identity of individual samples for subsequent analyses. For example, the sample designated Zsim30897 was initially identified as *Z. simplex* however in the phylogenetic network described above this sample clustered with *Z. palpebrosus* (Supplementary table 1). Associated R code and the phylogenetic network from *SplitsTree* v4.15.1 are described in DeRaad et al. (2024) and may be viewed at https://devonderaad.github.io/zosterops.rad/splitstree.html.

### 2.8 General SNP visualization and multivariate analyses

The filtered SNP data was converted to a genlight object in the R package *vcfR* (Knaus & Grünwald 2017). The R package *Poppr* (Kamvar et al., 2014) was used to generate a minimum spanning network (MSN) and a distance tree based on the unweighted pair group method (UPGMA). Bootstrapping was performed for the UPGMA tree with 100 replicates.

A principal component analysis (PCA) and a discriminant function analysis of principal components (DAPC) was conducted using the R package *adegenet* (Jombart 2008) following the methodology from Jombart *et al*. (2010). DAPC takes principal components (PCs) and partitions variance into within- and among-group components to maximize discrimination among the groups and makes no assumptions regarding recombination or panmixia, unlike *Structure* (Pritchard et al., 2000; Porras-Hurtado et al., 2013) or *Admixture* (Alexander et al., 2009).

Samples were grouped by a combination of species, subspecies, and geography with *Z. japonicus* split into the most far-flung *Z. japonicus alani* subspecies in the Ogasawara and Volcano Islands and a broader *Z. japonicus* group encompassing the remaining subspecies. *Z. montanus* samples were grouped into a Northern Philippine group including *Z. montanus whiteheadi* on Luzon and *Z. montanus halconensis* on Mindoro, *Z. montanus parkesi* on Palawan, and a southern group including the remaining *Z. montanus* subspecies. The remaining samples were assigned to groups based on their species.

PCA was conducted on the filtered SNP dataset (15,704 SNPs, 124 individual samples) using the sample groups described above. We generated plots in two dimensions (PCs 1-2) and in three dimensions (PCs 1-3) using the R package *plotly* (Sievert 2020).

We conducted a cross-validation analysis to help determine the optimal number of PCs to be retained in the DAPC by examining the proportion of successful assignments for each PC for the first 50 PCs and 100 replicate runs per PC (Supplementary figure 2). The median optimal number of PCs was calculated from these 100 runs and used in the subsequent DAPC analysis to estimate the posterior membership probability for each individual sample. We conducted two additional DAPC analyses based on 50% fewer and 50% greater than the optimal number of PCs.

In addition to PCA we conducted a second dimensionality reduction on the SNP data using the uniform manifold approximation and projection (UMAP) method (Healy & McInnes 2024). Analysis via both methods allowed for the visualization of the SNP data by a linear method emphasizing variation (PCA) and a non-linear method emphasizing global structure among groups (UMAP). UMAP was performed with the R package *umap* and plots generated in two and three dimensions using the R package *plotly* (Sievert 2020). Additional details for these analyses and results are available in the “Supplementary: MSN, Distance trees, PCA, UMAP, and DAPC analyses” document (https://hermmays.github.io/Zosterops1/ZosteropsSNPs_analysis.html).

### 2.9 Fixed differences, F_ST_, and Nei’s D

Pairwise fixed differences were measured from the SNP dataset in R using the *dartR* v2 package (Mijangos et al., 2022). Samples were grouped according to a combination of species and geography with *Z. japonicus* split into an Ogasawara and Volcano Islands group containing *Z. japonicus alani* and a ‘continental’ island group containing the remaining subspecies. *Z. montanus* was divided into four groups: the nominate subspecies in Indonesia, a northern Philippines group, a central/southern Philippines group, and the *Z. montanus parkesi* subspecies on Palawan. In every tree produced, and the PCA and DAPC analyses, our single sample from *Z. nigrorum catarmanensis* from the island of Camiguin South (KU110362) fell within *Z. montanus* with consistent and strong support and was considered a central/southern Philippine *Z. montanus* for these analyses. The remaining samples were assigned according to species.

Pairwise F_ST_ and pairwise Nei’s distances were measured using *StAMPP*. Additional population grouping was done for these analyses with *Z. japonicus* divided into populations in the northern, central, and southern Ryukyu Islands, the Korean Peninsula and adjacent islands, the main islands of Japan, the Ogasawara and Volcano Islands, the Izu Islands, and the introduced population in Hawaii. *Z. montanus* was divided into the Indonesian, northern Philippine, Visayas, Mindanao, and Palawan populations. *Z. simplex* was divided into those samples from the native range in Asia and introduced populations in North America. The rest of the samples were grouped only by species. Additional details for these analyses and results are available in the “Supplementary: Genetic differentiation and biogeographic breaks” document (https://hermmays.github.io/Zosterops1/Zosterops_SNPFst.html).

### 2.10 Species tree and delimitation

A species tree was produced from the SNP dataset using Singular Value Decomposition Quartets (SVDQuartets; Chifman & Kubatko 2014) implemented in PAUP* v4.0 (Swofford 2003; Wilgenbusch & Swofford 2003) and Phylogenetic Inference with Composite Likelihood (PICL; Swofford & Kubatko 2023; Kong et al., 2025; Kubatko et al., 2025). The SNP file was converted to nexus format using *vcf2phylip* (Ortiz 2019). A taxon file was created to assign individual samples to narrowly defined groups assorted primarily by species, subspecies, and island of origin. Samples from *Z. palpebrosus* were defined as the outgroup taxon. SVDQuartets was run in PAUP* using this taxon file and the nexus formatted SNP data by evaluating all quartets and 700 bootstrap replicates. A 50% majority rule consensus tree was then derived from the output trees using the ‘contree’ command in PAUP*, saved, and visualized in *FigTree* (Rambaut 2014).

We conducted two estimates of branch lengths using PICL based on the tree topology determined from the SVDQuartets. Taxon designations for individual samples were assigned by a combination of taxonomy and geography, similar to prior analyses. For the first run *Z. japonicus* was split into *Z. japonicus alani* from the Ogasawara and Volcano Islands and a *Z. japonicus* group with the remaining subspecies, a northern Philippine group of *Z. montanus* that included the subspecies from Luzon and Mindoro, and another grouping the *Z. montanus parkesi* subspecies from Palawan. The remainder of the *Z. montanus* populations were split into *Z. montanus* from the Visayas, Camiguin South, and Mindanao and another group containing nominate *Z. montanus* subspecies samples from Sulawesi and Java. As in prior analyses *Z.nigrorum catarmanensis* (KU110362) on Camiguin South was grouped with the central/southern Philippine *Z. montanus* samples. The remainder of the samples were grouped by species.

For the second run taxon assignments were based on any population lineages from the SVDQuartets analysis with bootstrap values greater than 60%. Poorly supported lineages (<60%) were collapsed, with two exceptions. *Z. japonicus stejnegeri* on Kozushima and Miyakejima in the Izu Islands exhibited bootstrap support greater than 60% but because these sister populations were deeply embedded in a group of very poorly supported lineages, all continental island populations of *Z. japonicus*, along with the Izu Island populations of *Z. japonicus stejnegeri*, were collapsed into a single ‘continental’ *Z. japonicus* clade. Also, despite the mismatched *Z. nigrorum catarmanensis* sample resting within a poorly supported group along with the *Z. montanus* populations in the Visayas, we treated this sample as its own group to explore the taxonomic mismatch. For both runs the parameters were the same except for the taxon/population assignments for the individual samples to determine branch lengths on more collapsed (run 1) and less collapsed (run 2) versions of the SVDQuartets consensus tree. 100 bootstrap replicates were performed to obtain 95% confidence intervals associated with each split time.

We took the resulting species tree with the topology determined by SVDQuartets and branch lengths by run 2 of PICL and used this to test species hypotheses in *Delineate* (Sukumaran et al., 2021). We conducted five independent runs of *Delineate* each with a different constraint schema. Because of a comparatively deep split between *Z. nigrorum nigrorum* on Panay and the other *Z. nigrorum* subspecies in this study, this lineage was left unconstrained in all five analyses. All of the lineages in the recently diverged *Z. japonicus*/*meyeni*/*montanus* clade, including the mismatched *Z. nigrorum catarmanesnis*, were left unconstrained in run 1. In run 2 only the southern Philippine and Indonesian *Z. montanus* within *Z. japonicus*/*meyeni*/*montanus* clade were constrained as a species lineage. In run 3 both the southern *Z. montanus* and *Z. meyeni* lineages were constrained. In run 4 only *Z. meyeni* was constrained. In run 5 only *Z. japonicus* was constrained (both the ‘continental’ *Z. japonicus* and *Z. japonicus alani*). For each of these runs the ‘estimate partitions’ command was run in *Delineate* with a nexus formatted version of the consensus tree from SVQuartets and PICL with branch lengths in coalescent units (2N_e_ generations) and a file designating the lineage assignments as either constrained (1) or unconstrained (0). For each run only those partitions contributing to 95% of the cumulative probability were reported. Additional details for these analyses and results are available in the “Supplementary: Species trees and delimitation with SVDQuartets, PICL, and DELINEATE” document (https://hermmays.github.io/Zosterops1/Zosterops_SVDQ.html).

### 2.11 Morphometric analyses

Linear morphometric data including culmen length, nares-to-tip bill length, bill depth, bill width, tarsus length, wing cord, and tail length were analyzed from 354 round museum skins from the American Museum of Natural History (New York, NY, USA), Cincinnati Museum Center (Cincinnati, OH, USA), The Field Museum (Chicago, IL, USA), and the Smithsonian National Museum of Natural History (Washington DC, USA). All measurements were recorded by the same researcher (HLM) in mm using either digital calipers or a wing rule. Culmen length was measured as exposed culmen from the tip of the bill to the point where the bill meets the forehead plumage. Bill length from nares-to-tip was measured from the distal end of the right nare to the tip of the bill. Bill depth and width were measured at the nares. Tarsus (tarsometatarsus) length was measured from the base of the phalanges to the tibiotarsal joint. Wing cord was measured unflattened with a wing rule. Tail length was measured from the body to the tip of the longest rectrix by placing the same wing rule in the center of the tail.

A PCA was conducted and PC1-2 and PC1-3 were each plotted in R with specimens grouped solely according to species. Specimens were further divided into 11 groups for pairwise statistics including a ‘continental’ group of *Z. japonicus* specimens from the main islands of Japan, the Ryukyu Archipelago, and the Korean Peninsula and surrounding islands, an ‘oceanic’ *Z. japonicus* group including those subspecies *stejnegeri* and *alani* on oceanic islands in the Pacific (Izu, Ogasawara, and Volcano Islands), two *Z. montanus* groups including a northern Philippine group on the islands of Luzon and Mindoro, and a southern *Z. montanus* group in the central and southern Philippines and Indonesia, and the *Z. nigrorum catarmanensis* subspecies on the Philippine island of Camiguin South. The remaining samples, *Z. erythropleurus*, *Z. everetti*, *Z. meyeni*, *Z. nigrorum* (excluding *catarmanesnis*), *Z. palpebrosus*, and *Z. simplex*, were each grouped according to species. Box-and-whisker plots were generated for both grouping schemas, solely by species and by a combination of geography and species, tests for normality were conducted, and pairwise Wilcoxon tests were conducted for each pair of groups.

A complete list of specimens in this analysis is available in Supplementary table 5. Additional details for these analyses and results are available in the “Supplementary: Linear morphometrics” document (https://hermmays.github.io/Zosterops1/ZosteropsMorphometrics.html).

## 3. Results

### 3.1 Sanger sequenced loci

159 individual samples were sequenced for at least one of eight Sanger sequenced loci. We produced sequence data for 132 individual samples for COI and 143 samples for ND2. Across the six nuclear nuDNA loci we sequenced an average of 88 individual samples per locus (87 – 91 samples) and 5,685 bp across all eight loci including 646 bp and 978 bp for COI and ND2, respectively, and an average of 676.8 bp per nuDNA locus (559 – 727 bp). There were 119 and 235 segregating sites for COI and ND2, respectively, and an average of 24.7 segregating sites per nuDNA locus. Basic population genetic statistics, distance networks, and distance-based and maximum likelihood trees for each locus are summarized in Supplementary table 6 and detailed in the “Supplementary: Sanger sequencing summary statistics and trees” document (https://hermmays.github.io/Zosterops1/Summary_Sanger.html).

### 3.2 StarBEAST2

The estimated samples sizes (ESS) computed from the combined log files from the StarBEAST2 runs from nuDNA and mtDNA varied but generally indicated poor MCMC convergence. However, statistics from the combined log files from those runs with only mtDNA partitions showed no ESS for any parameter less than 200 with most well over 1,000 indicating good convergence of the MCMC.

Both the StarBEAST2 species tree employing all eight loci as partitions (Figure 2a) and the tree restricted to only mtDNA partitions (Figure 2b) showed poor resolution. Species tree resolution was not improved with the addition of the nuDNA partitions. The tree restricted to the mtDNA partitions tree exhibited more highly-supported nodes (0.9 – 1.0 posterior probability). Both species trees supported a monophyletic *Z. japonicus*, albeit with better posterior support in the mtDNA partitioned analysis (1.0) than that with all loci (0.6). Posterior support for a *Z. japonicus*, *Z. meyeni*, and *Z. montanus* clade was 0.86 and 0.99 for the combined runs and the mtDNA only runs, respectively. However, this clade also unexpectedly included samples from *Z. everetti* and the *Z. nigrorum catarmanensis* sample from Camiguin South. *Z. erythropleurus*, *Z. simplex*, and *Z. nigrorum* (minus the *catarmanensis* subspecies) each formed well supported monophyletic clades in both analyses (Figure 2).

**Figure 2:**
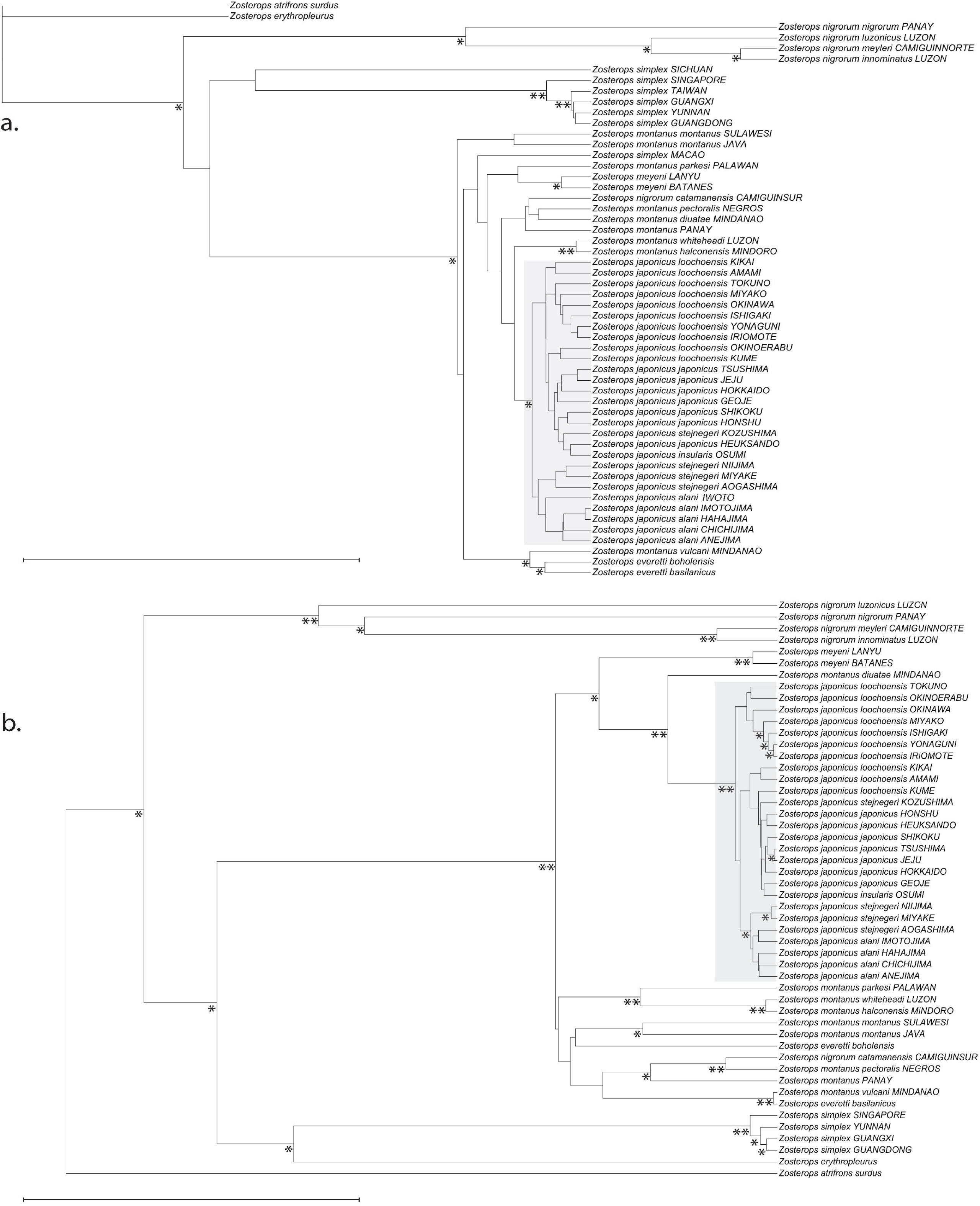
Species trees generated from 8 loci (2 mtDNA, 6 cdDNA) in StarBEAST2 using a taxon set based on taxonomy and geography and unlinked partitions for all 8 loci (a) and a reduced partitioning including only the mtDNA loci, COI and ND2 (b). Posterior probability support is shown where support is greater than 0.5. Support 0.5 - 0.9 is denoted by a single asterisk(*) at the bottom left of the corresponding node. Support greater than 0.9 is indicated by a double asterisk(**) in the same orientation. Z. atrifrons surdus is a designated outgroup in both trees. In each tree a monophyletic Z. japonicus clade is highlighted in grey. The scale bar for each tree represents approximately 2 million years.

Based on the clock rates provided in substitutions/site/lineage/Ma we were able to generate estimates of node ages within the Asiatic white-eyes but only within broad 95% highest posterior densities (HPD). For the A runs with nuDNA and mtDNA we found the root of the tree to be at 3.09 Ma (0.26 – 5.9 Ma 95% HPD) before the present (BP) and the *Z. japonicus* clade at 0.36 Ma (0.07 – 0.49 Ma 95% HPD) BP. The B runs based only on mtDNA the tree root was at a median time of 3.65 Ma (0.3 – 9.44 Ma 95% HPD) BP and the *Z. japonicus* clade dated to 0.24 Ma (0.02 – 0.65 Ma 95% HPD) BP. The C runs with taxon assignments limited to subspecies, rather than a combination of taxonomy and geography, showed an age for the root of 3.63 Ma (1.21 – 6.73 Ma 95% HPD) and 0.19 Ma (0.05 – 0.37 Ma 95% HPD) for *Z. japonicus*.

### 3.3 BPP and GDI

The BPP A10 analysis delimiting species from a fixed tree identified all five lineages as species regardless of the θ and τ priors or the species delimitation algorithm (1 or 0, Supplementary table 3). The GDI density distributions were consistent across all combinations of prior distributions for θ and τ. Density distributions for GDI measured from the *Z. meyeni* and *Z. simplex* branches fell within the range for a putative but ambiguous species (0.2 – 0.7) and in the case of *Z. simplex* exceeded the value suggested as a rule-of thumb to denote a distinct species (0.7; Jackson et al., 2017; Leaché et al., 2019). The GDI distribution for the *Z. montanus* lineage from the northern Philippines fell mostly within the range where species delimitation is ambiguous. The other distributions were either mostly or entirely less than 0.2, offering no support for independent species lineages in these groups (Figure 3).

**Figure 3:**
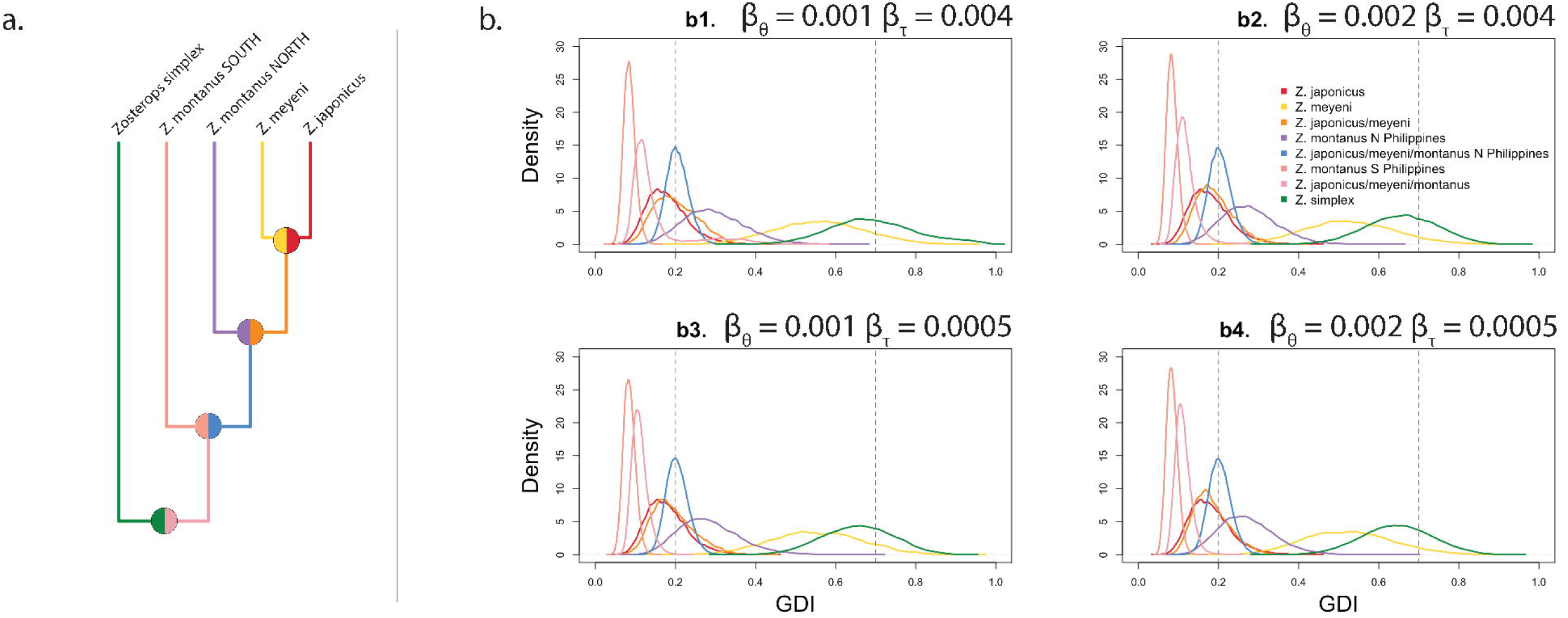
Genealogical diversity indices (GDI) density distributions as estimated from the Bayesian Phylogenetics and Phylogeography (BPP} program across four different combinations of priors for θ and τ using the multilocus Sanger sequencing dataset. α and β are parameters that define an inverse Γ distribution for the θ and τ priors. For all runs α = 3. Combinations of β for θ and τ prior distributions are shown above each set of GDI results (b). A putative species tree used in all BPP analyses is shown on the left (a) with color coded branches corresponding to GDI density distributions on the right (b).12 replicate runs were executed for each successively collapsed tree including the full species tree (a), a tree with Z. meyeni and Z. japonicus collapsed, a tree with Z. montanus, Z meyeni, and Z. japonicus collapsed, and a tree with both north and south Z. montanus collapsed along with Z. meyeni and Z. japonicus. Estimating θ and τ on successive collapses of a fixed species tree allows for the calculation of GDI for each branch of the tree. Dashed lines on the density plots for each set of estimates corresponds to a GDI of 0.2 and 0.7. These values have been used in previous studies in hueristic species delimitation with a GDI < 0.2 corresponding to population lineages within the same species, 0.2 < GDI < 0.7 corresponding to ambiguous species status, and 0.7 < GDI corresponding to strongly delimited species lineages (Jackson et al. 2017; Leache et al. 2019).

### 3.4 General SNP variation, PCA, UMAP, and DAPC

The final filtered SNP dataset included 15,704 SNPs across 124 individual samples with 5.3% missing data. Clustering of individual samples by distance is shown in an MSN (Supplementary Figure 3) and a UPGMA distance tree (Supplementary figure 4). The UPGMA tree reveals a well-supported clade with all *Z. japonicus*, *Z. meyeni*, and *Z. montanus* samples. Within this group is a strongly supported monophyletic *Z. japonicus*. In both the MSN and UPGMA tree is a mismatched *Z. nigrorum catarmanensis* sample (KU110362) clustering strongly with the *Z. montanus* samples from the southern Philippines.

Pairwise comparisons of fixed differences in the SNP dataset ranged from 0.0% (0/15,704) for the comparison between *Z. japonicus alani* from the Ogasawara and Volcano Islands and the remainder of the *Z. japonicus* populations to 10.5% (1,570/15,000) for the *Z. montanus* samples from Indonesia and *Z. palpebrosus* (Supplementary table 7). Pairwise Nei’s distances ranged from 0.002 between the Korean and main Japanese island populations of *Z. japonicus* and 0.15 between *Z. palpebrosus* and the introduced *Z. simplex* population in North America. For pairwise F_ST_ the Korean and main Japanese Island populations of Z. japonicus were the most similar (0.02) and *Z. palpebrosus* pairwise F_ST_ comparisons with the Korean, central Ryukyu, Izu, and Ogasawara populations of *Z. japonicus* were the most different (0.87, Supplementary figure 5).

The first three PCs explain 53.6% of the genetic variance in the SNP data (PC1 29.6%, PC2 16.3, PC3 7.7%). Distinct, non-overlapping clusters for a plot of PC1-2 are exhibited for *Z. simplex*, *Z. erythropleurus*, and *Z. palpebrosus*. Overlap among individual samples is evident within a cluster of *Z. everetti* and *Z. nigrorum* and within another cluster containing the samples from *Z. japonicus*, *Z. meyeni*, and *Z. montanus*. As consistently seen in other analyses, the *Z. nigrorum catarmanensis* sample (KU110362) clusters with the *Z. japonicus*/*meyeni*/*montaus* group (Figure 4). Adding a third axis of variation by plotting three PCs increases the separation between *Z. japonicus* samples and *Z. meyeni* and *Z. montanus* (Supplementary figure 6).

**Figure 4:**
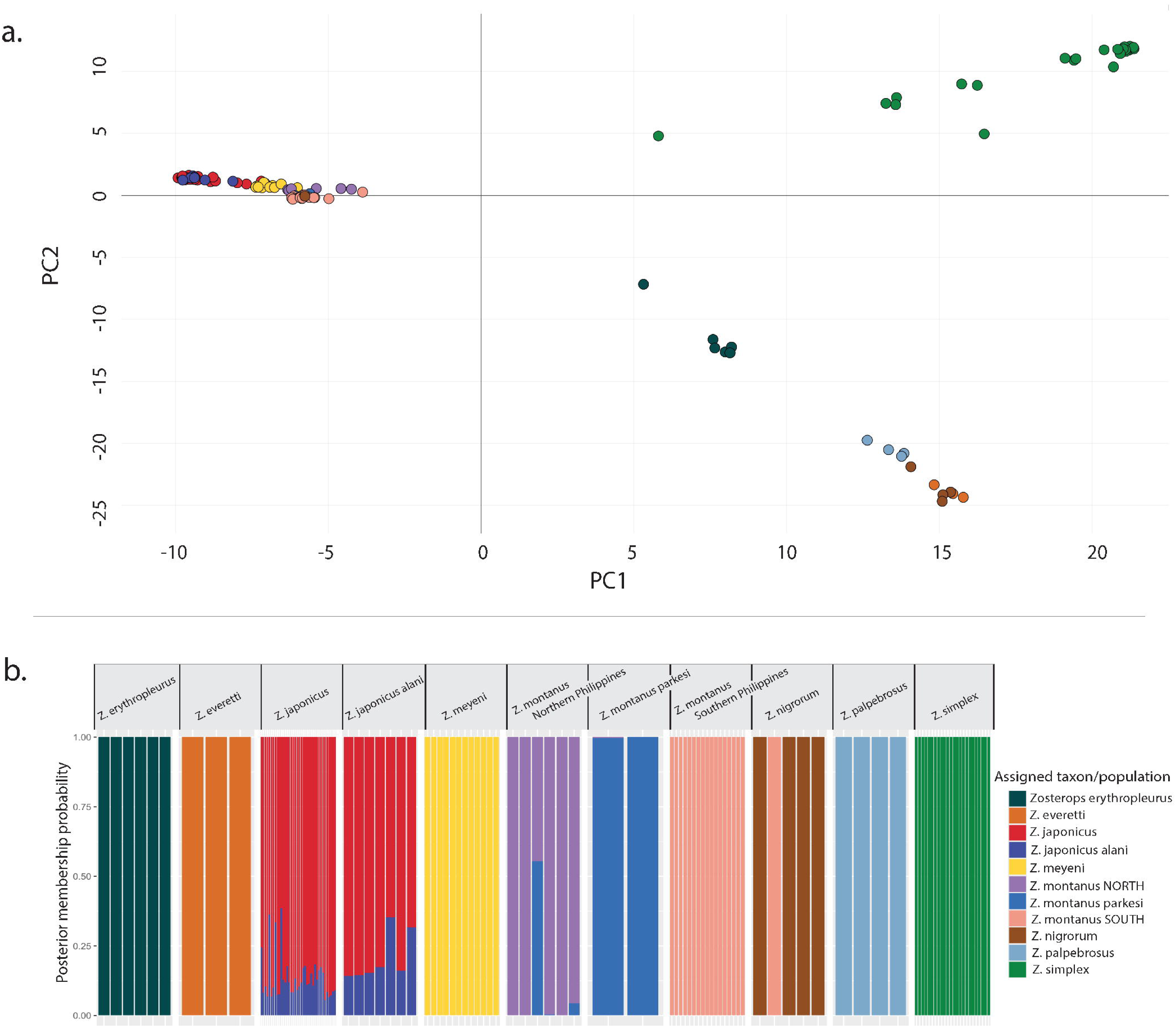
Plot of principle components 1 and 2 from a principle component analysis (PCA) of the Zosterops SNP dataset(a). Discriminant analysis of principle components (DAPC) posterior membership probabilities for individual samples within each taxonomic/geographic group based on the Zosterops SNP dataset (b). Cross validation was performed to determine the optimal number of principle components for discriminant analysis. The optimal number of principle components maximizes the percent improvement in discrimination among groups over that based on random assignments. For the DAPC in this figure the optimal number of principle components used was eight. DAPC results based on four and 12 principle components are included in the supplementary data. Note that one individual sample assigned to Zosterops nigrorum has a 100% posterior probability of membership with the Z. montanus populations in the Southern Philippines. This is the same individual sample that is also mismatched in other analyses. Color coding is the same for figures a. and b.

UMAP analysis of the SNP data revealed better discrimination among groups than PCA including a clear clustering of *Z. japonicus alani* from the rest of *Z. japonicus*. The mismatched *Z. nigrorum catarmanensis* sample (KU110362) was also evident on the UMAP plot (Supplementary figure 7).

We determined that the median optimal number of PCs retained for discrimination was eight, based on 100 replicate a-score estimates. Ancillary DAPC was also conducted for four (50% less) and 12 (50% greater) PCs. For DAPC with four PCs, discrimination was comparatively poor with unanimity only for *Z. erythropleurus*, *Z. palpebrosus*, and *Z. simplex*. DAPC could not effectively discriminate between *Z. japonicus* and *Z. japonicus alani* based on any of the DAPC analyses (4, 8, or 12 PCs). However, for eight PCs, DAPC unanimously discriminated samples into the correct group for all other groups outside of *Z. japonicus* and *Z. japonicus alani* except for *Z. montanus* in the northern Philippines (showing some ambiguity with *Z. montanus parkesi* from Palawan), and *Z. nigrorum*. For 12 PCs the DAPC unanimously discriminated samples for all the groups outside of *Z. japonicus* and *Z. japonicus alani* except for *Z. nigrorum*. In the case of *Z. nigrorum* the same *Z. nigrorum catarmanensis* sample mismatched in other analyses (KU110362) was identified in all DAPC analyses as belonging to the *Z. montanus* group in the southern Philippines (Figure 4b and Supplementary figure 8).

### 3.5 SVDQuartets and PICL species trees

The species tree topology from SVDQuartets was broadly consistent with other studies and other analyses in this study (Supplementary figure 9). Specifically, *Z. japonicus*, *Z. meyeni*, and *Z. montanus* form a single well-supported clade encompassing a paraphyletic *Z. montanus* and monophyletic *Z. meyeni* and *Z. japonicus*. Paraphyly in *Z. montanus* followed a pattern consistent with other studies (Jones & Kennedy 2008; Lim et al., 2019) where the northern subspecies of *Z. montanus* were closely allied with *Z. japonicus*/*meyeni*. Within *Z. japonicus* the only well supported split among subspecies was between *Z. japonicus alani* in the Ogasawara and Volcano Islands and the rest of *Z. japonicus* (96% bootstrap support). The other species each formed their own well-supported clades with one notable exception. The *Z. nigrorum catarmanensis* sample (KU110362) was consistently across all analyses embedded within the central and southern populations of *Z. montanus*.

Independent runs of PICL based on two schemas for collapsing taxa from the SVDQuartets topology produced broadly similar results in terms of split times (Figure 5 and Supplementary figure 10). SVDQuartets identified strongly supported nodes that in PICL were associated with small split times, in some cases indistinguishable from zero. The split between *Z. japonicus alani* in the Ogasawara and Volcano islands and the rest of *Z. japonicus*, for example, was well supported in SVDQuartets (96% bootstrap support) but was associated with a speciation/split time indistinguishable from zero. The same was true for the split between *Z. japonicus* and *Z. meyeni*, the splits within the southern Philippine and Indonesian lineages of *Z. montanus*, and the split between the southern *Z. montanus* and *Z. montanus parkesi* on Palawan.

**Figure 5:**
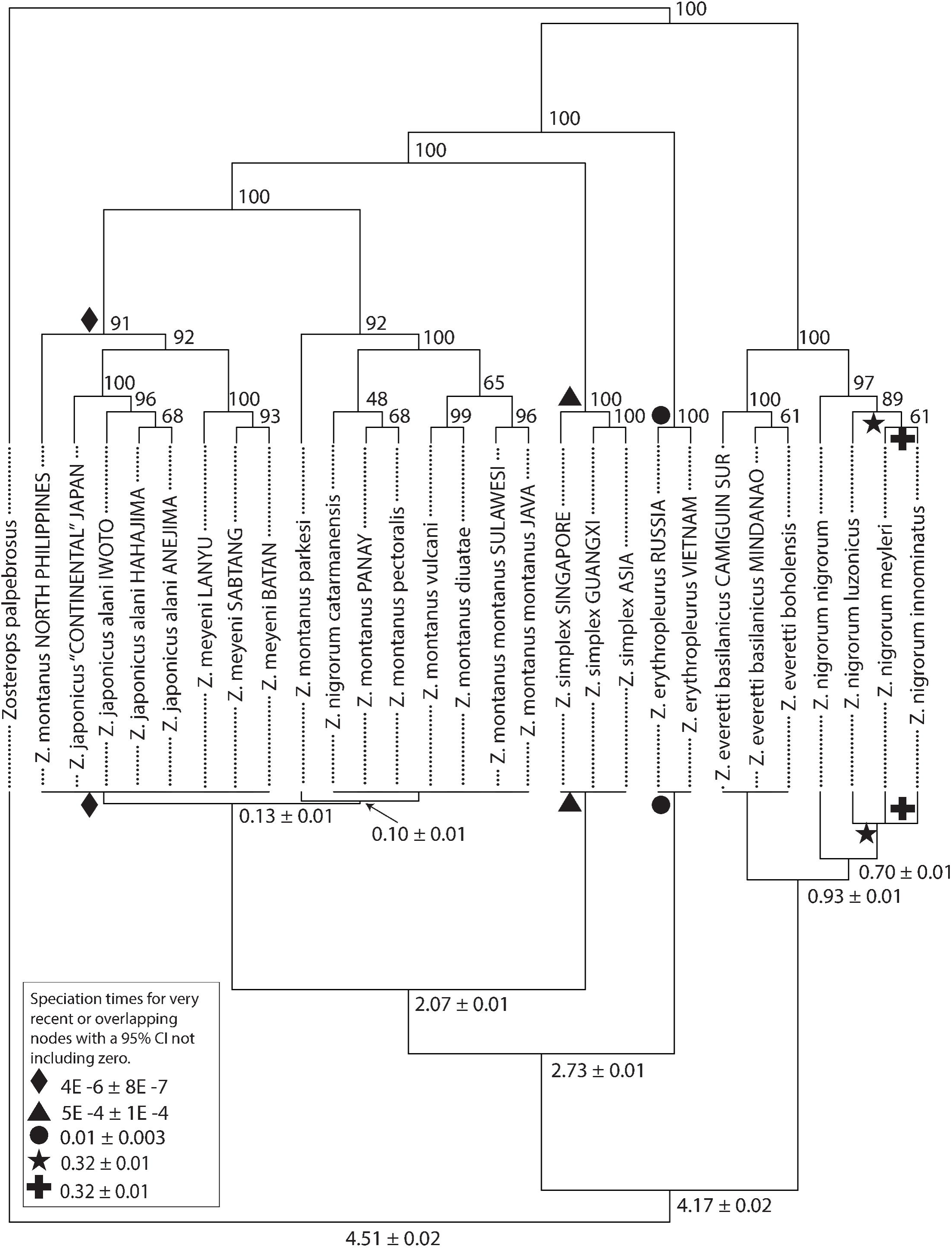
The top tree is the collapsed topology from the SVDQuartets analysis for PICL run 2 for Zosterops SNP data with percent support values at each node based on 700 bootstrap replicates. The bottom tree is the result from PICL run 2 with split times and 95% confidence intervals (Cls) labeled at each node when the time for the node was significantly different from zero. Nodes with very small or overlapping split times were difficult to label and are marked with symbols on both trees accompanied by a key showing the split time and 95% Cls for that marked node. Confidence intervals for split times in PICL were estimated from 100 bootstrap replicates. Any nodes with 95% Cls for split times overlapping with zero are unlabeled. Note that for some recent nodes there may be high bootstrap support from SVDQuartets but with a 95% Cl overlapping with zero. The ‘Z. simplex ASIA’ group includes samples from Guangdong, Guizhou, and Hunan provinces in Mainland China, Taiwan, Vietnam, and the introduced population of Z. simplex in Southern California, USA. The ‘Z. japonicus “CONTINENTAL’’JAPAN’ includes those populations of Z. japonicus on the main islands of Japan, the Korean Pennisula and adjacent islands, the Ryukyu archipelago, the lzu Islands, and the introduced population of Z. japonicus in Hawaii, USA.

### 3.6 Delineate

We conducted five independent runs of *Delineate* (Sukumaran et al., 2021) based on different constraints imposed on the tree generated in SVDQuartets and PICL. Results were broadly congruent with other studies in that there was strong support for a single species lineage for the clade containing *Z. japonicus*, *Z. meyeni*, and *Z. montanus* across four of the five runs. A single species consisting of *Z. japonicus*, *Z. meyeni*, and the northern Philippine populations of *Z. montanus* was delimited from *Z. montanus* populations in the central and southern Philippines, Indonesia, and Palawan, but this delimitation scenario was only supported in the analysis when existing constraints were introduced within each group (run 3). When there were only constraints for a single species within one or the other of these groups (*Z. japonicus*/*Z. meyeni*/north *Z. montanus* versus south and Palawan *Z. montanus*, runs 2, 4, and 5), or when there were no *a priori* constraints on this clade (run 1), *Delineate* best supported combining both groups within a single species lineage. Under the least constrained scenario there were 14 species partitions within the 95% cumulative probability out of 53,822,103 total partitions. The best supported three partitions lumped *Z. japonicus*, *Z. meyeni*, and *Z. montanus* into a single species in run 1. The fourth and fifth most likely partitions in this least constrained run split *Z. japonicus*/*meyeni*/north *montanus* into a species distinct from the rest of *Z. montanus* with an unconstrained probability of 0.013 and 0.010, respectively. Among the 14 partitions in run 1 within the 95% cumulative probability a *Z. japonicus*/*meyeni*/north *montanus* species was split from the southern *Z. montanus* in six of those partitions. The nominate *Z. nigrorum* population on Panay was strongly supported as a separate species lineage from the remaining *Z. nigrorum* subspecies, all on Luzon and adjacent islands, under all five constraint scenarios. The best species tree across all analyses supported the lumping of *Z. japonicus*, *Z. meyeni*, and *Z. montanus* and including the *Z. nigrorum catarmanensis* within a single clade, as well as a north-south species split between the more southerly nominate subspecies of *Z. nigrorum* on the island of Panay and the remaining subspecies on and adjacent to Luzon (Figure 6).

**Figure 6:**
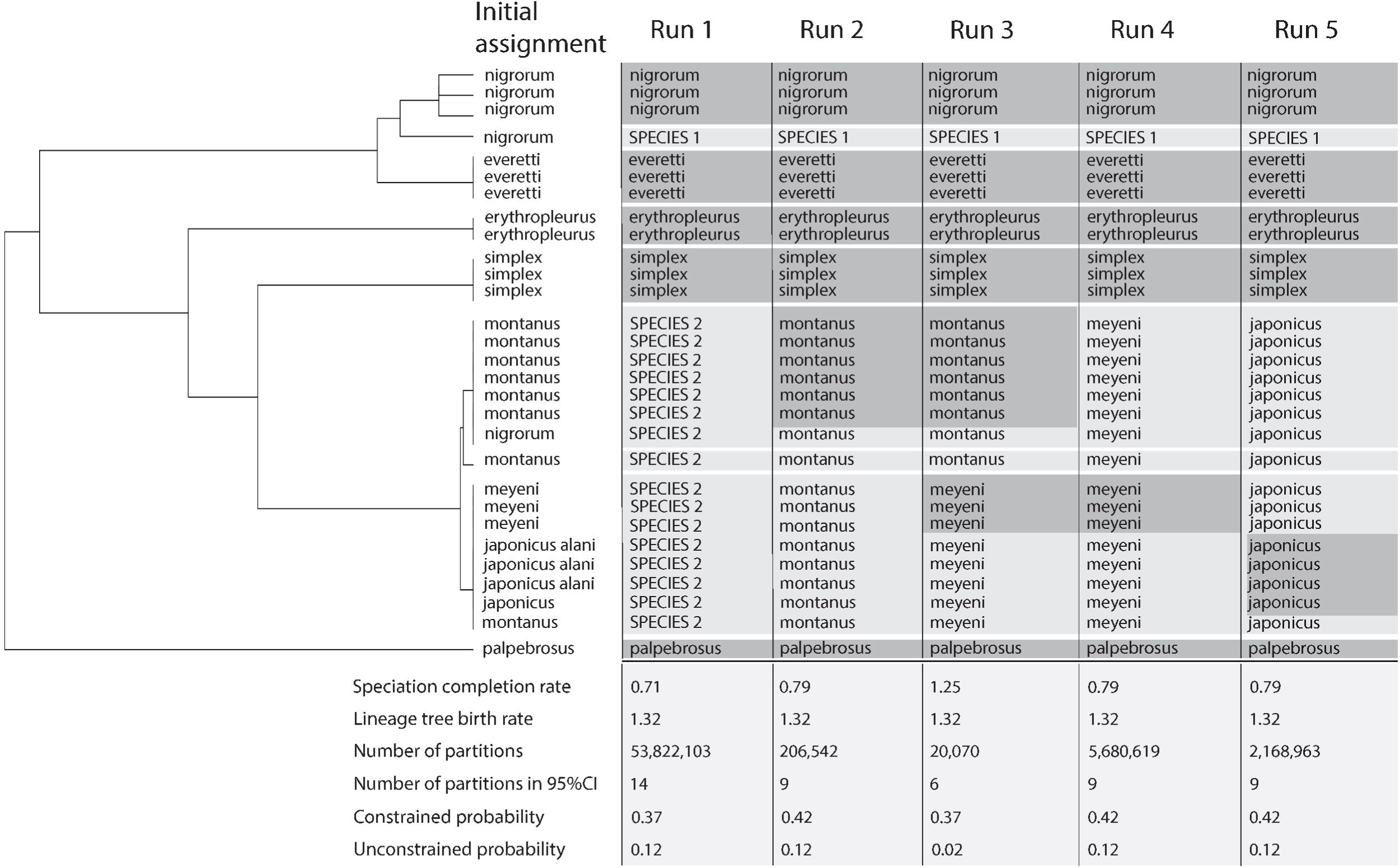
Results from Delineate analysis for the Zosterops SNP dataset. The tree used for the analysis is shown on the left with the initial taxon assignments shown for each lineage. Columns on the right of the tree show the most probable results for each of five runs of Delineate, each with different lineage constraints. Constrained lineages in each run are highlighted in dark grey and the unconstrained lineages are shown in light grey. The ‘SPECIES’ designation shows where a new species has been delineated in that run. The quantitative results of each run are shown at the bottom of each column. Run 1 had the fewest constraints with all Z.japonicus, Z.japonicus alani, Z. meyeni, Z. montanus, Z. nigrorum nigrorum (in the Z. nigrorum clade), and Z. nigrorum catarmanensis (in the Z. japonicus/meyeni clade) left unconstrained. Runs 2 - 4 included constrained lineages in either the Z. montanus clade (run 2), the Z. japonicus/meyeni clade (runs 4, 5), or constrained lineages in both clades (run 3). Z. nigrorum nigrorum in the Z. nigrorum clade was left unconstrained throughout all five runs. Run 1 delimited Z. nigrorum nigrorum as a new species (SPECIES 1) and all the lineages in the Z.japonicus/meyeni/montanus clade as a single species (SPECIES 2). lfthe southern Philippine Z. montanus lineages were contrained then all lineages in the Z. japonicus/meyeni/montanus clade were delimited as Z. montanus, including the mismatched Z. nigrorum catarmanensis lineage in the Z. montanus clade. If the Z. meyeni lineages were constrained then all lineages in the Z. japonicus/meyeni/montanus clade were delimited as Z. meyeni, again including the mismatched Z. nigrorum catarmanensis lineage. If both the southern Philippine Z. montanus lineages and the Z. meyeni lineages were constrained (run 3) then the Z. japonicus/meyeni/montanus clade was split into two species, one corresponding to the constrained Southern Philippine Z. montanus and the other containing the contrained Z. meyeni lineages with Z. japonicus and Z. montanus from the Northern Philippines.

### 3.7 Linear morphometrics

Seven linear morphometric measures (culmen, nare-to-tip bill length, bill depth, bill width, tarsus, wing cord, and tail length) were recorded and analyzed from 354 museum specimens (Supplementary table 5). Consistent trends were observed in morphology across species. *Z. japonicus* specimens were significantly larger across several measures and within *Z. japonicus* the oceanic subspecies *stejnegeri* and *alani* were larger still. Continental species (*Z. simplex*, *Z. erythropleurus*, and *Z. palpebrosus*) tended to be statistically smaller than the island species. The oceanic subspecies of *Z. japonicus* were statistically larger in at least 9/10 pairwise comparisons for culmen, nare-to-tip bill length, tarsus, wing cord, and tail length. The oceanic *Z. japonicus* were larger than *Z. erythropleurus* in every measure except for wing cord. Also, the *Z. nigrorum catarmanensis* subspecies on Camiguin South was significantly larger than other *Z. nigrorum* specimens measured in every metric except for tail length (Bonferroni corrected Wilcox rank sum test, p<0.05, Figure 7, Supplementary table 8).

**Figure 7:**
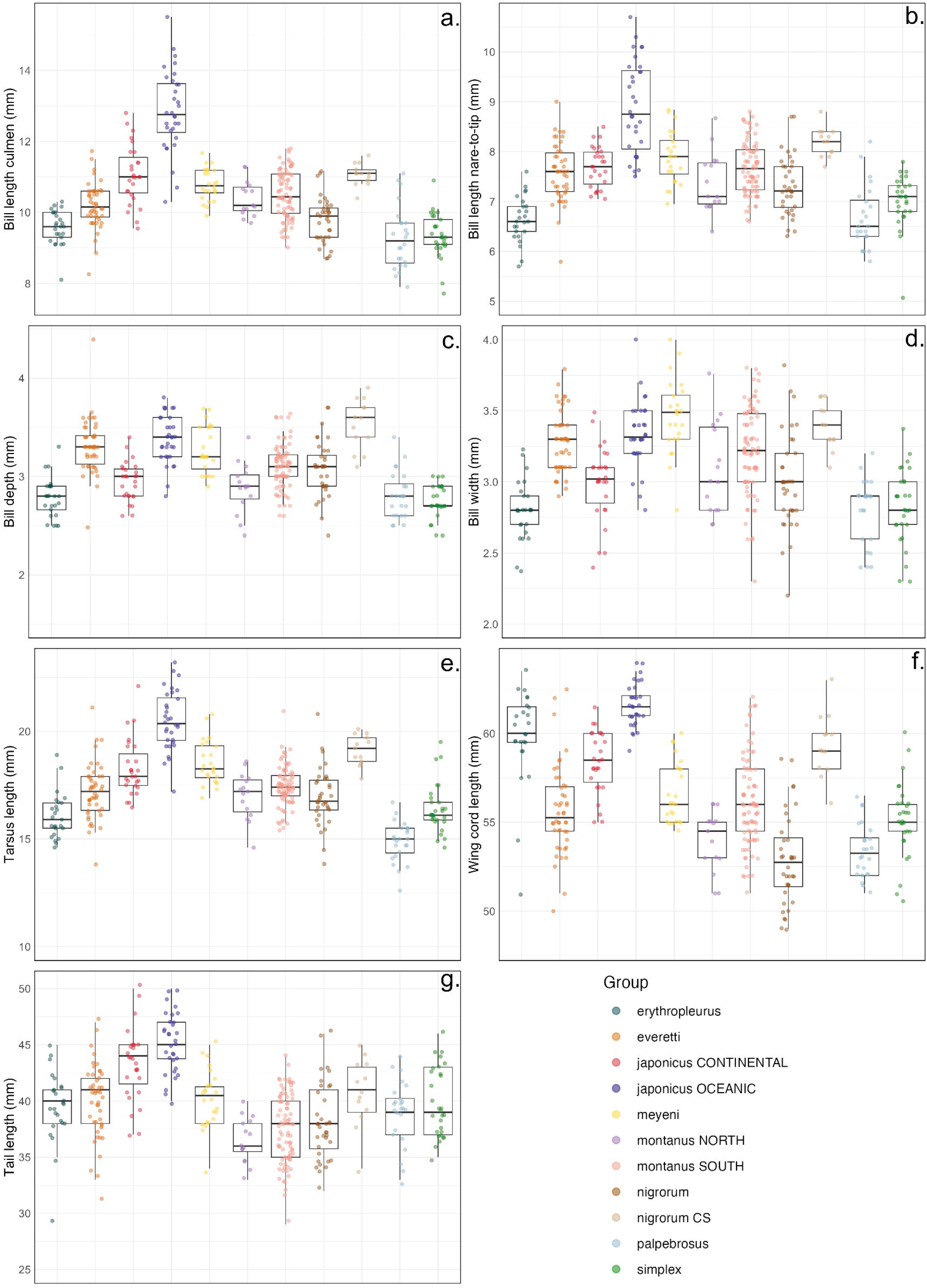
Box-and-whisker plots for seven linear morphological measures recorded from 354 museum study skins. All indiviual points are shown with some jitter to avoid too much visual overlap. Skins where poor condition or positioning excluded accurate measurements for one or more traits were excluded from analyses. Culmen length was measured as exposed culmen from the tip of the bill to the point where the bill meets the forehead plumage (a). Bill length nare-to-tip was measured from the distal end of the right nare to the tip of the bill (b) and bill depth (c) and width (d) were measured at the nares. Tarsometatarsus (tarsus) length was measured from the base of the phalanges to the tibiotarsal joint (e). Wing cord was measured unflattened (f) with a wing rule. Tail length (g) was measured from the center of the retrices using the same wing rule. Individual study skins were grouped by species and geography. The ‘japonicus CONTINENTAL’ group includes those populations of Z.japonicus on the main islands of Japan, the Korean Peninsula and adjacent islands, the Ryukyu archipelago, and Daito Islands. No specimens from either introduced populations in Hawaii or California, USA were included in these analyses. The ‘japonicus OCEANIC’ group included subspecies of Z. japonicus found on oceanic island chains in the Northern Pacfic including Z. japonicus stejnegeri in the lzu Islands and Z. japonicus alani in the Ogasawara and Volcano Islands. The ‘montanusNORTH’ group included those subspecies of Z. montanus on the islands of Luzon (Z. montanus whiteheadi) and Mindoro (Z. montanus halconensis) in the northern Philippines. The’montanus SOUTH’ group included subspecies and geographic locales for Z. montanus including found in the Visayas in the central Philippines, Mindanao and adjacent islands in the southern Philippines, and Indonesia. The’nigrorum CS’ group represents the Z. nigrorum catarmanensis subspecies found on the island of Camiguin Sur in the southern Philippines and discovered in phylogenetic analyses in this study to be mismatched in a clade with Z. montanus rather than with other Z. nigrorum populations. The reader should note that the y-axis for each set of plots does not begin at zero and careful attention should be paid to the scale when ascertaining differences between groups.

A PCA with groups designated according to species revealed somewhat distinct but spatially overlapping clusters. *Z. japonicus* formed a cluster overlapping with other species but extending further into morphospace (Figure 8a). Highlighting the *Z. japonicus* cluster showed that there was decreasing overlap with phylogenetic distance with the most overlap between *Z. japonicus* and *Z. meyeni* and the least between *Z. japonicus* and *Z. palpebrosus* (Figures 8b-e). However, while an estimate of Spearman’s rank correlation coefficient between pairwise comparisons of Euclidean distance from the morphometric and genetic (SNP) datasets revealed a trend towards increasing morphological distance with greater genetic distance (Figure 8f), this trend was not statistically significant either for all pairwise species comparisons (ρ = 0.18, p = 0.36) or for only those pairwise comparisons involving *Z. japonicus* (ρ = 0.54, p = 0.24).

**Figure 8:**
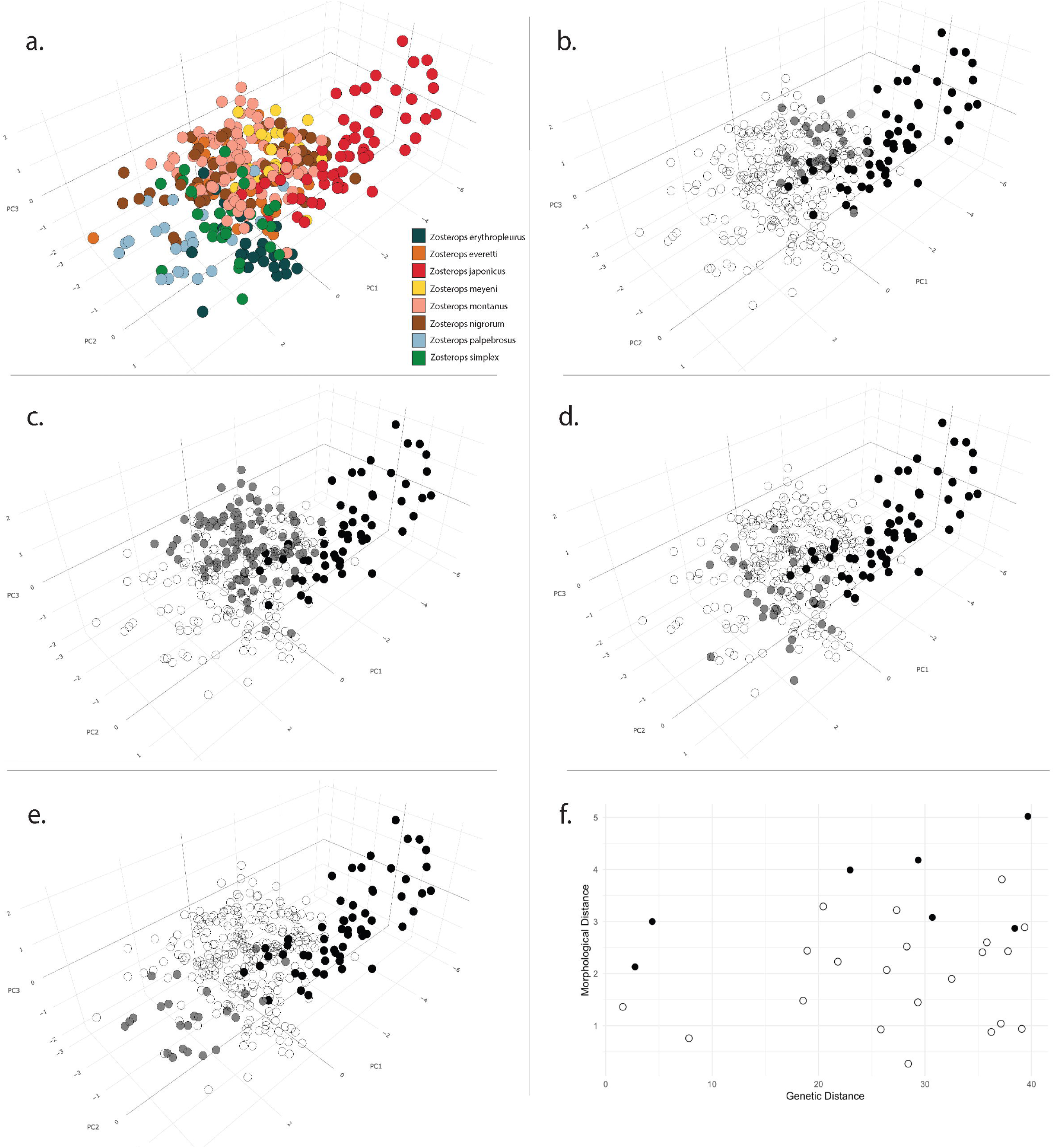
PCA based on seven linear morphological measures from 354 museum study skins. A plot of the first three PCs color coded by species appears in the upper left corner (a). Plots b - e highlight pairwise comparisons between Z. japonicus in black and four species highlighted in grey: Z. meyeni (b), Z. montanus (c), Z. simplex (d), and Z. palpebrosus (e). Other species on plots b - e are displayed as open circles. Also shown is a plot of the Euclidean distances for pairwise species comparisons of centroids based on the first three PCs from morphological and SNP-based genetic PCAs. Solid points are used for pairwise comparisons involving Z. japonicus. All other comparisons are indicated by open circles (f).

## 4. Discussion

### 4.1 The first Zosterops radiation in the Philippines and a new species

The Asiatic white-eyes underwent a rapid Pleistocene radiation that included two independent divergences in the Philippines and a rapid colonization of virtually all the islands of Japan. While caution is warranted when drawing conclusions from a molecular clock, our estimates from the allelic data, albeit imprecise, were consistent with the timing estimated from other studies (Moyle et al., 2009; Vinciguerra et al., 2023). The older divergence in the Philippines resulted in the more southerly distributed *Z. everetti* in the Visayas on Siquijor, Cebu, Samar, and Leyte, as well as Mindanao and the Sulu Archipelago and a more northerly distributed *Z. nigrorum* on the islands of central Philippines, including Panay and Negros in the Visayas, Cagayancillo, and Luzon and its adjacent islands. The presence of *Z. nigrorum* on Panay and Negros appears to exclude *Z. everetti* (Kennedy et al., 2000; Allen 2020).

Our study supports a species split between *Z. nigrorum nigrorum* on Panay and a northern clade including the subspecies on Luzon (*Z. nigrorum innominatus* and *Z. nigrorum luzonicus*) and the adjacent island of Camiguin North (*Z. nigrorum meyleri*). Deep divergences among the *Z. nigrorum* subspecies were found even in the poorly resolved Sanger sequencing trees. SNP-based trees produced well supported and comparatively deep splits between *Z. nigrorum nigrorum* on Panay and the other more recently derived subspecies. The divergence between *Z. nigrorum nigrorum* and the other *Z. nigrorum* subspecies was nearly as great as that between *Z. nigrorum* and its sister, *Z. everetti*. Additionally, when *Z. nigrorum nigrorum* was left unconstrained in Delineate it was consistently identified as a species lineage. Jones and Kennedy (2008) documented biogeographic patterns for *Ficedula hyperthra* and *Phylloscopus trivirgatus* like those in the *Z. nigrorum*/*everetti* clade with a northern clade consisting of populations on Panay, Negros, and Luzon and a more southerly clade on Mindanao. We believe that elevating the nominate *Z. nigrorum* subspecies on Panay to species should be considered, perhaps following additional genetic sampling and phenotypic analyses.

### 4.2 Zosterops nigrorum on Camiguin South is Z. montanus

Balete *et al*. (2006) record *Z. everetti* and *Z. nigrorum* on Camiguin South and identify the *Z. nigrorum* population as the endemic island subspecies, *Z. nigrorum catarmanensis*. Camiguin South is the only locale with sympatric *Z. everetti* and *Z. nigrorum*. *Z. nigrorum catarmanensis* represents a morphologically larger and geographically disjunct population of *Z. nigrorum* separated from the rest of the species by the islands of Cebu and Bohol where the species is absent (Rand & Rabor 1969; Allen 2020; van Balen 2020). Our study found strong support across Sanger sequencing, SNPs, and morphometrics that this population is in fact not *Z. nigrorum* but instead a yellow morph of *Z. montanus*. *Z. nigrorum* is unique among the Asiatic white-eyes in possessing all yellow underparts which is why the population of yellow white-eyes on Camiguin South were identified as *Z. nigrorum*. However, our study revealed not only that it is genetically mismatched, strongly sorting with *Z. montanus* from the Southern Philippines, but also that it is significantly larger than its conspecifics. The *Z. nigrorum catarmanensis* specimens in our study are closer in size to *Z. montanus*, and for some measures larger still. Yellow morphs are common in *Zosterops* and have confounded taxonomic arrangements. Other yellow subspecies are already recognized within *Z. montanus* including *Z. montanus obstinatus* and *Z. montanus difficilis* in the Muluku Islands (Moluccas) and Sumatra (Hartert 1900; Van Riper & van Balen 2020). Additionally, a yellow morph is present in populations of *Zosterops palpebrosus,* and identified as the *siamensis* subspecies in Myanmar and Western Thailand (Round et al., 2017). This result fits with reoccurring story of discordance between plumage and genetics in passerine birds where color polymorphism and evolutionary convergence often generates taxonomic confusion (e.g. Ng et al., 2022; Penalba et al., 2022; Garg et al., 2023).

It is not surprising that this phylogenetic mismatch is occurring on Camiguin South. Camiguin South is a geologically young island approximately 10km north of Minandao separated by a deep marine trench. The volcanic island was likely never connected to mainland Mindanao in its approximately 1Ma year history and is considered among the most unique centers of Philippine biodiversity (Heaney & Tabaranza Jr 2006). Modern discoveries of two endemic rodents (Heaney & Tabaranza Jr 2002; Heaney 2006) and one parrot (Tello et al., 2006) has revealed the island as the smallest area of island endemicism in the Philippines. Here we reveal that the large, yellow white-eyes on Camiguin South historically attributed to *Z. nigrorum catarmanensis* have been misclassified and instead represent the most phenotypically divergent population of *Z. montanus* in the Philippines.

### 4.3 A rapid second radiation of Zosterops in the Philippines and Japan

For decades, we have known the evolutionary relationships between *Z. japonicus*, *Z. meyeni*, and *Z. montanus* are complex (Van Riper & van Balen 2020). Throughout this study we have been using *Z. montanus* to distinguish populations in the Philippines, the Greater Sundas (excluding Borneo), the Lesser Sundas, and the Maluku Islands (Moluccas) including Seram, Buru, Obi, Bacan, Tidore, and Ternate from *Z. japonicus* on the islands of Japanese Archipelago, Ryukyu Islands, Korean Peninsula, Izu Islands, Ogasawara Islands, and Volcano Islands. However, as of 2019, *Z. montanus* and *Z. japonicus* were merged with priority retaining the name *Z. japonicus* for the entire group. While this rearrangement created a single white-eye species from Java to Hokkaido it retained separate species status of *Z. meyeni* (Clements et al., 2019). This taxonomy remains in place as of publication of the 2024 eBird/Clements list and the IOC 2025 v15.1 list (Clements et al., 2024; Gill et al., 2025).

This is a curious taxonomic arrangement for two reasons. First, at odds with their reputation as “great speciators”, a lumped *Z. japonicus*/*montanus* species creates the most latitudinally widespread breeding range for any Zosterops. In comparison, Diamond *et al*.’s (1976) original granting of the “great speciator” designation to the *Zosterops* [*griseotincta*] superspecies complex was in reference to the Solomon Islands where they concluded this group formed as many as six species, one of which with three subspecies. Recent studies have also documented a species-level radiation in the Solomons with single island endemics and significant isolation and differentiation even among the small and geographically proximate islands of the New Georgia group, separated by as little as 12 km (Manthey et al., 2020). Two altitudinally separate and reproductively isolated white-eyes (*Z. kulambangrae* and *Z. murphyi*) have evolved *in situ* on the 15 km wide island of Kolombangara in the Solomon Archipelago (Cowles & Uy 2019). Why then does *Z. japonicus* constitute a single species stretched from Java to Hokkaido, separated by thousands of kilometers of open ocean and fragmented island archipelagos?

One explanation for a single combined species lineage for *Z. japonicus*, *Z. meyeni*, *and Z. montanus* may simply be age. Our study shows that this clade represents a very rapid radiation, especially in Japan. The northern clade consisting of the *Z. montanus* subspecies on Luzon and Mindoro, *Z. meyeni*, and *Z. japonicus* in the Japanese Archipelago were distinguished in the SNP dataset by a strongly supported node in the UPGMA tree (87%), and SVDQuartets (91%). Although the divergence of this northern clade from the southern *Z. montanus* was recent (between 0.06 and 0.13 coalescent units from the present), it was statistically distinguishable from zero in both runs of PICL. Relative to the split times estimated by PICL for the other, more well-delimited, taxa, the divergence between these northern and southern clades is very recent.

Species delimitation analysis in *Delineate* overwhelmingly tended to lump *Z. japonicus*, *Z. meyeni*, and *Z. montanus* into a single species lineage. *Delineate* however identified novel species from among our samples (i.e., the *Z. nigrorum nigrorum* population on Panay). The sensitivity of *Delineate* seems to lie somewhere between the deepest split within *Z. nigrorum* and the split between the northern and southern groups within the *Z. japonicus*/*meyeni*/*montanus* clade. Roux *et al*. (2016) examined the relationship between ongoing migration and genetic divergence and discovered a “grey zone” between 0.5 – 2% divergence where lineages may be considered “semi-isolated”. The northern and southern groups within *Z. japonicus*/*meyeni*/*montanus* may currently be in this “grey zone”. Lineage sorting and reproductive isolation may have yet to occur in this recent expansion and despite the strong support in SVDQuartets many of these clades may be more indicative of structure as opposed to species.

Another explanation may be that there are isolated, evolutionarily independent lineages that may be delimited as species within this group, and we are looking at the wrong data, not enough of the right data, or using methodologies not equipped to detect the differences. For all its flaws the Sanger sequencing dataset nonetheless offered some insights by resolving a well-supported, monophyletic *Z. japonicus* and a GDI for *Z. meyeni* approached that of *Z. simplex*. For the SNP data DAPC readily distinguishes *Z. japonicus*, *Z. meyeni*, the northern *Z. montanus*, and the southern *Z. montanus* from one another as well as it did for the well-established species in the analysis without over splitting within *Z. japonicus*. DAPC also identified the same mismatched sample of *Z. nigrorum catarmanensis* flagged in every other analysis. Species splits within *Z. japonicus*/*meyeni*/*montanus* were revealed within the 95% cumulative probability for the least constrained analysis in *Delineate* but with low probability compared to lumping these groups. Finally, morphometrics distinguished *Z. japonicus* from other species across nearly every measure suggesting evidence of phenotypic evolution in this group.

A second curiosity involves *Z. meyeni*. Species limits between *Z. japonicus*, *Z. meyeni*, and *Z. montanus* are ambiguous at best. We found the same paraphyly in *Z. montanus* relative to *Z. japonicus* as in other studies with a topology consistent with geographic proximity (Jones & Kennedy 2008; Lim et al., 2019). This result led to the subsuming of *Z. montanus* into *Z. japonicus* based on nomenclatural priority, but *Z. meyeni* is consistently embedded in this group, including in this study, and yet has been retained as a separate species lineage.

There are three possible taxonomic resolutions to this problem. First, lump all these lineages into a single widespread and variable species. Taken to its logical conclusion this approach would include subsuming *Z. meyeni* into *Z. japonicus*. Second, delimit two species the first comprising a northern lineage including *Z. montanus* subspecies on Luzon and Mindoro, *Z. meyeni*, and *Z. japonicus* and the second the southerly populations of *Z. montanus* in the Visayas, Palawan, southern Philippines, Indonesia.

Finally, solve the problem of paraphyly in *Z. montanus* by elevating those populations in the northern Philippines to their own species while maintaining *Z. meyeni* and *Z. japonicus*. This solution has the appeal of keeping *Z. japonicus* distinct in recognition of its larger body size, darker coloration on the flanks, and consistent and distinct grouping even with genetic data with limited power (i.e., mtDNA). The first solution is the most conservative and likely the most difficult to reject as genetics seems to unite these lineages and the phenotypic diversity within this group seems mostly clinal and variable. The third solution is perhaps the weakest as genetic support for a monophyletic northern Philippine *Z. montanus* is poor and little phenotypically separates each lineage, particularly the northern *Z. montanus* and *Z. meyeni*, although *Z. japonicus* tends to be more distinct.

The second solution deserves further investigation. A phylogenetic break between a northern and southern group separating *Z. montanus* in the south from a northern group consisting of *Z. montanus* in the northern Philippines, *Z. meyeni*, and *Z. japonicus* is partially supported by the data. Split times for this break are small but significantly different from zero and while poorly supported relative to lumping these groups into a single species, this north-south split was within the 95% cumulative probability for the *Delineate* analysis. Again, this circles back to the perennial problem in species delimitation, namely distinguishing structure from species (McKay et al., 2013; Sukumaran & Knowles 2017). Our genetic dataset was comparatively modest in size and there are many other potential morphological, behavioral, and ecological characters to consider in this group. Future work comparing more SNPs from alignment of whole genomes and additional phenotypic variation potentially involved in reproductive isolation such as plumage color and song could provide additional tests of this hypothesis and more persuasively resolve the lingering species questions in Asiatic *Zosterops*.

### 4.4 Potential for speciation within Z. japonicus and Z. montanus

Another question we sought to address in this study was the evolutionary validity of the subspecies within *Z. japonicus*, and to a lesser degree within *Z. montanus*. The only distinction that could be arguably made for divisions within *Z. japonicus* from this study is regarding population structure between the oceanic populations, including *Z. japonicus stejnegeri* in the Izu Islands and *Z. japonicus alani* in the Ogasawara and Volcano Islands, and those populations on the Ryukyu Islands, the main islands of Japan, and the Korean Peninsula. Oceanic populations of *Z. japonicus* exhibited larger body size compared to continental populations and SVDQuartets strongly supported a distinct *Z. japonicus alani*, the most geographically isolated of all the subspecies. However, this variation is subtle and DAPC could not distinguish between *Z. japonicus alani* and the other *Z. japonicus* and, while well supported, we could not support a split time for *Z. japonicus alani* that was statistically different than zero in PICL.

This result is consistent with other studies. Hamao *et al*. (2013), for example, found little evidence of divergence in mtDNA or song among those populations of *Z. japonicus* in the Ryukyu Islands. The history of *Z. japonicus* in the oceanic islands of Japan is likely very complex and deserves additional study. The most geographically remote of any of the Asiatic white-eyes are the *Z. japonicus alani* populations in the Volcano Islands. At approximately 1,216km south of Tokyo, Iwoto (formerly Iwojima) is among the most isolated of these islands.

Zosterops have a documented capacity for oceanic dispersal. Ely (1971) recorded 11 pelagic sightings of *Z. japonicus* in the North Pacific (most likely originating from introduced source populations in Hawaii) and observed at least five individuals on Johnston Atoll, approximately the same distance from Honolulu (1,326km) as Iwoto is from Tokyo. Selection seems to have favored particular characters in these remote populations of *Z. japonicus*. Oceanic island populations of *Z. japonicus* have wing cord measures similar to that of *Z. erythropleurus*, the only long-distance migrant species in Zosterops. This is unlikely to be due to an allometric relationship as the continental *Z. erythropleurus* is among the smallest species in overall body size. Long wings are likely a trait favored both in long-distance migration (Outlaw 2011) and in oceanic dispersal.

This capacity for dispersal coupled with the propensity of humans to transport white-eyes either deliberately in the cage bird trade or inadvertently as stowaways has likely resulted in some gene flow even in geographically isolated populations. Sugita *et al*. (2016) found unique mtDNA haplotpyes in the *Z. japonicus alani* population in the Volcano Islands. Haplotypes shared with Volcano, Ryukyu, and Izu Island populations were found in the Ogasawara population, which is more populated and closer to Mainland Japan. This suggested some degree of mitochondrial gene flow taking place among these subspecies either due to introductions or natural dispersal. Together these studies and our own suggest that considerable gene flow is still taking place among populations in the islands of Japan with meaningful isolation occurring only on the most far-flung oceanic islands.

We were less able to confidentially draw conclusions regarding variation within *Z. montanus* but here too there seems to be very little genetic distinctiveness among subspecies, perhaps with the exception of *Z. montanus parkesi* on Palawan. We were unable to sample two subspecies of *Z. montanus* with either remote or range restricted distributions. Both are phenotypically distinct from most other *Z. montanus* with yellow underparts, and both have subspecific names that reflect the taxonomic confusion associated with each (*Z. montanus obstinatus* and *Z. montanus difficillis*). These obstinate and difficult populations deserve further study, as does *Z. montanus* throughout Indonesia.

### 4.5 Testing new analytical tools in complex datasets

This study is an important case study as it represents the first application of PICL on a complex empirical dataset. The use of composite likelihoods proved to be a computationally efficient approach to phylogenetic inference under the multispecies coalescent without the need to first estimate gene trees (Kubatko et al., 2025).

Approaches that are locus-based, estimating species trees from gene tree input, are poorly suited for SNP data. We anticipate that PICL will fill in this methodological gap in an environment where large SNP datasets in systematics are becoming commonplace.

### 4.6 Conclusions

Two waves of colonization characterize the Zosterops radiation throughout the island archipelagos of the Philippines and East Asia. The first of which is comparatively old and morphologically distinct and the second of which very recent and so far, poorly differentiated. These data pose many interesting questions. *Z. japonicus* in the southern Ryukyus is more distinct from *Z. simplex* in Taiwan, less than 200km distant, than they are from *Z. montanus* in Java, more than 3,600km distant. It remains a puzzle why a “great speciator” with documented species divergence between islands separated by as little as 12 km (Manthey et al., 2020) would be so difficult to differentiate between Indonesia and the Philippines or much less between the tropical archipelagos of the Philippines and Indonesia and the temperate islands of Japan. Within this second diversification, *Z. japonicus* populations of the Ryukyu Islands, Japan, the Korean Peninsula, and oceanic islands of the Western Pacific provide a good model for an island archipelago radiation in its early stages. We believe this study provides a baseline for future investigations of secondary contact, paraphyly, local adaptation, and biogeography in Zosterops, especially for the white-eyes of Japan and the Philippines.

## Funding

Herman L. Mays Jr was supported by NSF-MCA #2322123. Robert G. Moyle was supported by NSF-DEB #1557053. High performance computing at the University of Kansas Center for Research Computing was supported by NSF-MRI #2117449. Samples obtained for *Z. japonicus alani* from the Volcano Islands were obtained by Kazuto Kawakami with support from the Tokyo Metropolitan Government.

## Acknowledgements

This paper is dedicated to the memory of Robert S. Kennedy and his contributions to Philippine ornithology and conservation and his stewardship of the zoology collection at Cincinnati Museum Center. Valuable advice for running Delineate was provided by Jeet Sukumaran. Thanks to curators and collections managers for facilitating museum visits and tissue loans including Paul Sweet, Thomas Trombone, and Brian Smith (American Museum of Natural History, New York), Heather Farrington (Cincinnati Museum Center, Cincinnati), Ben Marks and John Bates (Field Museum of Natural History, Chicago), Allison Shultz (Natural History Museum of Los Angeles County, Los Angeles), Christopher Milensky (Smithsonian National Museum of Natural History, Washington D.C.), Lucas DeCicco (University of Kansas Biodiversity Institute and Natural History Museum, Lawrence), and Sharon Birks (University of Washington Burke Museum, Seattle).

## Supporting information

Additional supporting information as well as complete workflows and data analysis are available at https://hermmays.github.io/Zosterops1/. Data are also available on Dryad (**PENDING**). Sanger sequenced loci are available at NCBI (**ACCESSION NUMBERS PENDING**). The raw sequence data for each sample passing filtering protocols is available via NCBI’s Sequence Read Archive, at the BioProject PRJNA1079333, which can be found at: https://www.ncbi.nlm.nih.gov/sra/PRJNA1079333.

## Notes

https://hermmays.github.io/Zosterops1/

### Competing Interest Statement

The authors have declared no competing interest.

