## Supplemental tables for "The not-so-great speciator: Systematics and species limits in a rapid radiation, the Asiatic white-eye complex (*Zosterops spp.*)"

| Source | Catalog number | Alternate ID number | Project number | Alternate project number | Taxon | Locale | Lat | Long | Sanger | SNP |
|---|---|---|---|---|---|---|---|---|---|---|
| AMNH, USA | DOT12589 | RWD24671 | ZAsu001 | NA | *Zosterops atrifrons surdus* | Sulawesi, Indonesia | -1.6469 | 120.4358 | YES | NO |
| AMNH, USA | DOT12620 | RWD24710 | ZAsu002 | NA | *Zosterops atrifrons surdus* | Sulawesi, Indonesia | -0.7435 | 123.0114 | YES | NO |
| UWBM, USA | UWBM71717 | DRF144 | ZERxx001 | NA | *Zosterops erythropleurus* | Primorsky Krai, Russia | 42.97 | 132.89 | YES | NO |
| UWBM, USA | UWBM71926 | IVF241 | ZERxx002 | NA | *Zosterops erythropleurus* | Primorsky Krai, Russia | 46.02 | 135.25 | YES | YES |
| UWBM, USA | UWBM71981 | JML208 | ZERxx003 | NA | *Zosterops erythropleurus* | Primorsky Krai, Russia | 44.75 | 132.77 | YES | NO |
| UWBM, USA | UWBM72103 | SVD2446 | ZERxx004 | NA | *Zosterops erythropleurus* | Primorsky Krai, Russia | 44.92 | 131.89 | YES | YES |
| UWBM, USA | UWBM75187 | RYA486 | ZERxx005 | NA | *Zosterops erythropleurus* | Primorsky Krai, Russia | 44.3674 | 133.0265 | YES | YES |
| KU, USA | KU120458 | KU28088 (tissue) | Zery28088 | NA | *Zosterops erythropleurus* | Lao Cai, Vietnam | 21.949 | 104.255 | NO | YES |
| KU, USA | KU120457 | KU28090 (tissue) | Zery28090 | NA | *Zosterops erythropleurus* | Lao Cai, Vietnam | 21.949 | 104.255 | NO | YES |
| KU, USA | KU120460 | KU28091 (tissue) | Zery28091 | NA | *Zosterops erythropleurus* | Lao Cai, Vietnam | 21.949 | 104.255 | NO | YES |
| CMC, USA | NMP/CMNHB1641 | NA | ZEba001 | NA | *Zosterops everetti basilanicus* | Mindanao, Philippines | 7.0189 | 125.4984 | YES | NO |
| KU, USA | KU110475 | KU13949 (tissue) | ZEba002 | Zev13949 | *Zosterops everetti basilanicus* | Camiguin Sur, Philippines | 9.1925 | 124.7085 | YES | YES |
| KU, USA | KU122172 | KU28451 (tissue) | ZEba003 | Zev28451 | *Zosterops everetti basilanicus* | Mindanao, Philippines | 9.074 | 125.642 | YES | YES |
| KU, USA | KU123410 | KU31650 (tissue) | ZEbo001 | Zev31650 | *Zosterops everetti boholensis* | Samar, Philippines | 11.828 | 125.276 | YES | YES |
| NMNS, Japan | NSMT-DNA6000 | 2006.222 | ZJal001 | NA | *Zosterops japonicus alani* | Hahajima, Japan | 26.6735 | 142.1541 | YES | NO |
| NMNS, Japan | NSMT-DNA5999 | 2006.223 | ZJal002 | NA | *Zosterops japonicus alani* | Hahajima, Japan | 26.6735 | 142.1541 | YES | NO |
| NMNS, Japan | NSMT-DNA5952 | 2S64071ZJ | ZJal003 | NA | *Zosterops japonicus alani* | Hahajima, Japan | 26.6735 | 142.1541 | YES | YES |
| NMNS, Japan | NSMT-DNA5953 | 2S64072ZJ | Zjal004 | NA | *Zosterops japonicus alani* | Hahajima, Japan | 26.6735 | 142.1541 | YES | YES |
| NMNS, Japan | NSMT-DNA5954 | 2S64073ZJ | ZJal005 | NA | *Zosterops japonicus alani* | Hahajima, Japan | 26.6735 | 142.1541 | YES | YES |
| NMNS, Japan | NSMT-DNA6025 | ZJ2X21900 | ZJal006 | NA | *Zosterops japonicus alani* | Imotojima, Japan | 26.5587 | 142.2085 | YES | NO |
| NMNS, Japan | NSMT-DNA6026 | ZJ2X21901 | ZJal007 | NA | *Zosterops japonicus alani* | Imotojima, Japan | 26.5587 | 142.2085 | YES | NO |
| NMNS, Japan | NSMT-DNA6028 | ZJ2X21904 | ZJal008 | NA | *Zosterops japonicus alani* | Imotojima, Japan | 26.5587 | 142.2085 | YES | NO |
| NMNS, Japan | NSMT-DNA6034 | ZJ2X21905 | ZJal009 | NA | *Zosterops japonicus alani* | Imotojima, Japan | 26.5587 | 142.2085 | YES | NO |
| NMNS, Japan | NSMT-DNA6068 | ZJ2X21913 | ZJal010 | NA | *Zosterops japonicus alani* | Anejima, Japan | 26.5506 | 142.1581 | YES | YES |
| NMNS, Japan | NSMT-DNA6069 | ZJ2X21914 | ZJal011 | NA | *Zosterops japonicus alani* | Anejima, Japan | 26.5506 | 142.1581 | YES | YES |
| NMNS, Japan | NSMT-DNA6070 | ZJ2X21915 | ZJal012 | NA | *Zosterops japonicus alani* | Anejima, Japan | 26.5506 | 142.1581 | YES | YES |
| NMNS, Japan | NSMT-DNA6071 | ZJ2X21916 | ZJal013 | NA | *Zosterops japonicus alani* | Anejima, Japan | 26.5506 | 142.1581 | YES | NO |

| | | | | | | | | | | |
|---|---|---|---|---|---|---|---|---|---|---|
| NMNS, Japan | NSMT-DNA6072 | ZJ2X21917 | ZJal014 | NA | *Zosterops japonicus alani* | Anejima, Japan | 26.5506 | 142.1581 | YES | NO |
| NMNS, Japan | NSMT-DNA6027 | ZJ2X21903 | ZJal015 | NA | *Zosterops japonicus alani* | Imotojima, Japan | 26.5587 | 142.2085 | YES | NO |
| NMNS, Japan | NSMT-DNA9893 | ZJ2AA00993 | ZJal016 | NA | *Zosterops japonicus alani* | Iwoto, Japan | 24.7876 | 141.3156 | YES | NO |
| NMNS, Japan | NSMT-DNA9896 | ZJ2AA00996 | ZJal019 | NA | *Zosterops japonicus alani* | Iwoto, Japan | 24.7876 | 141.3156 | YES | NO |
| NMNS, Japan | NSMT-DNA9905 | ZJ2AA01127 | ZJal020 | NA | *Zosterops japonicus alani* | Iwoto, Japan | 24.7876 | 141.3156 | NO | YES |
| NMNS, Japan | NSMT-DNA9234 | 1308 | ZJlo001 | NA | *Zosterops japonicus alani* | Chichijima, Japan | 27.0744 | 142.2178 | YES | NO |
| NMNS, Japan | NSMT-DNA9235 | 1309 | ZJlo002 | NA | *Zosterops japonicus alani* | Chichijima, Japan | 27.0744 | 142.2178 | YES | NO |
| NMNS, Japan | NSMT-DNA9236 | 1310 | ZJlo003 | NA | *Zosterops japonicus alani* | Chichijima, Japan | 27.0744 | 142.2178 | YES | NO |
| NMNS, Japan | NSMT-DNA9237 | 1311 | ZJlo004 | NA | *Zosterops japonicus alani* | Chichijima, Japan | 27.0744 | 142.2178 | YES | NO |
| NMNS, Japan | NSMT-DNA9238 | 1312 | ZJlo005 | NA | *Zosterops japonicus alani* | Chichijima, Japan | 27.0744 | 142.2178 | YES | NO |
| NMNS, Japan | NSMT-DNA140763 | 2011.171 | ZJin001 | NA | *Zosterops japonicus insularis* | Kagoshima, Japan | 30.3715 | 130.6646 | YES | YES |
| NMNS, Japan | NSMT-DNA140654 | 2AC67744 | ZJin003 | NA | *Zosterops japonicus insularis* | Kagoshima, Japan | 30.3715 | 130.6646 | YES | YES |
| USNM, USA | NA | BRY431 | Z_HI_BRY431 | NA | *Zosterops japonicus japonicus* | Hawaii, USA | 19.566 | -155.453 | NO | YES |
| USNM, USA | NA | NAN290 | Z_HI_NAN290 | NA | *Zosterops japonicus japonicus* | Hawaii, USA | 19.566 | -155.453 | NO | YES |
| USNM, USA | NA | SOL783 | Z_HI_SOL783 | NA | *Zosterops japonicus japonicus* | Hawaii, USA | 19.566 | -155.453 | NO | YES |
| USNM, USA | NA | WAI078 | Z_HI_WAI078 | NA | *Zosterops japonicus japonicus* | Hawaii, USA | 19.566 | -155.453 | NO | YES |
| USNM, USA | NA | WAI087 | Z_HI_WAI087 | NA | *Zosterops japonicus japonicus* | Hawaii, USA | 19.566 | -155.453 | NO | YES |
| NMNS, Japan | NSMT-DNA2804 | 020-12131 | ZJja001 | NA | *Zosterops japonicus japonicus* | Geoje, Republic of Korea | 34.8806 | 128.6211 | YES | YES |
| NMNS, Japan | NSMT-DNA2806 | 020-12133 | ZJja002 | NA | *Zosterops japonicus japonicus* | Geoje, Republic of Korea | 34.8806 | 128.6211 | YES | YES |
| NMNS, Japan | NSMT-DNA2814 | 020-12141 | ZJja003 | NA | *Zosterops japonicus japonicus* | Jeju, Republic of Korea | 33.489 | 126.4983 | YES | YES |
| NMNS, Japan | NSMT-DNA2818 | 020-12144 | ZJja004 | NA | *Zosterops japonicus japonicus* | Jeju, Republic of Korea | 33.489 | 126.4983 | YES | YES |
| NMNS, Japan | NSMT-DNA2820 | 020-12145 | ZJja005 | NA | *Zosterops japonicus japonicus* | Jeju, Republic of Korea | 33.489 | 126.4983 | YES | YES |
| NMNS, Japan | NSMT-DNA2821 | 020-12146 | ZJja006 | NA | *Zosterops japonicus japonicus* | Jeju, Republic of Korea | 33.489 | 126.4983 | YES | NO |
| NMNS, Japan | NSMT-DNA2830 | 020-12155 | ZJja007 | NA | *Zosterops japonicus japonicus* | Jeju, Republic of Korea | 33.489 | 126.4983 | YES | NO |
| NMNS, Japan | NSMT-DNA2931 | 1210 | ZJja008 | NA | *Zosterops japonicus japonicus* | Honshu, Japan | 35.6388 | 139.7256 | YES | NO |
| NMNS, Japan | NSMT-DNA3518 | 12169 | ZJja009 | NA | *Zosterops japonicus japonicus* | Heuksando, Republic of Korea | 34.6707 | 125.4196 | YES | YES |
| NMNS, Japan | NSMT-DNA3526 | 12177 | ZJja010 | NA | *Zosterops japonicus japonicus* | Heuksando, Republic of Korea | 34.6707 | 125.4196 | YES | YES |
| NMNS, Japan | NSMT-DNA3527 | 12178 | ZJja011 | NA | *Zosterops japonicus japonicus* | Heuksando, Republic of Korea | 34.6707 | 125.4196 | YES | NO |

| | | | | | | | | | | |
|---|---|---|---|---|---|---|---|---|---|---|
| NMNS, Japan | NSMT-DNA1630 | 1999.27 | ZJja012 | NA | *Zosterops japonicus japonicus* | Honshu, Japan | 36.0835 | 140.0764 | YES | YES |
| NMNS, Japan | NSMT-DNA1696 | 2001.2 | ZJja013 | NA | *Zosterops japonicus japonicus* | Honshu, Japan | 36.0835 | 140.0764 | YES | YES |
| NMNS, Japan | NSMT-DNA3063 | 2003.78 | ZJja014 | NA | *Zosterops japonicus japonicus* | Honshu, Japan | 35.9086 | 139.6248 | YES | YES |
| NMNS, Japan | NSMT-DNA4257 | 2005.56 | ZJja015 | NA | *Zosterops japonicus japonicus* | Honshu, Japan | 35.7263 | 139.7168 | YES | NO |
| NMNS, Japan | NSMT-DNA4847 | 2006.100 | ZJja016 | NA | *Zosterops japonicus japonicus* | Shikoku, Japan | 34.3708 | 133.9198 | YES | YES |
| NMNS, Japan | NSMT-DNA5859 | 2006.175 | ZJja017 | NA | *Zosterops japonicus japonicus* | Honshu, Japan | 36.6024 | 137.6088 | YES | NO |
| NMNS, Japan | NSMT-DNA6662 | 2007.107 | ZJja018 | NA | *Zosterops japonicus japonicus* | Hokkaido, Japan | 42.9393 | 143.4477 | YES | NO |
| NMNS, Japan | NSMT-DNA7137 | 2007.189 | ZJja019 | NA | *Zosterops japonicus japonicus* | Honshu, Japan | 36.3402 | 139.4498 | YES | NO |
| NMNS, Japan | NSMT-DNA8065 | 2008.273 | ZJja021 | NA | *Zosterops japonicus japonicus* | Honshu, Japan | 35.6388 | 139.7256 | YES | NO |
| NMNS, Japan | NSMT-DNA8756 | 2009.105 | ZJja022 | NA | *Zosterops japonicus japonicus* | Honshu, Japan | 37.2832 | 138.9816 | YES | NO |
| NMNS, Japan | NSMT-DNA4920 | 2S99902 | ZJja024 | NA | *Zosterops japonicus japonicus* | Honshu, Japan | 36.7846 | 136.8348 | YES | NO |
| NMNS, Japan | NSMT-DNA4254 | 2Y57213ZJ | ZJja026 | NA | *Zosterops japonicus japonicus* | Honshu, Japan | 33.703 | 135.3759 | YES | NO |
| NMNS, Japan | NSMT-DNA3575 | ZJ02022 | ZJja030 | NA | *Zosterops japonicus japonicus* | Heuksando, Republic of Korea | 34.6707 | 125.4196 | NO | YES |
| NMNS, Japan | NSMT-DNA8045 | ZJ2S63651 | ZJja031 | NA | *Zosterops japonicus japonicus* | Honshu, Japan | 35.7032 | 139.6975 | YES | NO |
| NMNS, Japan | NSMT-DNA8046 | ZJ2S63652 | ZJja032 | NA | *Zosterops japonicus japonicus* | Honshu, Japan | 35.7032 | 139.6975 | YES | NO |
| AMNH, USA | DOT19086 | BDM354 | ZJja034 | NA | *Zosterops japonicus japonicus* | Tsushima, Japan | 34.4081 | 129.286 | YES | NO |
| AMNH, USA | DOT19087 | BDM355 | Zjja035 | NA | *Zosterops japonicus japonicus* | Tsushima, Japan | 34.4081 | 129.286 | YES | NO |
| NMNS, Japan | NSMT-DNA4884 | 1C70840 | ZJlo006 | NA | *Zosterops japonicus loochooensis* | Iriomote, Japan | 24.3301 | 123.8188 | YES | NO |
| NMNS, Japan | NSMT-DNA3592 | 2004.56 | ZJlo007 | NA | *Zosterops japonicus loochooensis* | Okinawa, Japan | 26.7456 | 128.1779 | YES | NO |
| NMNS, Japan | NSMT-DNA3593 | 2004.57 | ZJlo008 | NA | *Zosterops japonicus loochooensis* | Okinawa, Japan | 26.7456 | 128.1779 | YES | NO |
| NMNS, Japan | NSMT-DNA7724 | 2008.108 | ZJlo009 | NA | *Zosterops japonicus loochooensis* | Okinawa, Japan | 26.7456 | 128.1779 | YES | NO |
| NMNS, Japan | NSMT-DNA7566 | 2008.52 | ZJlo010 | NA | *Zosterops japonicus loochooensis* | Ishigaki, Japan | 24.4064 | 124.1754 | YES | NO |
| NMNS, Japan | NSMT-DNA7551 | 2008.68 | ZJlo011 | NA | *Zosterops japonicus loochooensis* | Ishigaki, Japan | 24.4064 | 124.1754 | YES | NO |
| NMNS, Japan | NSMT-DNA7692 | 2008.76 | ZJlo012 | NA | *Zosterops japonicus loochooensis* | Kikai, Japan | 28.317 | 129.9401 | YES | YES |
| NMNS, Japan | NSMT-DNA7693 | 2008.77 | ZJlo013 | NA | *Zosterops japonicus loochooensis* | Kikai, Japan | 28.317 | 129.9401 | YES | NO |
| NMNS, Japan | NSMT-DNA8754 | 2009.103 | ZJlo014 | NA | *Zosterops japonicus loochooensis* | Iriomote, Japan | 24.3301 | 123.8188 | YES | NO |
| NMNS, Japan | NSMT-DNA8778 | 2009.130 | ZJlo015 | NA | *Zosterops japonicus loochooensis* | Amami, Japan | 28.3192 | 129.2765 | YES | YES |
| NMNS, Japan | NSMT-DNA8889 | 2009.186 | ZJlo016 | NA | *Zosterops japonicus loochooensis* | Miyakojima, Japan | 24.7916 | 125.3302 | YES | YES |

| | | | | | | | | | | |
|---|---|---|---|---|---|---|---|---|---|---|
| NMNS, Japan | NSMT-DNA8962 | 2009.232 | ZJlo017 | NA | *Zosterops japonicus loochooensis* | Iriomote, Japan | 24.3301 | 123.8188 | YES | YES |
| NMNS, Japan | NSMT-DNA8974 | 2009.244 | ZJlo018 | NA | *Zosterops japonicus loochooensis* | Iriomote, Japan | 24.3301 | 123.8188 | YES | NO |
| NMNS, Japan | NSMT-DNA8976 | 2009.246 | ZJlo020 | NA | *Zosterops japonicus loochooensis* | Iriomote, Japan | 24.3301 | 123.8188 | YES | NO |
| NMNS, Japan | NSMT-DNA9620 | 2010.114 | ZJlo021 | NA | *Zosterops japonicus loochooensis* | Tokunoshima, Japan | 27.7904 | 128.9668 | YES | YES |
| NMNS, Japan | NSMT-DNA9634 | 2010.118 | ZJlo022 | NA | *Zosterops japonicus loochooensis* | Tokunoshima, Japan | 27.7904 | 128.9668 | YES | YES |
| NMNS, Japan | NSMT-DNA9655 | 2010.126 | ZJlo023 | NA | *Zosterops japonicus loochooensis* | Okinoerabu, Japan | 27.5737 | 128.5737 | YES | NO |
| NMNS, Japan | NSMT-DNA9552 | 2010.141 | ZJlo024 | NA | *Zosterops japonicus loochooensis* | Yonaguni, Japan | 24.468 | 123.0045 | YES | YES |
| NMNS, Japan | NSMT-DNA9187 | 2AA77978ZJ | ZJlo026 | NA | *Zosterops japonicus loochooensis* | Miyakojima, Japan | 24.8218 | 125.3137 | YES | NO |
| NMNS, Japan | NSMT-DNA9555 | 2AC67722ZJ | ZJlo027 | NA | *Zosterops japonicus loochooensis* | Yonaguni, Japan | 24.468 | 123.0045 | YES | NO |
| NMNS, Japan | NSMT-DNA9726 | 2AC67728ZJ | ZJlo031 | NA | *Zosterops japonicus loochooensis* | Kumejima, Japan | 26.3504 | 126.7712 | YES | YES |
| NMNS, Japan | NSMT-DNA9732 | 2AC67731ZJ | ZJlo032 | NA | *Zosterops japonicus loochooensis* | Kumejima, Japan | 26.3504 | 126.7712 | YES | NO |
| NMNS, Japan | NSMT-DNA9383 | MIY12 | ZJlo034 | NA | *Zosterops japonicus loochooensis* | Miyakojima, Japan | 24.7354 | 125.3 | YES | NO |
| NMNS, Japan | NSMT-DNA6467 | YA14 | ZJlo035 | NA | *Zosterops japonicus loochooensis* | Okinawa, Japan | 26.7456 | 128.1779 | YES | NO |
| NMNS, Japan | NSMT-DNA7586 | ZJ-Okinawa3 | ZJlo038 | NA | *Zosterops japonicus loochooensis* | Okinawa, Japan | 26.7456 | 128.1779 | YES | NO |
| NMNS, Japan | NSMT-DNA7552 | ZJ-Ishigaki1 | ZJlo039 | NA | *Zosterops japonicus loochooensis* | Ishigaki, Japan | 24.4064 | 124.1754 | YES | NO |
| NMNS, Japan | NSMT-DNA8779 | ZJ1E57254 | ZJlo042 | NA | *Zosterops japonicus loochooensis* | Amami, Japan | 28.3192 | 129.2765 | YES | NO |
| NMNS, Japan | NSMT-DNA8782 | ZJ1E57259 | ZJlo045 | NA | *Zosterops japonicus loochooensis* | Amami, Japan | 28.3192 | 129.2765 | YES | YES |
| NMNS, Japan | NSMT-DNA9205 | ZJ1E57295 | ZJlo046 | NA | *Zosterops japonicus loochooensis* | Miyakojima, Japan | 24.8218 | 125.3137 | NO | YES |
| NMNS, Japan | NSMT-DNA9661 | ZJ1F40445 | ZJlo050 | NA | *Zosterops japonicus loochooensis* | Okinoerabu, Japan | 27.5737 | 128.5737 | NO | YES |
| NMNS, Japan | NSMT-DNA9665 | ZJ1F40446 | ZJlo051 | NA | *Zosterops japonicus loochooensis* | Okinoerabu, Japan | 27.5737 | 128.5737 | YES | NO |
| NMNS, Japan | NSMT-DNA9666 | ZJ1F40447 | ZJlo052 | NA | *Zosterops japonicus loochooensis* | Okinoerabu, Japan | 27.5737 | 128.5737 | NO | YES |
| NMNS, Japan | NSMT-DNA8027 | ZJ_4 | ZJlo055 | NA | *Zosterops japonicus loochooensis* | Kikai, Japan | 28.317 | 129.9401 | YES | YES |
| NMNS, Japan | NSMT-DNA8034 | ZJ_11 | ZJst001 | NA | *Zosterops japonicus stejnegeri* | Niijima, Japan | 34.3771 | 139.2569 | YES | NO |
| NMNS, Japan | NSMT-DNA2361 | 1D36701ZJ | ZJst002 | NA | *Zosterops japonicus stejnegeri* | Aogashima, Japan | 32.457 | 139.7652 | YES | NO |
| NMNS, Japan | NSMT-DNA2378 | 1D36706ZJ | ZJst006 | NA | *Zosterops japonicus stejnegeri* | Aogashima, Japan | 32.457 | 139.7652 | YES | NO |
| NMNS, Japan | NSMT-DNA2090 | 2D05461ZJ | ZJst007 | NA | *Zosterops japonicus stejnegeri* | Kozushima, Japan | 34.2055 | 139.1345 | YES | YES |
| NMNS, Japan | NSMT-DNA2092 | 2D05463ZJ | ZJst008 | NA | *Zosterops japonicus stejnegeri* | Kozushima, Japan | 34.2055 | 139.1345 | YES | NO |
| NMNS, Japan | NSMT-DNA2093 | 2D05464ZJ | ZJst009 | NA | *Zosterops japonicus stejnegeri* | Kozushima, Japan | 34.2055 | 139.1345 | YES | YES |

| | | | | | | | | | | |
|---|---|---|---|---|---|---|---|---|---|---|
| NMNS, Japan | NSMT-DNA2094 | 2D05465ZJ | ZJst010 | NA | *Zosterops japonicus stejnegeri* | Kozushima, Japan | 34.2055 | 139.1345 | YES | YES |
| NMNS, Japan | NSMT-DNA2095 | 2D05466ZJ | ZJst011 | NA | *Zosterops japonicus stejnegeri* | Kozushima, Japan | 34.2055 | 139.1345 | YES | NO |
| NMNS, Japan | NSMT-DNA1971 | 2D05483ZJ | ZJst012 | NA | *Zosterops japonicus stejnegeri* | Miyakejima, Japan | 34.0788 | 139.5178 | YES | YES |
| NMNS, Japan | NSMT-DNA1972 | 2D05484ZJ | ZJst013 | NA | *Zosterops japonicus stejnegeri* | Miyakejima, Japan | 34.0788 | 139.5178 | YES | NO |
| NMNS, Japan | NSMT-DNA1974 | 2D05486ZJ | ZJst014 | NA | *Zosterops japonicus stejnegeri* | Miyakejima, Japan | 34.0788 | 139.5178 | YES | YES |
| NMNS, Japan | NSMT-DNA1981 | 2D05490ZJ | ZJst015 | NA | *Zosterops japonicus stejnegeri* | Miyakejima, Japan | 34.0788 | 139.5178 | YES | YES |
| NMNS, Japan | NSMT-DNA1992 | 2D60608ZJ | ZJst016 | NA | *Zosterops japonicus stejnegeri* | Miyakejima, Japan | 34.0788 | 139.5178 | YES | NO |
| NMNS, Japan | NSMT-DNA1835 | ZJ_3 | ZJst017 | NA | *Zosterops japonicus stejnegeri* | Niijima, Japan | 34.3771 | 139.2569 | YES | NO |
| NMNS, Japan | NSMT-DNA2841 | A23889 | ZJba001 | NA | *Zosterops meyeni batanis* | Lanyu Island, Taiwan | 22.0436 | 121.5484 | YES | NO |
| NMNS, Japan | NSMT-DNA2923 | A23890 | ZJba002 | NA | *Zosterops meyeni batanis* | Lanyu Island, Taiwan | 22.0436 | 121.5484 | YES | NO |
| KU, USA | KU112216 | KU17852 (tissue) | ZMxx001 | Zmey17852 | *Zosterops meyeni batanis* | Batan Island, Philippines | 20.457 | 121.99 | YES | YES |
| KU, USA | KU112108 | KU17876 (tissue) | Zmey17876 | NA | *Zosterops meyeni batanis* | Batan Island, Philippines | 20.47 | 121.991 | NO | YES |
| KU, USA | KU112109 | KU17877 (tissue) | Zmey17877 | NA | *Zosterops meyeni batanis* | Batan Island, Philippines | 20.47 | 121.991 | NO | YES |
| KU, USA | KU113064 | KU17920 (tissue) | Zmey17920 | NA | *Zosterops meyeni batanis* | Sabtang Island, Philippines | 20.285 | 121.871 | NO | YES |
| KU, USA | KU112377 | KU17922 (tissue) | Zmey17922 | NA | *Zosterops meyeni batanis* | Sabtang Island, Philippines | 20.285 | 121.871 | NO | YES |
| KU, USA | KU112378 | KU17923 (tissue) | Zmey17923 | NA | *Zosterops meyeni batanis* | Sabtang Island, Philippines | 20.285 | 121.871 | NO | YES |
| KU, USA | KU112110 | KU17925 (tissue) | Zmey17925 | NA | *Zosterops meyeni batanis* | Sabtang Island, Philippines | 20.285 | 121.871 | NO | YES |
| KU, USA | KU113362 | KU17865 (tissue) | ZMxx002 | NA | *Zosterops meyeni batanis* | Batan Island, Philippines | 20.47 | 121.991 | YES | YES |
| AMNH, USA | DOT19042 | BDM310 | ZMxx003 | NA | *Zosterops meyeni batanis* | Lanyu Island, Taiwan | 22.0811 | 121.5048 | YES | YES |
| AMNH, USA | DOT19043 | BDM311 | ZMxx004 | NA | *Zosterops meyeni batanis* | Lanyu Island, Taiwan | 22.0811 | 121.5048 | YES | YES |
| AMNH, USA | DOT19044 | BDM312 | ZMxx005 | NA | *Zosterops meyeni batanis* | Lanyu Island, Taiwan | 22.0811 | 121.5048 | YES | YES |
| AMNH. USA | DOT19046 | BDM314 | ZMxx006 | NA | *Zosterops meyeni batanis* | Lanyu Island, Taiwan | 22.0093 | 121.5721 | YES | YES |
| AMNH, USA | DOT19048 | BDM316 | ZMxx007 | NA | *Zosterops meyeni batanis* | Lanyu Island, Taiwan | 22.0093 | 121.5721 | YES | NO |
| AMNH, USA | DOT19056 | BDM324 | ZMxx008 | NA | *Zosterops meyeni batanis* | Lanyu Island, Taiwan | 22.0093 | 121.5721 | YES | NO |
| KU, USA | KU28375 | NA | Zmon28375 | ZMOdi001 | *Zosterops montanus diuatae* | Mindanao, Philippines | 9.079 | 125.665 | YES | YES |
| CMC, USA | CMCB36979 | NMP/CMNHB980 | ZMOha001 | NA | *Zosterops montanus halconensis* | Mindoro, Philippines | 13.3082 | 121.0765 | YES | YES |
| AMNH, USA | DOT12552 | NA | ZMOmo001 | NA | *Zosterops montanus montanus* | Sulawesi, Indonesia | -5.4081 | 119.8989 | YES | YES |
| KU, USA | KU129029 | KU31171 (tissue) | ZMOmo004 | NA | *Zosterops montanus montanus* | Java, Indonesia | -7.6145 | 110.7122 | YES | NO |

| | | | | | | | | | | |
|---|---|---|---|---|---|---|---|---|---|---|
| KU, USA | KU129021 | KU31159 (tissue) | ZMOmo002 | Zsim31159 | *Zosterops montanus montanus* | Java, Indonesia | -7.6145 | 110.7122 | YES | YES |
| KU, USA | KU129026 | KU31166 (tissue) | ZMOmo003 | Zsim31166 | *Zosterops montanus montanus* | Java, Indonesia | -7.6145 | 110.7122 | YES | YES |
| FMNH, USA | FMNH455085 | NA | ZMOpa001 | NA | *Zosterops montanus parkesi* | Palawan, Philippines | 8.8183 | 117.675 | YES | YES |
| FMNH, USA | FMNH455086 | NA | ZMOpa002 | NA | *Zosterops montanus parkesi* | Palawan, Philippines | 8.8183 | 117.675 | YES | YES |
| KU, USA | KU116591 | KU20892 (tissue) | Zmon20892 | Zmon20892 | *Zosterops montanus pectoralis* | Negros, Philippines | 10.6743 | 123.1886 | NO | YES |
| KU, USA | KU116590 | KU20893 (tissue) | ZMOpe001 | Zmon20893 | *Zosterops montanus pectoralis* | Negros, Philippines | 10.6743 | 123.1886 | YES | YES |
| KU, USA | KU116887 | KU20899 (tissue) | ZMOpe003 | Zmon20899 | *Zosterops montanus pectoralis* | Negros, Philippines | 10.6743 | 123.1886 | YES | YES |
| KU, USA | KU116593 | KU20902 (tissue) | Zmon20902 | NA | *Zosterops montanus pectoralis* | Negros, Philippines | 10.6743 | 123.1886 | NO | YES |
| KU, USA | KU116605 | KU20909 (tissue) | ZMOpe002 | Zmon20909 | *Zosterops montanus pectoralis* | Negros, Philippines | 10.6655 | 123.1787 | YES | YES |
| CMC, USA | NMP/CMNHB426 | NA | ZMOxx001 | NA | *Zosterops montanus ssp.* | Panay, Philippines | 11.4062 | 122.1235 | YES | YES |
| CMC, USA | NMP/CMNHB442 | NA | ZMOxx002 | NA | *Zosterops montanus ssp.* | Panay, Philippines | 11.4062 | 122.1235 | YES | YES |
| CMC, USA | NMP/CMNHB495 | NA | ZMOxx003 | NA | *Zosterops montanus ssp.* | Panay, Philippines | 11.3936 | 122.1661 | YES | YES |
| CMC, USA | NMP/CMNHB596 | NA | ZMOxx004 | NA | *Zosterops montanus ssp.* | Panay, Philippines | 11.3936 | 122.1661 | YES | NO |
| CMC, USA | CMCB36819 | NMP/CMNHB556 | ZMOxx005 | NA | *Zosterops montanus ssp.* | Panay, Philippines | 11.4062 | 122.1235 | YES | YES |
| CMC, USA | CMCB36894 | NMP/CMNHB704 | ZMOxx006 | NA | *Zosterops montanus ssp.* | Panay, Philippines | 11.4109 | 122.2881 | YES | NO |
| CMC, USA | CMCB36896 | NMP/CMNHB769 | ZMOxx007 | NA | *Zosterops montanus ssp.* | Panay, Philippines | 11.4109 | 122.2881 | YES | NO |
| CMC, USA | CMCB36898 | NMP/CMNHB801 | ZMOxx008 | NA | *Zosterops montanus ssp.* | Panay, Philippines | 11.4109 | 122.2881 | YES | NO |
| CMC, USA | NMP/CMNHB1354 | NA | ZMOvu001 | NA | *Zosterops montanus vulcani* | Mindanao, Philippines | 7.0189 | 125.4984 | YES | NO |
| CMC, USA | CMCB35836 | RSK94B3154 | ZMOvu002 | NA | *Zosterops montanus vulcani* | Mindanao, Philippines | 7.8514 | 126.0458 | YES | YES |
| CMC, USA | CMCB35884 | RSK94B3240 | ZMOvu003 | NA | *Zosterops montanus vulcani* | Mindanao, Philippines | 7.8514 | 126.0458 | YES | YES |
| CMC, USA | CMCB38163 | NMP/CMNHB1249 | ZMOvu004 | NA | *Zosterops montanus vulcani* | Mindanao, Philippines | 7.0189 | 125.4984 | YES | YES |
| CMC, USA | CMCB39029 | NMP/CMNHB1711 | ZMOvu005 | NA | *Zosterops montanus vulcani* | Mindanao, Philippines | 7.0189 | 125.4984 | YES | YES |
| CMC, USA | CMCB39198 | NMP/CMNHB2178 | ZMOvu006 | NA | *Zosterops montanus vulcani* | Mindanao, Philippines | 6.0234 | 124.6414 | YES | NO |
| CMC, USA | CMCB39200 | NMP/CMNHB2250 | ZMOvu007 | NA | *Zosterops montanus vulcani* | Mindanao, Philippines | 6.0234 | 124.6414 | YES | NO |
| CMC, USA | CMCB36615 | NMP/CMNHB342 | ZMOwh002 | NA | *Zosterops montanus whiteheadi* | Luzon, Philippines | 15.5286 | 119.9608 | YES | YES |
| FMNH, USA | FMNH429299 | NA | ZMOwh003 | NA | *Zosterops montanus whiteheadi* | Luzon, Philippines | 17.467 | 121.067 | YES | YES |
| FMNH, USA | FMNH446736 | NA | ZMOwh004 | NA | *Zosterops montanus whiteheadi* | Luzon, Philippines | 15.4819 | 120.1194 | YES | YES |
| FMNH, USA | FMNH454952 | NA | ZMOwh005 | NA | *Zosterops montanus whiteheadi* | Luzon, Philippines | 17.0373 | 121.101 | YES | YES |

| | | | | | | | | | | |
|---|---|---|---|---|---|---|---|---|---|---|
| FMNH, USA | FMNH462072 | NA | ZMOwh006 | NA | *Zosterops montanus whiteheadi* | Luzon, Philippines | 16.5976 | 120.8986 | YES | YES |
| KU, USA | KU110362 | KU14341 (tissue) | ZNca001 | Zni14341 | *Zosterops nigrorum catarmanensis* | Camiguin Sur, Philippines | 9.178 | 124.7197 | YES | YES |
| KU, USA | KU114596 | KU19650 (tissue) | ZNin001 | Zni19650 | *Zosterops nigrorum innominatus* | Luzon, Philippines | 15.6436 | 121.5149 | YES | YES |
| KU, USA | KU112603 | KU17984 (tissue) | ZNlu001 | Zni17984 | *Zosterops nigrorum luzonicus* | Luzon, Philippines | 14.039 | 122.786 | YES | YES |
| KU, USA | KU97296 | KU10863 (tissue) | ZNme001 | Zni10863 | *Zosterops nigrorum meyleri* | Camiguin Norte, Philippines | 18.929 | 121.899 | YES | YES |
| CMC, USA | CMCB36816 | NA | ZNni001 | NA | *Zosterops nigrorum nigrorum* | Panay, Philippines | 11.4062 | 122.1235 | YES | YES |
| AMNH, USA | AMNH833731 | DOT10981 | ZjaDOT-10981 | NA | *Zosterops palpebrosus* | Ha Giang, Vietnam | 22.7601 | 104.8805 | NO | YES |
| KU, USA | KU119438 | KU23498 (tissue) | Zpal23498 | NA | *Zosterops palpebrosus* | Dien Bien, Vietnam | 22.386 | 102.238 | NO | YES |
| KU, USA | KU119437 | KU23522 (tissue) | Zpal23522 | NA | *Zosterops palpebrosus* | Dien Bien, Vietnam | 22.386 | 102.238 | NO | YES |
| KU, USA | KU122986 | KU30897 (tissue) | Zsim30897 | NA | *Zosterops palpebrosus* | Kon Tum, Vietnam | 15.11 | 107.824 | NO | YES |
| LACM, USA | LACM122188 | NA | Z_LA_122188_2 | NA | *Zosterops simplex* | Los Angeles County, California, USA | 33.6966 | -118.018 | NO | YES |
| LACM, USA | LACM120577 | NA | Z_LA_122577_2 | NA | *Zosterops simplex* | Los Angeles County, California, USA | 33.6966 | -118.018 | NO | YES |
| LACM, USA | LACM122866 | NA | Z_LA_122866_2 | NA | *Zosterops simplex* | Los Angeles County, California, USA | 33.6966 | -118.018 | NO | YES |
| AMNH, USA | DOT5235 | GFB3386 | ZjaDOT-5235 | NA | *Zosterops simplex* | Taiwan | 23.828 | 120.7925 | NO | YES |
| NMNS, Japan | NSMT-DNA3126 | A26848 | ZJsi001 | NA | *Zosterops simplex* | Taiwan | 22.7613 | 121.1438 | YES | NO |
| NMNS, Japan | NSMT-DNA2844 | A26867 | ZJsi002 | NA | *Zosterops simplex* | Taiwan | 22.7613 | 121.1438 | NO | YES |
| NMNS, Japan | NSMT-DNA2846 | A26869 | ZJsi003 | NA | *Zosterops simplex* | Taiwan | 22.7613 | 121.1438 | YES | NO |
| NMNS, Japan | NSMT-DNA2848 | A26871 | ZJsi005 | NA | *Zosterops simplex* | Taiwan | 22.7613 | 121.1438 | YES | NO |
| GAS, China | S00239 | NA | ZJsi015 | NA | *Zosterops simplex* | Macao, China | 22.1987 | 113.5439 | YES | NO |
| GAS, China | S00247 | NA | ZJsi017 | NA | *Zosterops simplex* | Guangxi, China | 22.8152 | 108.3275 | NO | YES |
| GAS, China | S00391 | NA | ZJsi018 | NA | *Zosterops simplex* | Sichuan, China | 30.6509 | 104.0757 | YES | NO |
| GAS, China | S01120 | NA | ZJsi023 | NA | *Zosterops simplex* | Guangdong, China | 23.1317 | 113.2663 | NO | YES |
| GAS, China | S01121 | NA | ZJsi024 | NA | *Zosterops simplex* | Guangdong, China | 23.1317 | 113.2663 | NO | YES |
| GAS, China | S01129 | NA | ZJsi025 | NA | *Zosterops simplex* | Guangdong, China | 23.1317 | 113.2663 | YES | NO |
| GAS, China | S01469 | NA | ZJsi026 | NA | *Zosterops simplex* | Yunnan, China | 25.0453 | 102.7097 | YES | NO |
| GAS, China | S01791 | NA | ZJsi027 | NA | *Zosterops simplex* | Guangxi, China | 22.8152 | 108.3275 | NO | YES |
| GAS, China | S01792 | NA | ZJsi028 | NA | *Zosterops simplex* | Guangxi, China | 22.8152 | 108.3275 | NO | YES |
| GAS, China | S01800 | NA | ZJsi029 | NA | *Zosterops simplex* | Guangxi, China | 22.8152 | 108.3275 | NO | YES |

| | | | | | | | | | | |
|---|---|---|---|---|---|---|---|---|---|---|
| GAS, China | S01802 | NA | ZJsi031 | NA | *Zosterops simplex* | Guangxi, China | 22.8152 | 108.3275 | NO | YES |
| GAS, China | S01803 | NA | ZJsi032 | NA | *Zosterops simplex* | Guangxi, China | 22.8152 | 108.3275 | NO | YES |
| UWBM, USA | UWBM76175 | CSW6561 | ZPxx001 | NA | *Zosterops simplex* | Singapore | 1.3189 | 103.8512 | NO | YES |
| UWBM, USA | UWBM117271 | KLE625 | ZPxx002 | NA | *Zosterops simplex* | Singapore | 1.3521 | 103.8198 | NO | YES |
| KU, USA | KU96551 | KU10336 (tissue) | Zsim10336 | NA | *Zosterops simplex* | Guangxi, China | 21.84 | 107.88 | NO | YES |
| KU, USA | KU98946 | KU11102 (tissue) | Zsim11102 | NA | *Zosterops simplex* | Guizhou, China | 28.2261 | 107.1596 | NO | YES |
| KU, USA | KU98952 | KU11220 (tissue) | Zsim11220 | NA | *Zosterops simplex* | Guizhou, China | 28.2261 | 107.1596 | NO | YES |
| KU, USA | KU97426 | KU11362 (tissue) | Zsim11362 | NA | *Zosterops simplex* | Guizhou, China | 29.1668 | 107.575 | NO | YES |
| KU, USA | KU110336 | KU13773 (tissue) | Zsim13773 | NA | *Zosterops simplex* | Guizhou, China | 25.4114 | 107.8872 | NO | YES |
| KU, USA | KU99789 | KU13809 (tissue) | Zsim13809 | NA | *Zosterops simplex* | Guizhou, China | 25.4114 | 107.8872 | NO | YES |
| KU, USA | KU119436 | KU23588 (tissue) | Zsim23588 | NA | *Zosterops simplex* | Dien Bien, Vietnam | 22.386 | 102.238 | NO | YES |
| KU, USA | KU120461 | KU28142 (tissue) | Zsim28142 | NA | *Zosterops simplex* | Lao Cai, Vietnam | 21.949 | 104.255 | NO | YES |
| KU, USA | KU92711 | KU6797 (tissue) | Zsim6797 | NA | *Zosterops simplex* | Hunan, China | 28.4167 | 114.1167 | NO | YES |

Supplementary table 1: Sample list of 208 individual samples used for genetic analyses. In total 124 of these samples are represented in the single nucleotide polymorphism (SNP) dataset and 159 samples are represented by at least one locus in the Sanger sequencing dataset. 75 samples are represented in both the Sanger and SNP datasets. Because work was distributed among two different laboratories some samples were given different working project numbers. Both project numbers used for those samples are shown here. Alternate identifying (ID) numbers are variously field numbers, a corresponding number for a tissue collection that match a voucher number, or some other identifying number at that institution. NA indicates that some identifying code was not available. For some samples formal catalog numbers are absent. Museum collections abbreviated in the source column are as follows; American Museum of Natural History, New York, USA (AMNH, USA), Cincinnati Museum Center, Cincinnati, USA (CMC, USA), Field Museum of Natural History, Chicago, USA (FMNH, USA), Institute of Zoology of the Guangdong Academy of Sciences, Guangzhou, People's Republic of China (GAS, China), University of Kansas Biodiversity Institute and Natural History Museum (KU, USA), Natural History Museum of Los Angeles County, Los Angeles, USA (LACM, USA), National Museum of Nature and Science, Tokyo, Japan (NMNS, Japan), Smithsonian National Museum of Natural History, Washington DC, USA (USMN, USA), and the University of Washington Burke Museum, Seattle, USA (UWBM, USA). Latitude (Lat) and longitude (Long) were either obtained from museum records or georeferenced to an approximate location from the museum locality data using Berkeley Mapper (https://berkeleymapper.berkeley.edu/). For brevity, locale information here is limited to either island or province/state and country.

| Oligonucleotide name | 5'-CS tag + target sequence (5' → 3') | Locus description | Location/reference species | References |
|---|---|---|---|---|
| L5216-CS1 | ACACTGACGACATGGTTCTACAGGCCCATACCCCGRAAATG | NADH dehydrogenase subunit 2 (ND2) | mtDNA/avian | Sorenson et al. 1999 |
| H6313-CS2 | TACGGTAGCAGAGACTTGGTCTACTCCTRTTTAAGGCTTTGAAGGC | NADH dehydrogenase subunit 2 (ND2) | mtDNA/avian | Sorenson et al. 1999 |
| BIRDF1-CS1 | ACACTGACGACATGGTTCTACATTCTCCAACCACAAAGACATTGGCAC | Cytochrome c Oxidase Subunit 1 (COI) | mtDNA/avian | Hebert et al. 2004 |
| BIRDR2-CS2 | TACGGTAGCAGAGACTTGGTCTACTACATGTGAGATGATTCCGAATCCAG | Cytochrome c Oxidase Subunit 1 (COI) | mtDNA/avian | Hebert et al. 2004 |
| B3GNT9F-CS1 | ACACTGACGACATGGTTCTACACCATTGGAAAGGCTTCTTCAC | betaGal beta-1,3-N-acetylglucosaminyltransferase 9 | chromosome 11/*Taeniopygia guttata* | Liu et al. 2018 |
| B3GNT9R-CS2 | TACGGTAGCAGAGACTTGGTCTTTTCTGCTCATTGCTATCAAATC | betaGal beta-1,3-N-acetylglucosaminyltransferase 9 | chromosome 11/*Taeniopygia guttata* | Liu et al. 2018 |
| BIRC2F-CS1 | ACACTGACGACATGGTTCTACAATCTGCAAGCTGTTCAGGC | baculoviral IAP repeat containing 2 | chromosome 1/*Taeniopygia guttata* | Liu et al. 2018 |
| BIRC2R-CS2 | TACGGTAGCAGAGACTTGGTCTCCGGGCTGGGTTTTATTAC | baculoviral IAP repeat containing 2 | chromosome 1/*Taeniopygia guttata* | Liu et al. 2018 |
| CCDCF-CS1 | ACACTGACGACATGGTTCTACACCAAAGTTACTGGAGAACCTTCC | coiled-coil domain containing 141 | chromosome 7/*Taeniopygia guttata* | Liu et al. 2018 |
| CCDCR-CS2 | TACGGTAGCAGAGACTTGGTCTATACAGAGCTGTCATCTTCAGCC | coiled-coil domain containing 141 | chromosome 7/*Taeniopygia guttata* | Liu et al. 2018 |
| GPRC6F-CS1 | ACACTGACGACATGGTTCTACAACAACAAAACCTACTGGGC | G protein-coupled receptor, class C, group 6, member A | chromosome 3/*Taeniopygia guttata* | Liu et al. 2018 |
| GPRC6R-CS2 | TACGGTAGCAGAGACTTGGTCTCAGGGATAAATACAATCCATGCT | G protein-coupled receptor, class C, group 6, member A | chromosome 3/*Taeniopygia guttata* | Liu et al. 2018 |
| LACTBLF-CS1 | ACACTGACGACATGGTTCTACACCAGTGAGATGGAGGCAGAG | lactamase, beta-like 1 | chromosome 21/*Taeniopygia guttata* | Liu et al. 2018 |
| LACTBLR-CS2 | TACGGTAGCAGAGACTTGGTCTATCCGCTCCCAGATGGC | lactamase, beta-like 1 | chromosome 21/*Taeniopygia guttata* | Liu et al. 2018 |
| USP38F-CS1 | ACACTGACGACATGGTTCTACACCTGAGATCCTTAGTGGAGACAA | ubiquitin carboxyl-terminal hydrolase 38 | chromosome 4/*Taeniopygia guttata* | Liu et al. 2018 |
| USP38R-CS2 | TACGGTAGCAGAGACTTGGTCTGCATCCATGAGGTCTTTTTGTAG | ubiquitin carboxyl-terminal hydrolase 38 | chromosome 4/*Taeniopygia guttata* | Liu et al. 2018 |

Supplementary table 2: Polymerase chain reaction (PCR) primers for generating amplicons for Sanger sequencing. Each primer has a 5’ extension that facilitates amplicon sequencing of both DNA strands. Primers with the forward extension 5’-ACACTGACGACATGGTTCTACA are labeled with CS1. Primers with the reverse extension 5’-TACGGTAGCAGAGACTTGGTCT are labeled with CS2. Each primer is accompanied with a description of the locus as well as its chromosomal location. Nuclear coding genes are identified based on their location in the Zebra Finch (*Taeniopygia guttata*) genome.

| BPP control file | Inverse $\Gamma$ prior distribution $\alpha$ for $\theta$ and $\tau$ | Inverse $\Gamma$ distribution $\beta_\theta$ | Mean $\theta$ prior | Inverse $\Gamma$ distribution $\beta\tau$ | Mean $\tau$ prior | Model | burnin | sample | nsample | species delimitation algorithm | Result | posterior |
|---|---|---|---|---|---|---|---|---|---|---|---|---|
| ZosteropsA10_RUNA01.bpp.ctl | 3 | 0.01 | 0.005 | 0.002 | 0.001 | GTR | 10000 | 2 | 100000 | 1 | 1111 | 1 |
| ZosteropsA10_RUNA02.bpp.ctl | 3 | 0.02 | 0.01 | 0.002 | 0.001 | GTR | 10000 | 2 | 100000 | 1 | 1111 | 1 |
| ZosteropsA10_RUNA03.bpp.ctl | 3 | 0.1 | 0.5 | 0.002 | 0.001 | GTR | 10000 | 2 | 100000 | 1 | 1111 | 1 |
| ZosteropsA10_RUNA04.bpp.ctl | 3 | 0.01 | 0.005 | 0.0005 | 0.00025 | GTR | 10000 | 2 | 100000 | 1 | 1111 | 1 |
| ZosteropsA10_RUNA05.bpp.ctl | 3 | 0.02 | 0.01 | 0.0005 | 0.00025 | GTR | 10000 | 2 | 100000 | 1 | 1111 | 1 |
| ZosteropsA10_RUNA06.bpp.ctl | 3 | 0.1 | 0.5 | 0.0005 | 0.00025 | GTR | 10000 | 2 | 100000 | 1 | 1111 | 1 |
| ZosteropsA10_RUNA07.bpp.ctl | 3 | 0.01 | 0.005 | 0.002 | 0.001 | GTR | 10000 | 2 | 100000 | 0 | 1111 | 1 |
| ZosteropsA10_RUNA08.bpp.ctl | 3 | 0.02 | 0.01 | 0.002 | 0.001 | GTR | 10000 | 2 | 100000 | 0 | 1111 | 1 |
| ZosteropsA10_RUNA09.bpp.ctl | 3 | 0.1 | 0.5 | 0.002 | 0.001 | GTR | 10000 | 2 | 100000 | 0 | 1111 | 1 |
| ZosteropsA10_RUNA10.bpp.ctl | 3 | 0.01 | 0.005 | 0.0005 | 0.00025 | GTR | 10000 | 2 | 100000 | 0 | 1111 | 1 |
| ZosteropsA10_RUNA11.bpp.ctl | 3 | 0.02 | 0.01 | 0.0005 | 0.00025 | GTR | 10000 | 2 | 100000 | 0 | 1111 | 1 |
| ZosteropsA10_RUNA12.bpp.ctl | 3 | 0.1 | 0.5 | 0.0005 | 0.00025 | GTR | 10000 | 2 | 100000 | 0 | 1111 | 1 |
| ZosteropsA10_RUNA13.bpp.ctl | 3 | 0.001 | 0.0005 | 0.004 | 0.002 | GTR | 10000 | 2 | 100000 | 0 | 1111 | 1 |
| ZosteropsA10_RUNA14.bpp.ctl | 3 | 0.002 | 0.001 | 0.004 | 0.002 | GTR | 10000 | 2 | 100000 | 0 | 1111 | 1 |
| ZosteropsA10_RUNA15.bpp.ctl | 3 | 0.001 | 0.0005 | 0.0005 | 0.00025 | GTR | 10000 | 2 | 100000 | 0 | 1111 | 1 |
| ZosteropsA10_RUNA16.bpp.ctl | 3 | 0.002 | 0.001 | 0.0005 | 0.00025 | GTR | 10000 | 2 | 100000 | 0 | 1111 | 1 |

Supplementary table 3: Description of replicate runs of BPP for species delimitation on a fixed tree (A10).

| BPP control file | Inverse $\Gamma$ prior distribution $\alpha$ for $\theta$ and $\tau$ | Inverse $\Gamma$ distribution $\beta_\theta$ | Mean $\theta$ prior | Inverse $\Gamma$ distribution $\beta\tau$ | Mean $\tau$ prior | Tree |
|---|---|---|---|---|---|---|
| ZosteropsA00_RUNB1.bpp.ctl | 3 | 0.001 | 0.0005 | 0.004 | 0.002 | TIP |
| ZosteropsA00_RUNB2.bpp.ctl | 3 | 0.001 | 0.0005 | 0.004 | 0.002 | TIP |
| ZosteropsA00_RUNB3.bpp.ctl | 3 | 0.001 | 0.0005 | 0.004 | 0.002 | TIP |
| ZosteropsA00_RUNB4.bpp.ctl | 3 | 0.002 | 0.001 | 0.004 | 0.002 | TIP |
| ZosteropsA00_RUNB5.bpp.ctl | 3 | 0.002 | 0.001 | 0.004 | 0.002 | TIP |
| ZosteropsA00_RUNB6.bpp.ctl | 3 | 0.002 | 0.001 | 0.004 | 0.002 | TIP |
| ZosteropsA00_RUNB7.bpp.ctl | 3 | 0.001 | 0.0005 | 0.0005 | 0.00025 | TIP |
| ZosteropsA00_RUNB8.bpp.ctl | 3 | 0.001 | 0.0005 | 0.0005 | 0.00025 | TIP |
| ZosteropsA00_RUNB9.bpp.ctl | 3 | 0.001 | 0.0005 | 0.0005 | 0.00025 | TIP |
| ZosteropsA00_RUNB10.bpp.ctl | 3 | 0.002 | 0.001 | 0.0005 | 0.00025 | TIP |
| ZosteropsA00_RUNB11.bpp.ctl | 3 | 0.002 | 0.001 | 0.0005 | 0.00025 | TIP |
| ZosteropsA00_RUNB12.bpp.ctl | 3 | 0.002 | 0.001 | 0.0005 | 0.00025 | TIP |
| ZosteropsA00_RUNC1.bpp.ctl | 3 | 0.001 | 0.0005 | 0.004 | 0.002 | MID1 |
| ZosteropsA00_RUNC2.bpp.ctl | 3 | 0.001 | 0.0005 | 0.004 | 0.002 | MID1 |
| ZosteropsA00_RUNC3.bpp.ctl | 3 | 0.001 | 0.0005 | 0.004 | 0.002 | MID1 |
| ZosteropsA00_RUNC4.bpp.ctl | 3 | 0.002 | 0.001 | 0.004 | 0.002 | MID1 |
| ZosteropsA00_RUNC5.bpp.ctl | 3 | 0.002 | 0.001 | 0.004 | 0.002 | MID1 |
| ZosteropsA00_RUNC6.bpp.ctl | 3 | 0.002 | 0.001 | 0.004 | 0.002 | MID1 |
| ZosteropsA00_RUNC7.bpp.ctl | 3 | 0.001 | 0.0005 | 0.0005 | 0.00025 | MID1 |
| ZosteropsA00_RUNC8.bpp.ctl | 3 | 0.001 | 0.0005 | 0.0005 | 0.00025 | MID1 |
| ZosteropsA00_RUNC9.bpp.ctl | 3 | 0.001 | 0.0005 | 0.0005 | 0.00025 | MID1 |
| ZosteropsA00_RUNC10.bpp.ctl | 3 | 0.002 | 0.001 | 0.0005 | 0.00025 | MID1 |
| ZosteropsA00_RUNC11.bpp.ctl | 3 | 0.002 | 0.001 | 0.0005 | 0.00025 | MID1 |
| ZosteropsA00_RUNC12.bpp.ctl | 3 | 0.002 | 0.001 | 0.0005 | 0.00025 | MID1 |
| ZosteropsA00_RUND1.bpp.ctl | 3 | 0.001 | 0.0005 | 0.004 | 0.002 | MID2 |
| ZosteropsA00_RUND2.bpp.ctl | 3 | 0.001 | 0.0005 | 0.004 | 0.002 | MID2 |
| ZosteropsA00_RUND3.bpp.ctl | 3 | 0.001 | 0.0005 | 0.004 | 0.002 | MID2 |
| ZosteropsA00_RUND4.bpp.ctl | 3 | 0.002 | 0.001 | 0.004 | 0.002 | MID2 |
| ZosteropsA00_RUND5.bpp.ctl | 3 | 0.002 | 0.001 | 0.004 | 0.002 | MID2 |
| ZosteropsA00_RUND6.bpp.ctl | 3 | 0.002 | 0.001 | 0.004 | 0.002 | MID2 |
| ZosteropsA00_RUND7.bpp.ctl | 3 | 0.001 | 0.0005 | 0.0005 | 0.00025 | MID2 |
| ZosteropsA00_RUND8.bpp.ctl | 3 | 0.001 | 0.0005 | 0.0005 | 0.00025 | MID2 |
| ZosteropsA00_RUND9.bpp.ctl | 3 | 0.001 | 0.0005 | 0.0005 | 0.00025 | MID2 |
| ZosteropsA00_RUND10.bpp.ctl | 3 | 0.002 | 0.001 | 0.0005 | 0.00025 | MID2 |
| ZosteropsA00_RUND11.bpp.ctl | 3 | 0.002 | 0.001 | 0.0005 | 0.00025 | MID2 |
| ZosteropsA00_RUNC12.bpp.ctl | 3 | 0.002 | 0.001 | 0.0005 | 0.00025 | MID2 |
| ZosteropsA00_RUNE1.bpp.ctl | 3 | 0.001 | 0.0005 | 0.004 | 0.002 | ROOT |
| ZosteropsA00_RUNE2.bpp.ctl | 3 | 0.001 | 0.0005 | 0.004 | 0.002 | ROOT |
| ZosteropsA00_RUNE3.bpp.ctl | 3 | 0.001 | 0.0005 | 0.004 | 0.002 | ROOT |
| ZosteropsA00_RUNE4.bpp.ctl | 3 | 0.002 | 0.001 | 0.004 | 0.002 | ROOT |
| ZosteropsA00_RUNE5.bpp.ctl | 3 | 0.002 | 0.001 | 0.004 | 0.002 | ROOT |
| ZosteropsA00_RUNE6.bpp.ctl | 3 | 0.002 | 0.001 | 0.004 | 0.002 | ROOT |
| ZosteropsA00_RUNE7.bpp.ctl | 3 | 0.001 | 0.0005 | 0.0005 | 0.00025 | ROOT |
| ZosteropsA00_RUNE8.bpp.ctl | 3 | 0.001 | 0.0005 | 0.0005 | 0.00025 | ROOT |

| | | | | | | |
|---|---|---|---|---|---|---|
| ZosteropsA00_RUNE9.bpp.ctl | 3 | 0.001 | 0.0005 | 0.0005 | 0.00025 | ROOT |
| ZosteropsA00_RUNE10.bpp.ctl | 3 | 0.002 | 0.001 | 0.0005 | 0.00025 | ROOT |
| ZosteropsA00_RUNE11.bpp.ctl | 3 | 0.002 | 0.001 | 0.0005 | 0.00025 | ROOT |
| ZosteropsA00_RUNE12.bpp.ctl | 3 | 0.002 | 0.001 | 0.0005 | 0.00025 | ROOT |

Supplementary table 4: Description of replicate runs of BPP for estimation of $\theta$ and $\tau$ for GDI calculations (A00).

| Museum | Catalog number | Locale | Taxon | Group | Culmen length (mm) | Nare-to-tip (mm) | Bill depth (mm) | Bill width (mm) | Tarsus (mm) | Wing cord (mm) | Tail length (mm) |
|---|---|---|---|---|---|---|---|---|---|---|---|
| American Museum of Natural History | 418684 | Shandong, China | *Z. erythropleurus* | erythropleurus | 9.2 | 6.4 | 2.9 | 2.8 | 17.2 | 59.5 | 41 |
| American Museum of Natural History | 418685 | Shandong, China | *Z. erythropleurus* | erythropleurus | 9.6 | 5.7 | 2.7 | 3.1 | 16.2 | 62 | 39 |
| American Museum of Natural History | 418686 | Shandong, China | *Z. erythropleurus* | erythropleurus | 9.1 | 6.3 | 2.5 | 2.7 | 15.9 | 51 | 42 |
| American Museum of Natural History | 418687 | Shandong, China | *Z. erythropleurus* | erythropleurus | 9.8 | 6.7 | 2.9 | 2.8 | 15.6 | 63.5 | 41 |
| American Museum of Natural History | 418688 | Shandong, China | *Z. erythropleurus* | erythropleurus | 8.1 | 5.8 | 2.5 | 2.8 | 14.8 | 57.5 | 41 |
| American Museum of Natural History | 418689 | Shandong, China | *Z. erythropleurus* | erythropleurus | 9.6 | 6.6 | 2.7 | 2.7 | 16.9 | 61 | 44 |
| American Museum of Natural History | 418690 | NOLOCALE | *Z. erythropleurus* | erythropleurus | 9.4 | 6.8 | 2.5 | 2.6 | 16 | 62.5 | 40 |
| American Museum of Natural History | 699704 | Yunnan, China | *Z. erythropleurus* | erythropleurus | 10.2 | 7.6 | 2.8 | 2.7 | 15.2 | 62 | 39 |
| American Museum of Natural History | 699705 | Yunnan, China | *Z. erythropleurus* | erythropleurus | 9.9 | 7.2 | 2.8 | 3 | 15.5 | 59 | 38 |
| American Museum of Natural History | 699706 | Yunnan, China | *Z. erythropleurus* | erythropleurus | 10 | 6.9 | 3.1 | 2.9 | 15.5 | 60 | 38 |
| American Museum of Natural History | 699707 | Yunnan, China | *Z. erythropleurus* | erythropleurus | 10 | 6.9 | 2.6 | 3 | 14.6 | 59 | 37 |
| American Museum of Natural History | 699708 | Yunnan, China | *Z. erythropleurus* | erythropleurus | 9.3 | 6.8 | 2.7 | 2.8 | 16.5 | 61.5 | 41 |
| American Museum of Natural History | 699709 | Yunnan, China | *Z. erythropleurus* | erythropleurus | 10 | 6.5 | 2.8 | 2.8 | 15.6 | 60.5 | 38 |
| American Museum of Natural History | 699710 | Shanghai, China | *Z. erythropleurus* | erythropleurus | 10.1 | 6.5 | 3.3 | 2.8 | 14.9 | 61.5 | 44 |
| American Museum of Natural History | 699711 | Beijing, China | *Z. erythropleurus* | erythropleurus | 9.1 | 6.6 | 2.6 | 2.6 | 16.8 | 59.5 | 40 |
| American Museum of Natural History | 699712 | Beijing, China | *Z. erythropleurus* | erythropleurus | 9.7 | 6.1 | 2.9 | 2.8 | 16.4 | 57.5 | 41 |
| American Museum of Natural History | 699713 | Beijing, China | *Z. erythropleurus* | erythropleurus | 9.7 | 6.3 | 2.9 | 2.9 | 18.9 | 54 | 37 |
| American Museum of Natural History | 699714 | Beijing, China | *Z. erythropleurus* | erythropleurus | 9.4 | 7 | 3.1 | 2.8 | 16.7 | 59.5 | 45 |
| American Museum of Natural History | 699715 | Beijing, China | *Z. erythropleurus* | erythropleurus | 9.1 | 6.2 | 2.5 | 2.4 | 18.3 | 59.5 | 40 |
| American Museum of Natural History | 699716 | Beijing, China | *Z. erythropleurus* | erythropleurus | 10.3 | 6.4 | 2.8 | 2.6 | 15 | 61.2 | 42 |
| American Museum of Natural History | 699738 | Iwoto, Japan | *Z. japonicus alani* | japonicus OCEANIC | 12.5 | 8.1 | 3.8 | 3.5 | 22.2 | 62 | 45 |
| American Museum of Natural History | 699739 | Iwoto, Japan | *Z. japonicus alani* | japonicus OCEANIC | 11.8 | 7.9 | 3.4 | 3.2 | 20.4 | 59 | 46 |
| American Museum of Natural History | 699740 | Iwoto, Japan | *Z. japonicus alani* | japonicus OCEANIC | 11.9 | 8.4 | 3.5 | 3.6 | 20.3 | 61.5 | 44 |
| American Museum of Natural History | 699742 | Iwoto, Japan | *Z. japonicus alani* | japonicus OCEANIC | 12.5 | 8.5 | 3.3 | 3.2 | 18.5 | 60.5 | 47 |
| American Museum of Natural History | 699743 | Iwoto, Japan | *Z. japonicus alani* | japonicus OCEANIC | 11.9 | 7.5 | 3.7 | 3.4 | 18.7 | 60 | 48 |
| American Museum of Natural History | 699744 | Iwoto, Japan | *Z. japonicus alani* | japonicus OCEANIC | 12.8 | 7.9 | 3.5 | 3.6 | 20.3 | 62 | 46 |

| | | | | | | | | | | | |
|---|---|---|---|---|---|---|---|---|---|---|---|
| American Museum of Natural History | 699745 | Iwoto, Japan | *Z. japonicus alani* | japonicus OCEANIC | 12.3 | 7.7 | 3.4 | 3.3 | 21.1 | 60.5 | 47 |
| American Museum of Natural History | 699746 | Iwoto, Japan | *Z. japonicus alani* | japonicus OCEANIC | 12.7 | 8.2 | 3.6 | 3.3 | 20.2 | 61 | 44 |
| American Museum of Natural History | 785351 | Daito Islands, Japan | *Z. japonicus daitoensis* | japonicus OCEANIC | 10.7 | 7.6 | 3.1 | 2.9 | 17.2 | 61 | 41 |
| American Museum of Natural History | 699764 | Yakushima, Japan | *Z. japonicus insularis* | japonicus CONTINENTAL | 12.5 | 8.3 | 2.8 | 3.2 | 22.1 | 60.5 | 50 |
| American Museum of Natural History | 699786 | Tanegashima, Japan | *Z. japonicus insularis* | japonicus CONTINENTAL | 11.4 | 7.9 | 3.1 | 3.1 | 17.7 | 58 | 46 |
| American Museum of Natural History | 785303 | South Korea | *Z. japonicus japonicus* | japonicus CONTINENTAL | 12 | 8.1 | 2.8 | 2.8 | 18.6 | 59.5 | 43 |
| American Museum of Natural History | 785340 | Honshu, Japan | *Z. japonicus japonicus* | japonicus CONTINENTAL | 10.8 | 7.7 | 2.6 | 2.5 | 17.4 | 58.5 | 43 |
| American Museum of Natural History | 839406 | Tsushima, Japan | *Z. japonicus japonicus* | japonicus CONTINENTAL | 11.7 | 8.3 | 3 | 3 | 17.9 | 60 | 45 |
| American Museum of Natural History | 839413 | Tsushima, Japan | *Z. japonicus japonicus* | japonicus CONTINENTAL | 10.4 | 7.3 | 3 | 2.5 | 17.5 | 58 | 43 |
| American Museum of Natural History | 839425 | Tsushima, Japan | *Z. japonicus japonicus* | japonicus CONTINENTAL | 12.8 | 8 | 3.2 | 3.1 | 17.1 | 61.5 | 49 |
| American Museum of Natural History | 839434 | Tsushima, Japan | *Z. japonicus japonicus* | japonicus CONTINENTAL | 11.3 | 8.5 | 3.1 | 3.2 | 18.6 | 60 | 42 |
| American Museum of Natural History | 699789 | Ishigaki, Japan | *Z. japonicus loochooensis* | japonicus CONTINENTAL | 11.4 | 7.9 | 2.9 | 3.1 | 18.1 | 58 | 45 |
| American Museum of Natural History | 699809 | Amami, Japan | *Z. japonicus loochooensis* | japonicus CONTINENTAL | 10.6 | 7.8 | 2.8 | 2.4 | 17.9 | 57 | 45 |
| American Museum of Natural History | 699815 | Amami, Japan | *Z. japonicus loochooensis* | japonicus CONTINENTAL | 11 | 7.7 | 3 | 3.1 | 18.2 | 57.5 | 44 |
| American Museum of Natural History | 699816 | Amami, Japan | *Z. japonicus loochooensis* | japonicus CONTINENTAL | 10 | 7.2 | 2.8 | 3 | 19.6 | 57 | 41 |
| American Museum of Natural History | 699819 | Amami, Japan | *Z. japonicus loochooensis* | japonicus CONTINENTAL | 10.2 | 7.6 | 2.7 | 2.8 | 17.1 | 55 | 48 |
| American Museum of Natural History | 699823 | Amami, Japan | *Z. japonicus loochooensis* | japonicus CONTINENTAL | 10.6 | 7.4 | 3 | 3.1 | 19.3 | 55 | 43 |
| American Museum of Natural History | 699826 | Amami, Japan | *Z. japonicus loochooensis* | japonicus CONTINENTAL | 10.9 | 7.2 | 2.8 | 3 | 17.7 | 55.5 | 45 |
| American Museum of Natural History | 461439 | Miyakejima, Japan | *Z. japonicus stejnegeri* | japonicus OCEANIC | 14.6 | 9.6 | 3.2 | 3.3 | 19.6 | 63.5 | 50 |
| American Museum of Natural History | 461440 | Hachijojima, Japan | *Z. japonicus stejnegeri* | japonicus OCEANIC | 13.6 | 9.9 | 3.1 | 3.2 | 22.6 | 62 | 47 |
| American Museum of Natural History | 703551 | Miyakejima, Japan | *Z. japonicus stejnegeri* | japonicus OCEANIC | 14.1 | 10.1 | 3.6 | 3.5 | 20.1 | 63 | 48 |
| American Museum of Natural History | 703552 | Miyakejima, Japan | *Z. japonicus stejnegeri* | japonicus OCEANIC | 13.6 | 9.7 | 3.3 | 3.4 | 19.3 | 63 | 48 |
| American Museum of Natural History | 785311 | Izu Islands, Japan | *Z. japonicus stejnegeri* | japonicus OCEANIC | 13.4 | 9.6 | 3.7 | 3.7 | 18.8 | 61 | 40 |
| American Museum of Natural History | 785314 | Izu Islands, Japan | *Z. japonicus stejnegeri* | japonicus OCEANIC | 10.3 | 7.9 | 2.8 | 2.8 | 19.5 | 60.5 | 44 |
| American Museum of Natural History | 785316 | Izu Islands, Japan | *Z. japonicus stejnegeri* | japonicus OCEANIC | 15.5 | 10.7 | 3.3 | 3.3 | 19.6 | 62 | 42 |
| American Museum of Natural History | 785324 | Izu Islands, Japan | *Z. japonicus stejnegeri* | japonicus OCEANIC | 14.4 | 10.1 | 3.2 | 3.2 | 21.2 | 60 | 46 |
| American Museum of Natural History | 810380 | Lanyu Island, Taiwan | *Z. meyeni batanis* | meyeni | 10.6 | 7.6 | 3.2 | 3.3 | 18.5 | 54.5 | 43 |
| American Museum of Natural History | 839394 | Lanyu Island, Taiwan | *Z. meyeni batanis* | meyeni | 11.4 | 8.1 | 3.5 | 3.9 | 17.4 | 59.5 | 41 |

| | | | | | | | | | | | |
|---|---|---|---|---|---|---|---|---|---|---|---|
| American Museum of Natural History | 839395 | Lanyu Island, Taiwan | *Z. meyeni batanis* | meyeni | 10.2 | 7.2 | 3 | 3.2 | 17.6 | 60 | 45 |
| American Museum of Natural History | 839396 | Lanyu Island, Taiwan | *Z. meyeni batanis* | meyeni | 10.7 | 7.6 | 3.2 | 3.4 | 17.9 | 58 | 41 |
| American Museum of Natural History | 839397 | Lanyu Island, Taiwan | *Z. meyeni batanis* | meyeni | 11.1 | 7.9 | 3 | 3.5 | 18.4 | 56 | 38 |
| American Museum of Natural History | 839397 | Lanyu Island, Taiwan | *Z. meyeni batanis* | meyeni | 10.3 | 7.9 | 2.9 | 3.3 | 19.3 | 55 | 37 |
| American Museum of Natural History | 839399 | Lanyu Island, Taiwan | *Z. meyeni batanis* | meyeni | 9.9 | 7.7 | 3 | 3.5 | 17.6 | 56.5 | 39 |
| American Museum of Natural History | 839402 | Lanyu Island, Taiwan | *Z. meyeni batanis* | meyeni | 10.3 | 7.4 | 3.2 | 3.2 | 17.9 | 55 | 38 |
| American Museum of Natural History | 346385 | Timor, Indonesia | *Z. montanus montanus* | montanus SOUTH | 9.7 | 6.9 | 3 | 3 | 15.4 | 59.5 | 42 |
| American Museum of Natural History | 346386 | Timor, Indonesia | *Z. montanus montanus* | montanus SOUTH | 9.3 | 6.9 | 2.8 | 2.8 | 17.1 | 59.5 | 44 |
| American Museum of Natural History | 346390 | Timor, Indonesia | *Z. montanus montanus* | montanus SOUTH | 9.7 | 7.1 | 2.7 | 3 | 15.7 | 56 | 42 |
| American Museum of Natural History | 459968 | Negros, Philippines | *Z. montanus pectoralis* | montanus SOUTH | 10.3 | 7.4 | 3.2 | 2.6 | 18.8 | 57 | 42 |
| American Museum of Natural History | 488892 | Negros, Philippines | *Z. montanus pectoralis* | montanus SOUTH | 9.3 | 7.2 | 3 | 2.6 | 17.1 | 52.5 | 37 |
| American Museum of Natural History | 488896 | Negros, Philippines | *Z. montanus pectoralis* | montanus SOUTH | 9.8 | 7.1 | 2.8 | 3 | 17 | 54.5 | 40 |
| American Museum of Natural History | 700146 | Mindanao, Philippines | *Z. montanus vulcani* | montanus SOUTH | 9.9 | 7.1 | 3.2 | 3.5 | 15.5 | 55.5 | 43 |
| American Museum of Natural History | 416949 | Luzon, Philippines | *Z. montanus whiteheadi* | montanus NORTH | 9.9 | 7.1 | 3.1 | 2.7 | 16.4 | 55 | 40 |
| American Museum of Natural History | 115079 | Shandong, China | *Z. simplex simplex* | simplex | 8 | 6.3 | 2.7 | 2.8 | 15.8 | 59 | 43 |
| American Museum of Natural History | 115084 | Shandong, China | *Z. simplex simplex* | simplex | 9.8 | 7.4 | 2.7 | 2.6 | 15.2 | 58 | 43 |
| American Museum of Natural History | 115085 | NOLOCALE | *Z. simplex simplex* | simplex | 8.7 | 7.1 | 2.7 | 2.9 | 16.3 | 57 | 39 |
| American Museum of Natural History | 204017 | NOLOCALE | *Z. simplex simplex* | simplex | 10 | 7.1 | 2.7 | 2.4 | 16 | 55 | 46 |
| American Museum of Natural History | 418694 | Shandong, China | *Z. simplex simplex* | simplex | 9.4 | 6.6 | 2.4 | 2.7 | 15.4 | 60 | 39 |
| American Museum of Natural History | 418701 | Guangdong, China | *Z. simplex simplex* | simplex | 9.1 | 6.8 | 3 | 2.8 | 16.1 | 55 | 39 |
| American Museum of Natural History | 699854 | Yunnan, China | *Z. simplex simplex* | simplex | 9.6 | 7 | 2.7 | 2.9 | 17.5 | 56 | 44 |
| American Museum of Natural History | 699856 | Yunnan, China | *Z. simplex simplex* | simplex | 9.6 | 6.7 | 2.5 | 2.9 | 16.9 | 56 | 38 |
| American Museum of Natural History | 699876 | Xiamen, China | *Z. simplex simplex* | simplex | 10 | 7.6 | 2.9 | 3.2 | 15 | 55.5 | 43 |
| American Museum of Natural History | 699889 | Jiujiang, China | *Z. simplex simplex* | simplex | 8.8 | 6.4 | 2.8 | 2.9 | 16.1 | 55.5 | 44 |
| American Museum of Natural History | 699906 | Taiwan | *Z. simplex simplex* | simplex | 9.1 | 7 | 3 | 3 | 16.1 | 50.5 | 35 |
| American Museum of Natural History | 699916 | Taiwan | *Z. simplex simplex* | simplex | 9.9 | 7.1 | 2.5 | 3.1 | 16.7 | 51 | 37 |
| American Museum of Natural History | 699918 | Taiwan | *Z. simplex simplex* | simplex | 9.2 | 6.9 | 2.6 | 2.7 | 16.4 | 54.5 | 41 |
| American Museum of Natural History | 699919 | Taiwan | *Z. simplex simplex* | simplex | 10.9 | 7.5 | 2.9 | 2.8 | 16.7 | 53 | 40 |

| | | | | | | | | | | | |
|---|---|---|---|---|---|---|---|---|---|---|---|
| American Museum of Natural History | 699923 | Taiwan | *Z. simplex simplex* | simplex | 9 | 7.2 | 2.7 | 2.9 | 15.9 | 54 | 42 |
| American Museum of Natural History | 785346 | Taiwan | *Z. simplex simplex* | simplex | 9.6 | 7.3 | 2.7 | 3.1 | 15.8 | 51.5 | 37 |
| Cincinnati Museum Center | B35473 | Mindanao, Philippines | *Z. everetti basilanicus* | everetti | 11.1 | 7.9 | 3.55 | 3.58 | 16.19 | 55.5 | 34 |
| Cincinnati Museum Center | B35474 | Mindanao, Philippines | *Z. everetti basilanicus* | everetti | 10.25 | 7.34 | 3.05 | 3.07 | 16.81 | 56 | 40 |
| Cincinnati Museum Center | B35475 | Mindanao, Philippines | *Z. everetti basilanicus* | everetti | 10.47 | 7.53 | 3.53 | 3.69 | 17.44 | 56 | 37 |
| Cincinnati Museum Center | B35476 | Mindanao, Philippines | *Z. everetti basilanicus* | everetti | 11.73 | 8.45 | 3.04 | 3.79 | 17.43 | 56 | 37 |
| Cincinnati Museum Center | B33961 | Biliran, Philippines | *Z. everetti boholensis* | everetti | 10.05 | 7.39 | 4.39 | 3.57 | 18.52 | 55 | 38 |
| Cincinnati Museum Center | B33359 | Batan, Philippines | *Z. meyeni batanis* | meyeni | 11.18 | 8.84 | 3.61 | 3.54 | 17.3 | 58 | 38 |
| Cincinnati Museum Center | B36979 | Mindoro, Philippines | *Z. montanus halconensis* | montanus NORTH | 10.85 | 7.82 | 2.94 | 3.15 | 17.29 | 53 | 36 |
| Cincinnati Museum Center | B36991 | Mindoro, Philippines | *Z. montanus halconensis* | montanus NORTH | 10.76 | 7.81 | 2.74 | 3.4 | 16.74 | 55 | 35 |
| Cincinnati Museum Center | B37023 | Negros, Philippines | *Z. montanus pectoralis* | montanus SOUTH | 11.03 | 8.57 | 3.22 | 3.57 | 18.84 | 58 | 38 |
| Cincinnati Museum Center | B37041 | Negros, Philippines | *Z. montanus pectoralis* | montanus SOUTH | 11.23 | 8.17 | 3.17 | 3.23 | 19.28 | 59 | 38 |
| Cincinnati Museum Center | B37042 | Negros, Philippines | *Z. montanus pectoralis* | montanus SOUTH | 10.42 | 7.99 | 2.81 | 3.19 | 17.06 | 56.5 | 35 |
| Cincinnati Museum Center | B37043 | Negros, Philippines | *Z. montanus pectoralis* | montanus SOUTH | 10.59 | 8.23 | 3.15 | 3.71 | 17.54 | 58 | 36 |
| Cincinnati Museum Center | B37052 | Negros, Philippines | *Z. montanus pectoralis* | montanus SOUTH | 11.05 | 8.38 | 3.15 | 3.51 | 17.79 | 60 | 40 |
| Cincinnati Museum Center | B37053 | Negros, Philippines | *Z. montanus pectoralis* | montanus SOUTH | 11.24 | 8.65 | 3.23 | 3.24 | 17.94 | 61.5 | 37 |
| Cincinnati Museum Center | B37054 | Negros, Philippines | *Z. montanus pectoralis* | montanus SOUTH | 10.9 | 8.15 | 2.88 | 3.46 | 17.54 | 60.5 | 38 |
| Cincinnati Museum Center | B37073 | Negros, Philippines | *Z. montanus pectoralis* | montanus SOUTH | 11.22 | 8.05 | 3.23 | 3.64 | 17.39 | 61.5 | 39 |
| Cincinnati Museum Center | B36817 | Panay, Philippines | *Z. montanus ssp* | montanus SOUTH | 11.08 | 8.03 | 3.03 | 3.72 | 17.41 | 54 | 37 |
| Cincinnati Museum Center | B36818 | Panay, Philippines | *Z. montanus ssp* | montanus SOUTH | 11.77 | 8.7 | 3.6 | 3.8 | 16.34 | 55 | 35 |
| Cincinnati Museum Center | B36819 | Panay, Philippines | *Z. montanus ssp* | montanus SOUTH | 11.39 | 8.38 | 3.14 | 3.31 | 17.24 | 56 | 37 |
| Cincinnati Museum Center | B36894 | Panay, Philippines | *Z. montanus ssp* | montanus SOUTH | 11.34 | 8.3 | 3.2 | 3.26 | 19.02 | 55 | 35 |
| Cincinnati Museum Center | B36895 | Panay, Philippines | *Z. montanus ssp* | montanus SOUTH | 11.44 | 8.48 | 3.28 | 3.08 | 19.16 | 53 | 34 |
| Cincinnati Museum Center | B36896 | Panay, Philippines | *Z. montanus ssp* | montanus SOUTH | 11.37 | 8.63 | 3.01 | 3.28 | 17.72 | 54 | 35 |
| Cincinnati Museum Center | B36897 | Panay, Philippines | *Z. montanus ssp* | montanus SOUTH | 11.55 | 8.36 | 3.46 | 3.49 | 17.48 | 51 | 35 |
| Cincinnati Museum Center | B36898 | Panay, Philippines | *Z. montanus ssp* | montanus SOUTH | 11.3 | 8.09 | 3.16 | 3.22 | 16.7 | 54 | 36 |
| Cincinnati Museum Center | B35538 | Mindanao, Philippines | *Z. montanus vulcani* | montanus SOUTH | 9.01 | 6.63 | 3.15 | 3.48 | 17.9 | 52 | 33 |
| Cincinnati Museum Center | B35539 | Mindanao, Philippines | *Z. montanus vulcani* | montanus SOUTH | 10.69 | 8.54 | 3.22 | 2.87 | 16.26 | 55.5 | 36 |

| | | | | | | | | | | | |
|---|---|---|---|---|---|---|---|---|---|---|---|
| Cincinnati Museum Center | B35540 | Mindanao, Philippines | *Z. montanus vulcani* | montanus SOUTH | 9.43 | 7.21 | 3.13 | 3.48 | 15.71 | 53 | 34 |
| Cincinnati Museum Center | B35541 | Mindanao, Philippines | *Z. montanus vulcani* | montanus SOUTH | 9.56 | 7.13 | 3.64 | 3.54 | 16.43 | 52 | 32 |
| Cincinnati Museum Center | B35543 | Mindanao, Philippines | *Z. montanus vulcani* | montanus SOUTH | 10.29 | 7.48 | 2.82 | 3.22 | 15.76 | 55 | 38 |
| Cincinnati Museum Center | B35626 | Mindanao, Philippines | *Z. montanus vulcani* | montanus SOUTH | 9.92 | 7.76 | 3.26 | 2.93 | 17.34 | 52 | 38 |
| Cincinnati Museum Center | B35836 | Mindanao, Philippines | *Z. montanus vulcani* | montanus SOUTH | 10.08 | 7.28 | 2.91 | 3.58 | 17.95 | 58.5 | 34 |
| Cincinnati Museum Center | B35837 | Mindanao, Philippines | *Z. montanus vulcani* | montanus SOUTH | 9.95 | 7.58 | 3.1 | 3.79 | 17.15 | 58.5 | 29 |
| Cincinnati Museum Center | B35838 | Mindanao, Philippines | *Z. montanus vulcani* | montanus SOUTH | 10.92 | 7.97 | 2.78 | 3.24 | 15.86 | 58 | 35 |
| Cincinnati Museum Center | B36301 | Mindanao, Philippines | *Z. montanus vulcani* | montanus SOUTH | 10.69 | 7.21 | 3.06 | 3.45 | 18.15 | 54 | 34 |
| Cincinnati Museum Center | B36302 | Mindanao, Philippines | *Z. montanus vulcani* | montanus SOUTH | 10.97 | 7.71 | 2.99 | 3.41 | 16.24 | 53.5 | 33 |
| Cincinnati Museum Center | B37376 | Mindanao, Philippines | *Z. montanus vulcani* | montanus SOUTH | 10.66 | 7.68 | 3.39 | 3.53 | 20.94 | 57.5 | 36 |
| Cincinnati Museum Center | B37674 | Mindanao, Philippines | *Z. montanus vulcani* | montanus SOUTH | 10.44 | 7.25 | 3.18 | 2.96 | 17.55 | 57 | 35 |
| Cincinnati Museum Center | B37675 | Mindanao, Philippines | *Z. montanus vulcani* | montanus SOUTH | 11.04 | 7.7 | 3.06 | 3.31 | 18.19 | 55 | 34 |
| Cincinnati Museum Center | B37676 | Mindanao, Philippines | *Z. montanus vulcani* | montanus SOUTH | 10.13 | 7.66 | 2.95 | 3.21 | 18.22 | 57 | 35 |
| Cincinnati Museum Center | B38163 | Mindanao, Philippines | *Z. montanus vulcani* | montanus SOUTH | 11.11 | 8.58 | 3.24 | 3.76 | 16.74 | 59 | 36 |
| Cincinnati Museum Center | B39029 | Mindanao, Philippines | *Z. montanus vulcani* | montanus SOUTH | 10.79 | 7.77 | 3 | 3.09 | 17.2 | 61.5 | 36 |
| Cincinnati Museum Center | B39198 | Mindanao, Philippines | *Z. montanus vulcani* | montanus SOUTH | 10.16 | 7.71 | 3.13 | 3.6 | 17.17 | 54.5 | 34 |
| Cincinnati Museum Center | B39200 | Mindanao, Philippines | *Z. montanus vulcani* | montanus SOUTH | 9.86 | 7.22 | 3.61 | 3.3 | 16.28 | 57 | 33 |
| Cincinnati Museum Center | B39201 | Mindanao, Philippines | *Z. montanus vulcani* | montanus SOUTH | 10.63 | 7.59 | 3.14 | 3.49 | 18.52 | 56 | 35 |
| Cincinnati Museum Center | B36615 | Luzon, Philippines | *Z. montanus whiteheadi* | montanus NORTH | 11.28 | 8.25 | 3.16 | 3.43 | 18.47 | 55 | 37 |
| Cincinnati Museum Center | B36616 | Luzon, Philippines | *Z. montanus whiteheadi* | montanus NORTH | 10.66 | 7.72 | 2.89 | 3.37 | 17.68 | 56 | 36 |
| Cincinnati Museum Center | B36617 | Luzon, Philippines | *Z. montanus whiteheadi* | montanus NORTH | 11.27 | 8.67 | 3.03 | 3.76 | 17.37 | 56 | 36 |
| Cincinnati Museum Center | B38226 | Luzon, Philippines | *Z. nigrorum innominatus* | nigrorum | 8.96 | 6.67 | 2.57 | 2.8 | 15.89 | 51 | 34 |
| Cincinnati Museum Center | B38227 | Luzon, Philippines | *Z. nigrorum innominatus* | nigrorum | 8.77 | 6.83 | 2.74 | 2.92 | 16.47 | 49.5 | 32 |
| Cincinnati Museum Center | B38228 | Luzon, Philippines | *Z. nigrorum innominatus* | nigrorum | 8.71 | 6.42 | 2.71 | 2.54 | 16.42 | 49 | 34 |
| Cincinnati Museum Center | B38321 | Luzon, Philippines | *Z. nigrorum innominatus* | nigrorum | 9.82 | 7.22 | 3.17 | 3 | 15.44 | 53 | 34 |
| Cincinnati Museum Center | B38322 | Luzon, Philippines | *Z. nigrorum innominatus* | nigrorum | 9.29 | 6.63 | 2.87 | 2.95 | 15.8 | 50 | 35 |
| Cincinnati Museum Center | B36816 | Panay, Philippines | *Z. nigrorum nigrorum* | nigrorum | 9.95 | 6.76 | 2.76 | 2.76 | 16.52 | 54 | 37 |
| Field Museum | 227802 | Mindanao, Philippines | *Z. everetti basilanicus* | everetti | 10.6 | 9 | 3.4 | 3.5 | 16.8 | 58.5 | 42 |

| | | | | | | | | | | | |
|---|---|---|---|---|---|---|---|---|---|---|---|
| Field Museum | 275478 | Mindanao, Philippines | *Z. everetti basilanicus* | everetti | 10.9 | 7.6 | 3.1 | 3.4 | 16 | 53.5 | 33 |
| Field Museum | 275481 | Mindanao, Philippines | *Z. everetti basilanicus* | everetti | 9.8 | 7.1 | 3 | 3 | 16.3 | 56.5 | 45 |
| Field Museum | 279653 | Mindanao, Philippines | *Z. everetti basilanicus* | everetti | 9.4 | 7 | 3.3 | 3.4 | 16.9 | 54 | 38 |
| Field Museum | 279658 | Mindanao, Philippines | *Z. everetti basilanicus* | everetti | 10.1 | 8.3 | 3.5 | 3.4 | 17.6 | 56.5 | 43 |
| Field Museum | 279662 | Mindanao, Philippines | *Z. everetti basilanicus* | everetti | 9.7 | 7.5 | 3.4 | 3.1 | 19.3 | 55 | 43 |
| Field Museum | 279667 | Mindanao, Philippines | *Z. everetti basilanicus* | everetti | 10.2 | 7.2 | 3 | 3.1 | 18.5 | 51 | 36 |
| Field Museum | 284302 | Mindanao, Philippines | *Z. everetti basilanicus* | everetti | 9.7 | 8.2 | 3.6 | 3.6 | 19.6 | 54.5 | 46 |
| Field Museum | 284635 | Camiguin, Philippines | *Z. everetti basilanicus* | everetti | 11.2 | 8.4 | 3.6 | 3.4 | 17.9 | 58.5 | 42 |
| Field Museum | 223705 | Bohol, Philippines | *Z. everetti boholensis* | everetti | 10.1 | 8 | 3.3 | 3 | 21.1 | 55 | 41 |
| Field Museum | 223707 | Bohol, Philippines | *Z. everetti boholensis* | everetti | 10.4 | 7.9 | 3.5 | 3.3 | 19.6 | 55 | 42 |
| Field Museum | 223709 | Bohol, Philippines | *Z. everetti boholensis* | everetti | 9.7 | 8.4 | 3.2 | 3.3 | 17.9 | 54.5 | 44 |
| Field Museum | 223710 | Bohol, Philippines | *Z. everetti boholensis* | everetti | 9.9 | 7.6 | 3.1 | 3.3 | 18.3 | 55 | 38 |
| Field Museum | 223713 | Bohol, Philippines | *Z. everetti boholensis* | everetti | 9.7 | 7.3 | 3.2 | 3.1 | 17.4 | 57.5 | 43 |
| Field Museum | 223718 | Bohol, Philippines | *Z. everetti boholensis* | everetti | 9.6 | 7.6 | 3.3 | 3.1 | 17.4 | 53.5 | 39 |
| Field Museum | 223719 | Bohol, Philippines | *Z. everetti boholensis* | everetti | 10.5 | 7.7 | 3.3 | 3 | 17.9 | 57 | 40 |
| Field Museum | 223720 | Bohol, Philippines | *Z. everetti boholensis* | everetti | 10 | 8.1 | 3.3 | 3.2 | 16.6 | 53.5 | 41 |
| Field Museum | 223721 | Bohol, Philippines | *Z. everetti boholensis* | everetti | 10 | 7.7 | 3.4 | 3.6 | 17.9 | 57.5 | 41 |
| Field Museum | 248393 | Samar, Philippines | *Z. everetti boholensis* | everetti | 10.6 | 7.7 | 3.5 | 3.2 | 16.8 | 56 | 41 |
| Field Museum | 248395 | Samar, Philippines | *Z. everetti boholensis* | everetti | 10.6 | 7.8 | 3.3 | 3.3 | 18.1 | 62.5 | 41 |
| Field Museum | 248398 | Samar, Philippines | *Z. everetti boholensis* | everetti | 9.9 | 7.6 | 3.1 | 3.1 | 15.3 | 53 | 38 |
| Field Museum | 276657 | Leyte, Philippines | *Z. everetti boholensis* | everetti | 10.6 | 7.7 | 3.4 | 3.2 | 17.7 | 55 | 47 |
| Field Museum | 276661 | Leyte, Philippines | *Z. everetti boholensis* | everetti | 10.8 | 7.6 | 3.1 | 3.4 | 18.3 | 58 | 41 |
| Field Museum | 276664 | Leyte, Philippines | *Z. everetti boholensis* | everetti | 10.5 | 7.2 | 3.4 | 3.4 | 17 | 55.5 | 41 |
| Field Museum | 276667 | Leyte, Philippines | *Z. everetti boholensis* | everetti | 11.5 | 8.1 | 3.3 | 3.3 | 16.9 | 55.5 | 40 |
| Field Museum | 276668 | Leyte, Philippines | *Z. everetti boholensis* | everetti | 10.6 | 7.3 | 3.1 | 3.1 | 16.6 | 59 | 41 |
| Field Museum | 276671 | Leyte, Philippines | *Z. everetti boholensis* | everetti | 10.3 | 7 | 3.6 | 3.1 | 16.4 | 58 | 42 |
| Field Museum | 276674 | Leyte, Philippines | *Z. everetti boholensis* | everetti | 10.6 | 7 | 3.3 | 3.1 | 17.1 | 56 | 39 |
| Field Museum | 217590 | Siquijor, Philippines | *Z. everetti siquijorensis* | everetti | 9.9 | 8.3 | 3.3 | 2.9 | 15.9 | 50 | 38 |

| | | | | | | | | | | | |
|---|---|---|---|---|---|---|---|---|---|---|---|
| Field Museum | 217593 | Siquijor, Philippines | *Z. everetti siquijorensis* | everetti | 9.2 | 7 | 2.9 | 3 | 17.3 | 53 | 37 |
| Field Museum | 219261 | Siquijor, Philippines | *Z. everetti siquijorensis* | everetti | 9.7 | 7.3 | 3.2 | 3.3 | 16.9 | 57 | 43 |
| Field Museum | 219262 | Siquijor, Philippines | *Z. everetti siquijorensis* | everetti | 10.5 | 8 | 3.2 | 3.3 | 17.5 | 54.5 | 43 |
| Field Museum | 222859 | Siquijor, Philippines | *Z. everetti siquijorensis* | everetti | 10.6 | 8.3 | 3.4 | 3.6 | 17.8 | 57 | 37 |
| Field Museum | 74153 | Iwoto, Japan | *Z. japonicus alani* | japonicus OCEANIC | 11.8 | 8.7 | 3.5 | 3 | 21.9 | 61 | 50 |
| Field Museum | 74154 | Minamitorishima, Japan | *Z. japonicus alani* | japonicus OCEANIC | 12.5 | 8.6 | 3.7 | 3.5 | 20.6 | 61 | 42 |
| Field Museum | 65667 | Honshu, Japan | *Z. japonicus japonicus* | japonicus CONTINENTAL | 11 | 7.8 | 2.6 | 3.1 | 19.3 | 58 | 44 |
| Field Museum | 74150 | Honshu, Japan | *Z. japonicus japonicus* | japonicus CONTINENTAL | 11.7 | 8.1 | 2.9 | 3 | 19.5 | 60 | 44 |
| Field Museum | 74151 | Honshu, Japan | *Z. japonicus japonicus* | japonicus CONTINENTAL | 11 | 7.1 | 2.7 | 3 | 20.5 | 59 | 45 |
| Field Museum | 257081 | South Korea | *Z. japonicus japonicus* | japonicus CONTINENTAL | 12.3 | 7.5 | 3 | 2.9 | 18.2 | 58 | 45 |
| Field Museum | 257082 | South Korea | *Z. japonicus japonicus* | japonicus CONTINENTAL | 12.1 | 7.9 | 3.4 | 2.8 | 20.4 | 60 | 42 |
| Field Museum | 65661 | Niijima, Japan | *Z. japonicus stejnegeri* | japonicus OCEANIC | 13.2 | 9.5 | 3.7 | 3.4 | 20 | 63 | 44 |
| Field Museum | 74152 | Miyakejima, Japan | *Z. japonicus stejnegeri* | japonicus OCEANIC | 12.7 | 8.8 | 3.3 | 3.3 | 19.3 | 61.5 | 43 |
| Field Museum | 246943 | Oshima, Japan | *Z. japonicus stejnegeri* | japonicus OCEANIC | 12.8 | 8.7 | 3.2 | 3.3 | 21.8 | 60 | 41 |
| Field Museum | 246944 | Oshima, Japan | *Z. japonicus stejnegeri* | japonicus OCEANIC | 13.1 | 9.4 | 2.9 | 3.2 | 20.7 | 62.5 | 49 |
| Field Museum | 246945 | Oshima, Japan | *Z. japonicus stejnegeri* | japonicus OCEANIC | 12.1 | 8.5 | 3.2 | 3.4 | 22.8 | 61.5 | 44 |
| Field Museum | 246946 | Oshima, Japan | *Z. japonicus stejnegeri* | japonicus OCEANIC | 13.8 | 9 | 3.4 | 3.5 | 20.9 | 61 | 42 |
| Field Museum | 246947 | Miyakejima, Japan | *Z. japonicus stejnegeri* | japonicus OCEANIC | 13 | 9.7 | 3.3 | 3.4 | 21.7 | 61 | 46 |
| Field Museum | 246948 | Miyakejima, Japan | *Z. japonicus stejnegeri* | japonicus OCEANIC | 14.2 | 10.1 | 3.4 | 3.5 | 23.2 | 64 | 45 |
| Field Museum | 246949 | Miyakejima, Japan | *Z. japonicus stejnegeri* | japonicus OCEANIC | 13.7 | 10.3 | 3.6 | 3.3 | 22 | 64 | 48 |
| Field Museum | 246950 | Miyakejima, Japan | *Z. japonicus stejnegeri* | japonicus OCEANIC | 13.9 | 9.3 | 3.1 | 4 | 21.5 | 62 | 43 |
| Field Museum | 186366 | Batan, Philippines | *Z. meyeni batanis* | meyeni | 11.2 | 8.7 | 3.5 | 3.5 | 18 | 55.5 | 41 |
| Field Museum | 186367 | Batan, Philippines | *Z. meyeni batanis* | meyeni | 11.2 | 7.9 | 3.4 | 3.3 | 19.7 | 58.5 | 44 |
| Field Museum | 219895 | Batan, Philippines | *Z. meyeni batanis* | meyeni | 11.2 | 8.2 | 3.5 | 3.4 | 19.4 | 56 | 42 |
| Field Museum | 219896 | Batan, Philippines | *Z. meyeni batanis* | meyeni | 10.8 | 8.4 | 3.3 | 4 | 18.3 | 59.5 | 44 |
| Field Museum | 20250 | Banton, Philippines | *Z. meyeni meyeni* | meyeni | 10.8 | 7.4 | 3.2 | 2.8 | 18.2 | 55.5 | 40 |
| Field Museum | 20749 | Banton, Philippines | *Z. meyeni meyeni* | meyeni | 11.4 | 8.8 | 3.2 | 3.8 | 19.5 | 56.5 | 43 |
| Field Museum | 24058 | Calayan, Philippines | *Z. meyeni meyeni* | meyeni | 10.6 | 7.8 | 3 | 3.4 | 18.9 | 55 | 41 |

| | | | | | | | | | | | |
|---|---|---|---|---|---|---|---|---|---|---|---|
| Field Museum | 24059 | Calayan, Philippines | *Z. meyeni meyeni* | meyeni | 10.6 | 8.3 | 3 | 3.1 | 20.6 | 55 | 38 |
| Field Museum | 187918 | Negros, Philippines | *Z. montanus pectoralis* | montanus SOUTH | 10.2 | 7.5 | 3.1 | 2.9 | 18.7 | 54.5 | 38 |
| Field Museum | 191359 | Negros, Philippines | *Z. montanus pectoralis* | montanus SOUTH | 9.9 | 7.8 | 3.1 | 3 | 17.6 | 55 | 32 |
| Field Museum | 217458 | Negros, Philippines | *Z. montanus pectoralis* | montanus SOUTH | 10.4 | 7.8 | 3 | 2.8 | 17.4 | 54 | 38 |
| Field Museum | 217460 | Negros, Philippines | *Z. montanus pectoralis* | montanus SOUTH | 9.5 | 7.4 | 3 | 2.3 | 17.1 | 56.5 | 39 |
| Field Museum | 219515 | Negros, Philippines | *Z. montanus pectoralis* | montanus SOUTH | 10 | 7.5 | 2.9 | 3.2 | 17.3 | 52 | 33 |
| Field Museum | 257249 | Negros, Philippines | *Z. montanus pectoralis* | montanus SOUTH | 11.7 | 8.8 | 3.1 | 3.1 | 18 | 59 | 42 |
| Field Museum | 257250 | Negros, Philippines | *Z. montanus pectoralis* | montanus SOUTH | 10.8 | 8.1 | 3.1 | 2.9 | 17.6 | 56 | 42 |
| Field Museum | 257252 | Negros, Philippines | *Z. montanus pectoralis* | montanus SOUTH | 11.8 | 8.6 | 3.6 | 3.6 | 17.9 | 56.5 | 42 |
| Field Museum | 257257 | Negros, Philippines | *Z. montanus pectoralis* | montanus SOUTH | 11.3 | 7.5 | 3.3 | 2.7 | 18 | 57.5 | 40 |
| Field Museum | 257259 | Negros, Philippines | *Z. montanus pectoralis* | montanus SOUTH | 11.2 | 7.9 | 3 | 3 | 17.4 | 58 | 42 |
| Field Museum | 257264 | Negros, Philippines | *Z. montanus pectoralis* | montanus SOUTH | 11.3 | 8 | 3.6 | 2.8 | 18.3 | 58 | 40 |
| Field Museum | 184500 | Mindanao, Philippines | *Z. montanus vulcani* | montanus SOUTH | 11.4 | 7.4 | 2.9 | 3 | 17 | 56 | 41 |
| Field Museum | 227704 | Mindanao, Philippines | *Z. montanus vulcani* | montanus SOUTH | 10.9 | 7.3 | 3.1 | 3.2 | 17.5 | 53.5 | 34 |
| Field Museum | 227708 | Mindanao, Philippines | *Z. montanus vulcani* | montanus SOUTH | 10 | 7.3 | 3.4 | 3.2 | 17.1 | 56 | 39 |
| Field Museum | 227716 | Mindanao, Philippines | *Z. montanus vulcani* | montanus SOUTH | 10.6 | 8.1 | 3.1 | 3 | 18.2 | 58.5 | 42 |
| Field Museum | 227717 | Mindanao, Philippines | *Z. montanus vulcani* | montanus SOUTH | 11 | 7.2 | 3.3 | 3.4 | 17.4 | 56 | 36 |
| Field Museum | 227719 | Mindanao, Philippines | *Z. montanus vulcani* | montanus SOUTH | 10.1 | 6.6 | 2.7 | 2.8 | 17.8 | 55 | 39 |
| Field Museum | 227722 | Mindanao, Philippines | *Z. montanus vulcani* | montanus SOUTH | 10.1 | 7.5 | 3.3 | 3.1 | 17.7 | 55.5 | 39 |
| Field Museum | 227729 | Mindanao, Philippines | *Z. montanus vulcani* | montanus SOUTH | 10.4 | 7.8 | 3.2 | 3.3 | 17.3 | 55.5 | 40 |
| Field Museum | 227731 | Mindanao, Philippines | *Z. montanus vulcani* | montanus SOUTH | 10.9 | 8 | 3 | 3.1 | 15.8 | 55 | 34 |
| Field Museum | 227732 | Mindanao, Philippines | *Z. montanus vulcani* | montanus SOUTH | 10.8 | 7.8 | 3 | 3.5 | 17.1 | 55.5 | 41 |
| Field Museum | 227742 | Mindanao, Philippines | *Z. montanus vulcani* | montanus SOUTH | 10.5 | 7.5 | 3.1 | 3.3 | 16.9 | 61 | 41 |
| Field Museum | 227744 | Mindanao, Philippines | *Z. montanus vulcani* | montanus SOUTH | 11.1 | 7.8 | 3.2 | 3.1 | 17.8 | 54 | 36 |
| Field Museum | 227762 | Mindanao, Philippines | *Z. montanus vulcani* | montanus SOUTH | 10.3 | 7.9 | 3.4 | 3.1 | 17 | 54.5 | 37 |
| Field Museum | 227770 | Mindanao, Philippines | *Z. montanus vulcani* | montanus SOUTH | 11.1 | 7.8 | 3 | 3.1 | 18.5 | 61 | 41 |
| Field Museum | 275462 | Mindanao, Philippines | *Z. montanus vulcani* | montanus SOUTH | 10.4 | 7.1 | 3.4 | 3.2 | 17.3 | 56.5 | 41 |
| Field Museum | 275471 | Mindanao, Philippines | *Z. montanus vulcani* | montanus SOUTH | 9.3 | 7.1 | 2.8 | 3.3 | 18.5 | 53 | 36 |

| | | | | | | | | | | | |
|---|---|---|---|---|---|---|---|---|---|---|---|
| Field Museum | 275472 | Mindanao, Philippines | *Z. montanus vulcani* | montanus SOUTH | 9.5 | 7.1 | 3.3 | 3.1 | 17.5 | 53.5 | 36 |
| Field Museum | 279633 | Mindanao, Philippines | *Z. montanus vulcani* | montanus SOUTH | 9.7 | 7.3 | 3.1 | 3.2 | 16.7 | 56.5 | 39 |
| Field Museum | 279637 | Mindanao, Philippines | *Z. montanus vulcani* | montanus SOUTH | 9.7 | 6.8 | 2.9 | 2.9 | 17.2 | 56 | 38 |
| Field Museum | 279640 | Mindanao, Philippines | *Z. montanus vulcani* | montanus SOUTH | 9.6 | 6.9 | 2.6 | 3.1 | 17.5 | 55 | 37 |
| Field Museum | 279646 | Mindanao, Philippines | *Z. montanus vulcani* | montanus SOUTH | 9.8 | 7.5 | 3 | 3.4 | 16.7 | 56 | 40 |
| Field Museum | 279649 | Mindanao, Philippines | *Z. montanus vulcani* | montanus SOUTH | 10.4 | 7.1 | 3.2 | 3.4 | 18.4 | 54 | 41 |
| Field Museum | 357634 | Mindanao, Philippines | *Z. montanus vulcani* | montanus SOUTH | 11.5 | 8.1 | 3.1 | 3.6 | 16.7 | 55.5 | 40 |
| Field Museum | 392321 | Mindanao, Philippines | *Z. montanus vulcani* | montanus SOUTH | 10.4 | 7.4 | 2.7 | 3.2 | 16.9 | 55 | 41 |
| Field Museum | 392322 | Mindanao, Philippines | *Z. montanus vulcani* | montanus SOUTH | 9.8 | 6.8 | 2.6 | 2.6 | 18.1 | 54.5 | 40 |
| Field Museum | 19746 | Luzon, Philippines | *Z. montanus whiteheadi* | montanus NORTH | 10 | 7 | 2.8 | 3 | 15.8 | 53 | 38 |
| Field Museum | 19747 | Luzon, Philippines | *Z. montanus whiteheadi* | montanus NORTH | 10.1 | 7 | 2.5 | 2.7 | 17.1 | 53 | 38 |
| Field Museum | 184493 | Luzon, Philippines | *Z. montanus whiteheadi* | montanus NORTH | 10.6 | 7.1 | 2.6 | 3 | 14.6 | 51 | 33 |
| Field Museum | 184494 | Luzon, Philippines | *Z. montanus whiteheadi* | montanus NORTH | 10.1 | 6.9 | 2.9 | 3.1 | 17.9 | 52 | 35 |
| Field Museum | 252895 | Luzon, Philippines | *Z. montanus whiteheadi* | montanus NORTH | 9.7 | 6.9 | 2.9 | 2.8 | 18.6 | 54.5 | 37 |
| Field Museum | 254259 | Luzon, Philippines | *Z. montanus whiteheadi* | montanus NORTH | 10.6 | 6.9 | 3.4 | 2.8 | 17.8 | 54 | 36 |
| Field Museum | 429299 | Luzon, Philippines | *Z. montanus whiteheadi* | montanus NORTH | 9.8 | 6.4 | 2.4 | 2.9 | 17.2 | 56 | 39 |
| Field Museum | 449785 | Luzon, Philippines | *Z. montanus whiteheadi* | montanus NORTH | 10.2 | 7.4 | 3 | 2.8 | 15.9 | 51 | 34 |
| Field Museum | 184515 | Luzon, Philippines | *Z. nigrorum aureiloris* | nigrorum | 9.3 | 6.3 | 2.4 | 2.2 | 16.7 | 51.5 | 38 |
| Field Museum | 184570 | Luzon, Philippines | *Z. nigrorum aureiloris* | nigrorum | 9.3 | 8.1 | 2.9 | 2.8 | 16.9 | 50.5 | 37 |
| Field Museum | 254260 | Luzon, Philippines | *Z. nigrorum aureiloris* | nigrorum | 9.9 | 6.9 | 3.1 | 3 | 16.7 | 52 | 37 |
| Field Museum | 254263 | Luzon, Philippines | *Z. nigrorum aureiloris* | nigrorum | 8.7 | 6.4 | 3.1 | 2.8 | 15.3 | 50.5 | 40 |
| Field Museum | 254270 | Luzon, Philippines | *Z. nigrorum aureiloris* | nigrorum | 10 | 7.1 | 3.1 | 2.7 | 17.6 | 52 | 38 |
| Field Museum | 254303 | Luzon, Philippines | *Z. nigrorum aureiloris* | nigrorum | 10 | 7.3 | 2.9 | 3.1 | 17 | 54 | 38 |
| Field Museum | 257406 | Luzon, Philippines | *Z. nigrorum aureiloris* | nigrorum | 9.5 | 6.8 | 3.2 | 2.7 | 17.8 | 53 | 37 |
| Field Museum | 257408 | Luzon, Philippines | *Z. nigrorum aureiloris* | nigrorum | 9.7 | 6.9 | 3 | 2.7 | 16.7 | 50 | 37 |
| Field Museum | 257414 | Luzon, Philippines | *Z. nigrorum aureiloris* | nigrorum | 9.1 | 7.2 | 3.4 | 2.5 | 17.1 | 53 | 35 |
| Field Museum | 284587 | Camiguin, Philippines | *Z. nigrorum catarmanensis* | nigrorum CS | 11.4 | 8.4 | 3.4 | 3.3 | 17.8 | 60 | 40 |
| Field Museum | 284592 | Camiguin, Philippines | *Z. nigrorum catarmanensis* | nigrorum CS | 11 | 8.3 | 3.6 | 3.2 | 20.1 | 58 | 40 |

| | | | | | | | | | | | |
|---|---|---|---|---|---|---|---|---|---|---|---|
| Field Museum | 284595 | Camiguin, Philippines | *Z. nigrorum catarmanensis* | nigrorum CS | 10.8 | 8.4 | 3.2 | 3.3 | 19.9 | 61 | 39 |
| Field Museum | 284609 | Camiguin, Philippines | *Z. nigrorum catarmanensis* | nigrorum CS | 10.4 | 7.9 | 3.1 | 3.6 | 19.9 | 59 | 43 |
| Field Museum | 284611 | Camiguin, Philippines | *Z. nigrorum catarmanensis* | nigrorum CS | 11.6 | 8 | 3.5 | 3.6 | 19.4 | 60 | 44 |
| Field Museum | 284612 | Camiguin, Philippines | *Z. nigrorum catarmanensis* | nigrorum CS | 11.2 | 8.2 | 3.4 | 3.1 | 19.2 | 63 | 45 |
| Field Museum | 284613 | Camiguin, Philippines | *Z. nigrorum catarmanensis* | nigrorum CS | 10.9 | 8 | 3.7 | 3.4 | 19.5 | 59 | 34 |
| Field Museum | 284615 | Camiguin, Philippines | *Z. nigrorum catarmanensis* | nigrorum CS | 11.4 | 8.4 | 3.2 | 3.4 | 18.4 | 56 | 41 |
| Field Museum | 284621 | Camiguin, Philippines | *Z. nigrorum catarmanensis* | nigrorum CS | 11.1 | 8.8 | 3.9 | 3.5 | 19.7 | 60 | 43 |
| Field Museum | 284622 | Camiguin, Philippines | *Z. nigrorum catarmanensis* | nigrorum CS | 11.1 | 8.2 | 3.6 | 3.5 | 18.5 | 61 | 42 |
| Field Museum | 286516 | Camiguin, Philippines | *Z. nigrorum catarmanensis* | nigrorum CS | 11.1 | 7.9 | 3.7 | 3.3 | 18.8 | 58 | 42 |
| Field Museum | 286521 | Camiguin, Philippines | *Z. nigrorum catarmanensis* | nigrorum CS | 11.1 | 7.7 | 3.8 | 3.3 | 18.8 | 57.5 | 38 |
| Field Museum | 286523 | Camiguin, Philippines | *Z. nigrorum catarmanensis* | nigrorum CS | 10.8 | 8.3 | 3.8 | 3.6 | 18.6 | 59 | 39 |
| Field Museum | 259878 | Luzon, Philippines | *Z. nigrorum innominatus* | nigrorum | 8.9 | 7.5 | 2.9 | 2.8 | 17.5 | 53 | 38 |
| Field Museum | 259893 | Luzon, Philippines | *Z. nigrorum innominatus* | nigrorum | 10.1 | 7.9 | 3.1 | 2.9 | 18.3 | 52.5 | 36 |
| Field Museum | 259894 | Luzon, Philippines | *Z. nigrorum innominatus* | nigrorum | 9.3 | 7.1 | 2.8 | 2.7 | 16.8 | 49.5 | 41 |
| Field Museum | 259899 | Luzon, Philippines | *Z. nigrorum innominatus* | nigrorum | 10.3 | 7.2 | 2.9 | 3 | 17.7 | 52 | 35 |
| Field Museum | 266569 | Luzon, Philippines | *Z. nigrorum luzonicus* | nigrorum | 10 | 7.3 | 3.4 | 3.1 | 16.3 | 56 | 41 |
| Field Museum | 266570 | Luzon, Philippines | *Z. nigrorum luzonicus* | nigrorum | 10.3 | 7.7 | 3.3 | 3.2 | 17 | 49 | 36 |
| Field Museum | 266571 | Luzon, Philippines | *Z. nigrorum luzonicus* | nigrorum | 9.9 | 7.7 | 3.1 | 3 | 17.4 | 53 | 39 |
| Field Museum | 266577 | Luzon, Philippines | *Z. nigrorum luzonicus* | nigrorum | 9.2 | 7.2 | 3.1 | 2.8 | 17.9 | 51.5 | 41 |
| Field Museum | 20060 | Ticao, Philippines | *Z. nigrorum nigrorum* | nigrorum | 9.5 | 7.8 | 3 | 3.3 | 19.2 | 55.5 | 41 |
| Field Museum | 20251 | Ticao, Philippines | *Z. nigrorum nigrorum* | nigrorum | 10.1 | 8.7 | 2.9 | 3 | 16.6 | 56 | 34 |
| Field Museum | 191341 | Negros, Philippines | *Z. nigrorum nigrorum* | nigrorum | 10.2 | 7.4 | 3.2 | 3.3 | 16.7 | 53 | 37 |
| Field Museum | 191348 | Negros, Philippines | *Z. nigrorum nigrorum* | nigrorum | 9.3 | 7 | 3.7 | 3 | 16.1 | 51.5 | 38 |
| Field Museum | 246424 | Negros, Philippines | *Z. nigrorum nigrorum* | nigrorum | 10.4 | 7.2 | 3.1 | 3.2 | 18.5 | 53.5 | 43 |
| Field Museum | 251501 | Negros, Philippines | *Z. nigrorum nigrorum* | nigrorum | 10.2 | 7.8 | 3.7 | 3 | 17.8 | 54.5 | 43 |
| Field Museum | 20247 | Cagayancillo, Philippines | *Z. nigrorum richmondi* | nigrorum | 11.1 | 8.7 | 3.4 | 3.6 | 17.9 | 57 | 46 |
| Field Museum | 20248 | Cagayancillo, Philippines | *Z. nigrorum richmondi* | nigrorum | 10.5 | 8 | 3.1 | 3.4 | 20.8 | 57 | 38 |
| Field Museum | 257639 | Sri Lanka | *Z. palpebrosus egregius* | palpebrosus | 8.7 | 6.3 | 2.6 | 2.8 | 15.9 | 51.5 | 37 |

| | | | | | | | | | | | |
|---|---|---|---|---|---|---|---|---|---|---|---|
| Field Museum | 257640 | Sri Lanka | *Z. palpebrosus egregius* | palpebrosus | 9 | 6.4 | 2.7 | 3 | 14.2 | 56.5 | 41 |
| Field Museum | 257641 | Sri Lanka | *Z. palpebrosus egregius* | palpebrosus | 9.1 | 6.6 | 2.6 | 2.9 | 14.1 | 52 | 40 |
| Field Museum | 257645 | Sri Lanka | *Z. palpebrosus egregius* | palpebrosus | 8.3 | 6 | 2.5 | 2.5 | 15.4 | 52.5 | 39 |
| Field Museum | 244012 | Ghats, India | *Z. palpebrosus nilgiriensis* | palpebrosus | 11.1 | 8.2 | 3.2 | 3 | 15.3 | 53 | 42 |
| Field Museum | 244019 | Ghats, India | *Z. palpebrosus nilgiriensis* | palpebrosus | 10.9 | 7.5 | 3.1 | 3 | 13.5 | 56 | 41 |
| Field Museum | 244020 | Ghats, India | *Z. palpebrosus nilgiriensis* | palpebrosus | 10.4 | 7.4 | 3.4 | 2.9 | 15 | 55 | 44 |
| Field Museum | 244021 | Ghats, India | *Z. palpebrosus nilgiriensis* | palpebrosus | 9.9 | 6.9 | 2.9 | 2.9 | 14.7 | 54 | 43 |
| Field Museum | 244022 | Ghats, India | *Z. palpebrosus nilgiriensis* | palpebrosus | 10.8 | 7.9 | 3 | 2.9 | 16.7 | 56 | 43 |
| Field Museum | 243993 | Kathiawar, India | *Z. palpebrosus occidentis* | palpebrosus | 7.9 | 5.8 | 2.8 | 2.8 | 15.5 | 53.5 | 39 |
| Field Museum | 243996 | Kathiawar, India | *Z. palpebrosus occidentis* | palpebrosus | 8.6 | 6 | 2.8 | 2.6 | 15.5 | 55 | 40 |
| Field Museum | 243997 | Kathiawar, India | *Z. palpebrosus occidentis* | palpebrosus | 8.7 | 6.3 | 2.9 | 2.5 | 14.4 | 53.5 | 39 |
| Field Museum | 243998 | Kathiawar, India | *Z. palpebrosus occidentis* | palpebrosus | 8.2 | 6 | 3.1 | 2.6 | 15.5 | 54 | 39 |
| Field Museum | 244004 | Kathiawar, India | *Z. palpebrosus occidentis* | palpebrosus | 8.4 | 6.4 | 2.8 | 3.2 | 16.2 | 53 | 40 |
| Field Museum | 244054 | North Central India | *Z. palpebrosus occidentis* | palpebrosus | 8.5 | 6.5 | 2.8 | 3.1 | 15 | 54 | 39 |
| Field Museum | 243980 | Nepal | *Z. palpebrosus palpebrosus* | palpebrosus | 9.4 | 6.5 | 2.7 | 2.8 | 14.7 | 52 | 37 |
| Field Museum | 243981 | Nepal | *Z. palpebrosus palpebrosus* | palpebrosus | 8.5 | 6.3 | 2.7 | 2.9 | 13.7 | 52.5 | 34 |
| Field Museum | 243987 | Nepal | *Z. palpebrosus palpebrosus* | palpebrosus | 9.4 | 6.8 | 2.8 | 2.4 | 14.9 | 52 | 40 |
| Field Museum | 244033 | Nepal | *Z. palpebrosus palpebrosus* | palpebrosus | 9.5 | 6.4 | 2.5 | 2.9 | 12.6 | 51 | 36 |
| Field Museum | 244036 | Nepal | *Z. palpebrosus palpebrosus* | palpebrosus | 9.3 | 6 | 2.5 | 2.4 | 13.8 | 53 | 33 |
| Field Museum | 244069 | Assam, India | *Z. palpebrosus palpebrosus* | palpebrosus | 9.7 | 7.2 | 2.8 | 2.9 | 14.7 | 53.5 | 39 |
| Field Museum | 80334 | Laos | *Z. palpebrosus williamsoni* | palpebrosus | 9.7 | 7 | 2.6 | 2.9 | 15.1 | 51.5 | 37 |
| Field Museum | 80337 | Laos | *Z. palpebrosus williamsoni* | palpebrosus | 9 | 6.7 | 2.6 | 2.9 | 15.7 | 52 | 34 |
| Field Museum | 91670 | Laos | *Z. palpebrosus williamsoni* | palpebrosus | 9.7 | 7.1 | 3 | 2.4 | 15 | 54.5 | 40 |
| Field Museum | 59474 | Fujian, China | *Z. simplex simplex* | simplex | 9.9 | 7.4 | 2.7 | 3 | 18.7 | 54.5 | 36 |
| Field Museum | 76219 | Vietnam | *Z. simplex simplex* | simplex | 9.3 | 7.6 | 2.4 | 2.6 | 16 | 53 | 37 |
| Field Museum | 76221 | Vietnam | *Z. simplex simplex* | simplex | 9.3 | 7.1 | 3 | 2.7 | 14.6 | 55.5 | 39 |
| Field Museum | 80307 | Vietnam | *Z. simplex simplex* | simplex | 9.3 | 7.8 | 2.9 | 2.8 | 17.1 | 56 | 43 |
| Field Museum | 80309 | Vietnam | *Z. simplex simplex* | simplex | 9.3 | 7.2 | 2.9 | 2.3 | 19.5 | 55 | 42 |

| | | | | | | | | | | | |
|---|---|---|---|---|---|---|---|---|---|---|---|
| Field Museum | 80311 | Vietnam | *Z. simplex simplex* | simplex | 9 | 7.5 | 2.8 | 2.5 | 14.9 | 56.5 | 39 |
| Field Museum | 80312 | Vietnam | *Z. simplex simplex* | simplex | 9.4 | 6.8 | 2.7 | 2.3 | 16.3 | 55 | 37 |
| Field Museum | 80317 | Vietnam | *Z. simplex simplex* | simplex | 10 | 7.3 | 2.7 | 3.1 | 17.5 | 56 | 39 |
| Field Museum | 80318 | Vietnam | *Z. simplex simplex* | simplex | 9.1 | 6.8 | 2.9 | 2.7 | 18.8 | 54 | 44 |
| Field Museum | 109065 | Sichuan, China | *Z. simplex simplex* | simplex | 10.1 | 6.7 | 2.9 | 3 | 16.4 | 57 | 38 |
| National Museum | 335147 | Sichuan, China | *Z. erythropleurus* | erythropleurus | 9.53 | 7.22 | 2.8 | 2.93 | 15.08 | 62.5 | 39 |
| National Museum | 535696 | Thailand | *Z. erythropleurus* | erythropleurus | 9.39 | 6.58 | 2.92 | 3.23 | 15.59 | 61 | 39 |
| National Museum | 535697 | Thailand | *Z. erythropleurus* | erythropleurus | 10.16 | 7.11 | 2.66 | 2.64 | 16.67 | 60.5 | 29 |
| National Museum | 535698 | Thailand | *Z. erythropleurus* | erythropleurus | 9.64 | 7.3 | 2.73 | 3.19 | 15.55 | 60 | 35 |
| National Museum | 535699 | Thailand | *Z. erythropleurus* | erythropleurus | 9.11 | 6.67 | 2.9 | 2.37 | 16.35 | 59.5 | 38 |
| National Museum | 581661 | Mindanao, Philippines | *Z. everetti basilanicus* | everetti | 8.86 | 6.93 | 3.29 | 3.3 | 17.31 | 62 | 40 |
| National Museum | 581677 | Mindanao, Philippines | *Z. everetti basilanicus* | everetti | 9.9 | 6.57 | 3.22 | 2.95 | 16.11 | 54.5 | 42 |
| National Museum | 211094 | Bohol, Philippines | *Z. everetti boholensis* | everetti | 10.02 | 6.89 | 3.22 | 3.21 | 16.3 | 55 | 41 |
| National Museum | 578001 | Leyte, Philippines | *Z. everetti boholensis* | everetti | 10.63 | 7.26 | 3.42 | 3.3 | 15.59 | 61 | 38 |
| National Museum | 578004 | Leyte, Philippines | *Z. everetti boholensis* | everetti | 10.1 | 7.17 | 3.6 | 3.56 | 15.7 | 53.5 | 41 |
| National Museum | 578008 | Leyte, Philippines | *Z. everetti boholensis* | everetti | 10.25 | 7.84 | 3.65 | 3.29 | 15.62 | 54.5 | 35 |
| National Museum | 315717 | Sulu, Philippines | *Z. everetti mandibularis* | everetti | 9.86 | 7.15 | 3.08 | 3.55 | 15.5 | 53 | 40 |
| National Museum | 485424 | Borneo | *Z. everetti tahanensis* | everetti | 8.26 | 5.79 | 2.48 | 3.22 | 13.82 | 52.5 | 31 |
| National Museum | 383201 | Iwoto, Japan | *Z. japonicus alani* | japonicus OCEANIC | 12.36 | 7.88 | 3.42 | 3.33 | 20.56 | 61 | 46 |
| National Museum | 383202 | Iwoto, Japan | *Z. japonicus alani* | japonicus OCEANIC | 11.12 | 7.63 | 3.33 | 3.33 | 19.94 | 61.5 | 44 |
| National Museum | 627873 | Kitadaitojima, Japan | *Z. japonicus daitoensis* | japonicus CONTINENTAL | 11.04 | 7.98 | 3.15 | 3.49 | 17.47 | 58.5 | 44 |
| National Museum | 91558 | Honshu, Japan | *Z. japonicus japonicus* | japonicus CONTINENTAL | 10.5 | 8.07 | 3.3 | 3.28 | 16.68 | 60 | 37 |
| National Museum | 96110 | Hokkaido, Japan | *Z. japonicus japonicus* | japonicus CONTINENTAL | 10.98 | 7.53 | 2.82 | 2.66 | 16.68 | 57 | 39 |
| National Museum | 405520 | Okinawa, Japan | *Z. japonicus loochooensis* | japonicus CONTINENTAL | 9.55 | 7.17 | 3.07 | 3.43 | 16.46 | 60.5 | 41 |
| National Museum | 405521 | Okinawa, Japan | *Z. japonicus loochooensis* | japonicus CONTINENTAL | 9.72 | 7.05 | 3.04 | 3.03 | 17.65 | 56 | 39 |
| National Museum | 405522 | Okinawa, Japan | *Z. japonicus loochooensis* | japonicus CONTINENTAL | 10.08 | 7.18 | 3.08 | 3.02 | 17.79 | 59.5 | 37 |
| National Museum | 405525 | Okinawa, Japan | *Z. japonicus loochooensis* | japonicus CONTINENTAL | 11.2 | 7.61 | 2.95 | 3.25 | 18.17 | 58.5 | 40 |
| National Museum | 111658 | Oshima, Japan | *Z. japonicus stejnegeri* | japonicus OCEANIC | 12.39 | 9.1 | 3.69 | 2.98 | 19.34 | 62.5 | 45 |

| | | | | | | | | | | | |
|---|---|---|---|---|---|---|---|---|---|---|---|
| National Museum | 582851 | Batan, Philippines | *Z. meyeni batanis* | meyeni | 10.15 | 6.95 | 3.1 | 3.6 | 16.9 | 55 | 40 |
| National Museum | 582852 | Batan, Philippines | *Z. meyeni batanis* | meyeni | 11.09 | 8 | 3.51 | 3.68 | 18.88 | 55 | 34 |
| National Museum | 582853 | Batan, Philippines | *Z. meyeni batanis* | meyeni | 10.41 | 7.19 | 3.16 | 3.53 | 17.68 | 55.5 | 41 |
| National Museum | 582854 | Batan, Philippines | *Z. meyeni batanis* | meyeni | 11.67 | 8.41 | 3.68 | 3.68 | 18.17 | 57.5 | 39 |
| National Museum | 606561 | Lanyu Island, Taiwan | *Z. meyeni batanis* | meyeni | 10.66 | 7.67 | 3.21 | 3.48 | 18.08 | 59 | 39 |
| National Museum | 606567 | Lanyu Island, Taiwan | *Z. meyeni batanis* | meyeni | 10.6 | 7.33 | 3.69 | 3.64 | 20.79 | 56 | 38 |
| National Museum | 606570 | Green Island, Taiwan | *Z. meyeni batanis* | meyeni | 10.85 | 8.07 | 3.52 | 3.34 | 19.65 | 58 | 41 |
| National Museum | 202406 | Mindoro, Philippines | *Z. montanus halconensis* | montanus NORTH | 10.16 | 7.74 | 2.88 | 3.48 | 16.11 | 55 | 39 |
| National Museum | 582067 | Negros, Philippines | *Z. montanus pectoralis* | montanus SOUTH | 10.41 | 7.46 | 2.85 | 3.28 | 17.26 | 56.5 | 38 |
| National Museum | 582068 | Negros, Philippines | *Z. montanus pectoralis* | montanus SOUTH | 10.35 | 7.2 | 3.18 | 3.52 | 19 | 53 | 40 |
| National Museum | 192202 | Mindanao, Philippines | *Z. montanus vulcani* | montanus SOUTH | 10.01 | 7.39 | 2.97 | 3.23 | 17.4 | 59 | 41 |
| National Museum | 192203 | Mindanao, Philippines | *Z. montanus vulcani* | montanus SOUTH | 11.08 | 7.99 | 3.33 | 3.57 | 16.82 | 62 | 38 |
| National Museum | 607560 | Luzon, Philippines | *Z. nigrorum aureiloris* | nigrorum | 10 | 6.73 | 2.93 | 2.94 | 14.49 | 52 | 35 |
| National Museum | 192701 | Masbate, Philippines | *Z. nigrorum nigrorum* | nigrorum | 9.74 | 7.6 | 3.12 | 3.82 | 15.64 | 55 | 42 |
| National Museum | 315706 | Panay, Philippines | *Z. nigrorum nigrorum* | nigrorum | 10.06 | 7.27 | 3.26 | 3.39 | 13.84 | 52 | 38 |
| National Museum | 192696 | Cagayancillo, Philippines | *Z. nigrorum richmondi* | nigrorum | 11.16 | 8.03 | 3.54 | 3.64 | 19.05 | 58.5 | 43 |
| National Museum | 192697 | Cagayancillo, Philippines | *Z. nigrorum richmondi* | nigrorum | 11.03 | 7.7 | 3.26 | 3.4 | 16.36 | 58.5 | 46 |
| National Museum | 86143 | Hong Kong | *Z. simplex simplex* | simplex | 7.72 | 5.07 | 2.89 | 3.37 | 16.13 | 55 | 37 |
| National Museum | 86144 | Hong Kong | *Z. simplex simplex* | simplex | 9.17 | 6.97 | 2.86 | 2.76 | 15.88 | 55.5 | 36 |
| National Museum | 86145 | Hong Kong | *Z. simplex simplex* | simplex | 9.06 | 7.32 | 2.69 | 3.14 | 16.1 | 54.5 | 37 |

Supplementary table 5: Linear morphological measurements recorded from 354 museum study skins. Additional data was collected from skins where poor condition or positioning prevented accurate measurements for one or more traits. These data are not shown here and were excluded from subsequent analyses. Culmen length was measured as exposed culmen from the tip of the bill to the point where the bill meets the forehead plumage. Bill length from nares-to-tip was measured from the distal end of the right nare to the tip of the bill. Bill depth and width were measured at the nares. Tarsometatarsus (tarsus) length was measured from the base of the phalanges to the tibiotarsal joint. Wing cord was measured unflattened with a wing rule. Tail length was measured from the body to the tip of the longest rectrix by placing the same wing rule in the center of the tail. Individual study skins were grouped by species and geography. The ‘japonicus CONTINENTAL’ group includes those populations on the main islands of Japan, the Korean Peninsula and adjacent islands, the Ryukyu archipelago, and Daito Islands. The 'japonicus OCEANIC' group included subspecies of Z. japonicus found on Japanese oceanic island chains in the Pacific including *Z. japonicus stejnegeri* in the Izu Islands and *Z. japonicus alani* in the Ogasawara and Volcano Islands. No specimens from either introduced populations in Hawaii or Southern California were included in any morphometric analyses. The 'montanus NORTH' group included those subspecies of *Z. montanus* on the islands of Luzon (*Z. montanus whiteheadi*) and Mindoro (*Z. montanus halconensis*) in the northern Philippines. The 'montanus SOUTH' group included subspecies and geographic locales for *Z. montanus* found in the Visayas in the central Philippines, Mindanao and adjacent islands in the southern Philippines, and Indonesia. The 'nigrorum CS' group represents the *Z. nigrorum catarmanensis* subspecies found on the island of Camiguin Sur in the southern Philippines and discovered in phylogenetic analyses in this study to be mismatched in a clade with *Z. montanus* rather than with other *Z. nigrorum* samples.

| Locus | Locus type | Individuals | Number of sequences | Length | Segregating sites | π | Number of haplotypes | Haplotype diversity | θ per locus | θ per locus variance | θ per site | Tajima's D | Tajima's D p-value | R2 test | R2 test p-value |
|---|---|---|---|---|---|---|---|---|---|---|---|---|---|---|---|
| COI | mt | 132 | 132 | 646 | 119 | 0.021 | 52 | 0.95 | 21.81 | 103.93 | 0.034 | -1.27 | 0.205 | 0.05 | 0.077 |
| ND2 | mt | 143 | 143 | 978 | 235 | 0.020 | 57 | 0.92 | 42.45 | 103.93 | 0.043 | -1.72 | 0.085 | 0.04 | 0.008 |
| B3GNT9 | cd | 87 | 174 | 724 | 28 | 0.003 | 30 | 0.86 | 4.88 | 2.04 | 0.007 | -1.07 | 0.286 | 0.04 | 0.020 |
| BIRC2 | cd | 88 | 176 | 559 | 23 | 0.001 | 32 | 0.87 | 4.00 | 1.49 | 0.007 | -1.50 | 0.134 | 0.02 | 0.000 |
| CCDC | cd | 87 | 174 | 727 | 23 | 0.002 | 26 | 0.74 | 4.01 | 1.50 | 0.006 | -1.48 | 0.140 | 0.03 | 0.001 |
| GPRC6 | cd | 87 | 174 | 651 | 14 | 0.001 | 20 | 0.66 | 2.44 | 0.72 | 0.004 | -1.11 | 0.267 | 0.02 | 0.000 |
| LACTBL | cd | 89 | 178 | 686 | 37 | 0.001 | 41 | 0.75 | 6.43 | 3.16 | 0.009 | -1.35 | 0.176 | 0.01 | 0.000 |
| USP38 | cd | 91 | 182 | 714 | 17 | 0.001 | 31 | 0.87 | 2.94 | 0.93 | 0.004 | -0.78 | 0.438 | 0.03 | 0.002 |

Locus type denoted as mitochondrial (mt) or nuclear protein coding (cd). For phased nuclear DNA alignments the number of sequences is 2x the number of individuals sequenced. π is per site nucleotide diversity. Haplotype diversity calculated according to Nei and Tajima (1981). θ per locus and per locus variance in θ are estimated from the number of segregating sites (Watterson estimator). θ per site is calculated by dividing the θ per locus by the sequence length. Tajima's D and the Ramos-Onsins and Rozas (R2) tests are used to test for neutralilty and population growth, repsectively, at each locus. Population genetic parameters for each locus were estimated in the pegas and ape R packages

Supplementary table 6: Summary of Sanger sequenced loci statistics.

| | *Z. erythropleurus* | *Z. everetti* | *Z. japonicus* | *Z. japonicus* OGASAWARA | *Z. meyeni* | *Z. montanus* INDONESIA | *Z. montanus* NORTH | *Z. montanus* PALAWAN | *Z. montanus* SOUTH | *Z. nigrorum* | Z. palpebrosus |
|---|---|---|---|---|---|---|---|---|---|---|---|
| ***Z. everetii*** | 884 / 15,429<br>5.7% | | | | | | | | | | |
| ***Z. japonicus*** | 576 / 15,658<br>3.7% | 1,286 / 15,475<br>8.3% | | | | | | | | | |
| ***Z. japonicus* OGASAWARA** | 649 / 15,658<br>4.1% | 1,374 / 15,475<br>8.9% | 0 / 15,704<br>0.0% | | | | | | | | |
| ***Z. meyeni*** | 467 / 15,658<br>3.0% | 1,146 / 15, 475<br>7.4% | 4 / 15,704<br>0.0003% | 12 / 15,704<br>0.0008% | | | | | | | |
| ***Z. montanus* INDONESIA** | 646 / 15,565<br>4.2% | 1,388 / 15,382<br>9.0% | 169 / 15,611<br>1.1% | 237 / 15,611<br>1.5% | 76 / 15,611<br>0.005% | | | | | | |
| ***Z. montanus* NORTH** | 496 /15,647<br>3.2% | 1,170 / 15,464<br>7.6% | 27 / 15,693<br>0.002% | 44 / 15,693<br>0.003% | 1 / 15,693<br>0.0001% | 82 / 15,600<br>0.5% | | | | | |
| ***Z. montanus* PALAWAN** | 612 / 15,613<br>3.9% | 1,340 / 15,430<br>8.7% | 147 / 15,659<br>1.0% | 181 / 15,659<br>1.2% | 62 / 15,659<br>0.004% | 208 / 15,566<br>1.3% | 73 / 15,648<br>0.005% | | | | |
| ***Z. montanus* SOUTH** | 451 / 15,658<br>2.9% | 1,086 / 15,475<br>7.0% | 28 / 15,704<br>0.002% | 51 / 15,704<br>0.003% | 6 / 15,704<br>0.0004% | 16 / 15,611<br>0.001% | 4 / 15,693<br>0.0003% | 46 / 15,659<br>0.003% | | | |
| ***Z. nigrorum*** | 740 / 15,480<br>4.8% | 119 / 15,403<br>0.008 % | 1,128 / 15,526<br>7.3% | 1,205 / 15,526<br>7.8% | 982/ 15,526<br>6.3% | 1,218 / 15,433<br>7.9% | 1,003 / 15,515<br>6.5% | 1,171 / 15,481<br>7.6% | 933 / 15,526<br>6.0% | | |
| ***Z. palpebrosus*** | 1,021 / 15,057<br>6.8% | 1,332 / 14,881<br>9.0% | 1,487 / 15,093<br>9.9% | 1,576 / 15,093<br>10.4% | 1,340 / 15,093<br>8.9% | 1,570 / 15,000<br>10.5% | 1,361 / 15,082<br>9.0% | 1,531 / 15,051<br>10.2% | 1,287 / 15,093<br>8.5% | 1,163 / 14,929<br>7.8% | |
| ***Z. simplex*** | 509 / 15,658<br>3.3% | 983 / 15,475<br>6.4% | 380 / 15,704<br>2.4% | 439 / 15,704<br>2.8% | 320 / 15,704<br>2.0% | 481 / 15,611<br>3.1% | 326 / 15,693<br>2.1% | 450 / 15,659<br>2.9% | 306 / 15,704<br>1.9% | 866 / 15,526<br>5.6% | 1,196 /15093<br>7.9% |

Supplementary table 7: Pairwise fixed differences out of the total number of loci (top) and percent fixed differences (bottom) for each paired comparison between taxa/populations. Fixed differences were calculated using the 'gl.fixed.diff' function in the *dartR* package in R.

| **CULMEN LENGTH** | **erythropleurus** | **everetti** | **japonicus CONTINENTAL** | **japonicus OCEANIC** | **meyeni** | **montanus NORTH** | **montanus SOUTH** | **nigrourm** | **nigrorum CS** | **palpebrosus** |
|---|---|---|---|---|---|---|---|---|---|---|
| **everetti** | **1.50E-03** | | | | | | | | | |
| **japonicus CONTINENTAL** | **1.60E-06** | **1.30E-03** | | | | | | | | |
| **japonicus OCEANIC** | **7.60E-09** | **3.60E-11** | **9.80E-06** | | | | | | | |
| **meyeni** | **5.40E-07** | **3.20E-03** | 1.00E+00 | **6.70E-07** | | | | | | |
| **montanus NORTH** | **2.90E-03** | 1.00E+00 | 4.70E-01 | **1.30E-05** | 1.00E+00 | | | | | |
| **montanus SOUTH** | **2.80E-06** | 9.00E-01 | 2.80E-01 | **1.20E-12** | 1.00E+00 | 1.00E+00 | | | | |
| **nigrourm** | 1.00E+00 | 2.50E-01 | **9.90E-06** | **2.00E-10** | **6.90E-06** | 1.60E-01 | **1.40E-04** | | | |
| **nigrorum CS** | **3.30E-05** | **6.60E-04** | 1.00E+00 | **4.20E-04** | 1.00E+00 | 6.80E-02 | 2.50E-01 | **9.40E-05** | | |
| **palpebrosus** | 1.00E+00 | **1.70E-03** | **9.30E-06** | **2.50E-08** | **5.20E-05** | 1.00E-02 | **3.80E-06** | 4.60E-01 | **3.10E-04** | |
| **simplex** | 1.00E+00 | **2.50E-05** | **1.90E-07** | **1.40E-09** | **2.60E-07** | **3.50E-04** | **1.10E-08** | 6.40E-01 | **2.70E-05** | 1.00E+00 |
| **NARE-TO-TIP BILL LENGTH** | | | | | | | | | | |
| **everetti** | **1.00E-06** | | | | | | | | | |
| **japonicus CONTINENTAL** | **1.00E-06** | 1.00E+00 | | | | | | | | |
| **japonicus OCEANIC** | **8.40E-09** | **3.90E-07** | **6.50E-05** | | | | | | | |
| **meyeni** | **9.20E-07** | 1.00E+00 | 1.00E+00 | **4.90E-03** | | | | | | |
| **montanus NORTH** | **2.30E-02** | 1.00E+00 | 1.00E+00 | **2.90E-04** | 5.30E-01 | | | | | |
| **montanus SOUTH** | **4.50E-09** | 1.00E+00 | 1.00E+00 | **5.30E-08** | 1.00E+00 | 1.00E+00 | | | | |
| **nigrourm** | **2.10E-03** | 1.00E+00 | 2.30E-01 | **6.90E-08** | **1.80E-02** | 1.00E+00 | 9.90E-02 | | | |
| **nigrorum CS** | **3.30E-05** | **1.80E-02** | **4.30E-02** | 1.00E+00 | 1.00E+00 | **3.50E-02** | **3.80E-02** | **9.10E-04** | | |
| **palpebrosus** | 1.00E+00 | **6.20E-05** | **5.20E-05** | **6.60E-08** | **3.20E-05** | 1.10E-01 | **7.00E-07** | **1.30E-02** | **1.40E-04** | |
| **simplex** | 1.10E-01 | **2.70E-03** | **1.40E-04** | **2.60E-09** | **3.00E-05** | 1.00E+00 | **3.40E-05** | 1.00E+00 | **1.90E-05** | 4.40E-01 |
| **BILL DEPTH** | | | | | | | | | | |
| **everetti** | **5.30E-08** | | | | | | | | | |
| **japonicus CONTINENTAL** | 3.00E-01 | **3.10E-06** | | | | | | | | |
| **japonicus OCEANIC** | **1.50E-07** | 1.00E+00 | **2.30E-06** | | | | | | | |
| **meyeni** | **4.00E-06** | 1.00E+00 | **1.00E-03** | 1.00E+00 | | | | | | |
| **montanus NORTH** | 1.00E+00 | **2.30E-04** | 1.00E+00 | **2.00E-04** | **3.30E-03** | | | | | |

| | | | | | | | | | | |
|---|---|---|---|---|---|---|---|---|---|---|
| **montanus SOUTH** | **3.30E-06** | **4.40E-04** | 7.40E-02 | **1.90E-05** | 3.30E-01 | 1.00E-01 | | | | |
| **nigrourm** | **2.60E-03** | **1.10E-02** | 1.00E+00 | **1.00E-03** | 4.40E-01 | 1.00E+00 | 1.00E+00 | | | |
| **nigrorum CS** | **5.90E-05** | 4.10E-01 | **1.10E-04** | 1.00E+00 | 5.00E-01 | **1.30E-03** | **4.00E-04** | **2.70E-03** | | |
| **palpebrosus** | 1.00E+00 | **5.20E-07** | 1.00E+00 | **7.10E-07** | **3.50E-05** | 1.00E+00 | **7.50E-05** | **1.60E-02** | **1.20E-04** | |
| **simplex** | 1.00E+00 | **3.30E-10** | **2.40E-02** | **5.80E-09** | **1.00E-07** | 1.00E+00 | **7.90E-09** | **4.50E-05** | **1.40E-05** | 1.00E+00 |
| **BILL WIDTH** | | | | | | | | | | |
| **everetti** | **2.10E-08** | | | | | | | | | |
| **japonicus CONTINENTAL** | 1.50E-01 | **8.10E-04** | | | | | | | | |
| **japonicus OCEANIC** | **1.80E-07** | 1.00E+00 | **2.20E-04** | | | | | | | |
| **meyeni** | **7.40E-07** | 2.40E-01 | **2.20E-05** | 1.00E+00 | | | | | | |
| **montanus NORTH** | 3.50E-01 | 1.00E+00 | 1.00E+00 | 8.30E-01 | 7.70E-02 | | | | | |
| **montanus SOUTH** | **7.90E-07** | 1.00E+00 | **3.80E-02** | 1.00E+00 | **4.80E-02** | 1.00E+00 | | | | |
| **nigrourm** | 5.50E-01 | **8.40E-04** | 1.00E+00 | **1.10E-03** | **1.20E-04** | 1.00E+00 | **1.90E-02** | | | |
| **nigrorum CS** | **5.10E-05** | 1.00E+00 | **8.50E-04** | 1.00E+00 | 1.00E+00 | 7.40E-01 | 1.00E+00 | **1.10E-02** | | |
| **palpebrosus** | 1.00E+00 | **1.60E-08** | 1.60E-01 | **4.00E-07** | **1.30E-06** | 1.00E+00 | **1.60E-06** | 1.00E+00 | **4.60E-05** | |
| **simplex** | 1.00E+00 | **6.30E-08** | 7.50E-01 | **2.90E-07** | **5.60E-07** | 1.00E+00 | **2.30E-06** | 1.00E+00 | **6.50E-05** | 1.00E+00 |
| **TARSUS LENGTH** | | | | | | | | | | |
| **everetti** | **7.10E-03** | | | | | | | | | |
| **japonicus CONTINENTAL** | **1.20E-05** | **4.90E-02** | | | | | | | | |
| **japonicus OCEANIC** | **1.30E-08** | **5.30E-10** | **3.70E-05** | | | | | | | |
| **meyeni** | **4.10E-06** | **1.40E-03** | 1.00E+00 | **1.50E-04** | | | | | | |
| **montanus NORTH** | 4.40E-01 | 1.00E+00 | 2.30E-01 | **7.40E-06** | **1.30E-02** | | | | | |
| **montanus SOUTH** | **6.10E-06** | 1.00E+00 | 1.30E-01 | **6.80E-13** | **5.70E-04** | 1.00E+00 | | | | |
| **nigrourm** | 1.40E-01 | 1.00E+00 | **8.90E-03** | **2.30E-09** | **2.90E-04** | 1.00E+00 | 8.00E-01 | | | |
| **nigrorum CS** | **1.00E-04** | **3.30E-04** | 4.50E-01 | 4.30E-02 | 1.00E+00 | **1.80E-03** | **3.80E-05** | **2.70E-04** | | |
| **palpebrosus** | **1.00E-02** | **6.00E-08** | **7.90E-08** | **1.20E-08** | **1.70E-07** | **4.80E-04** | **4.10E-11** | **3.30E-06** | **4.10E-05** | |
| **simplex** | 1.00E+00 | **1.50E-02** | **4.50E-05** | **5.30E-09** | **1.10E-05** | 1.00E+00 | **3.00E-04** | 1.00E+00 | **1.90E-04** | **2.10E-04** |
| **WING CORD** | | | | | | | | | | |

| | | | | | | | | | | |
|---|---|---|---|---|---|---|---|---|---|---|
| **everetti** | **1.40E-05** | | | | | | | | | |
| **japonicus CONTINENTAL** | 1.90E-01 | **2.20E-04** | | | | | | | | |
| **japonicus OCEANIC** | 1.60E-01 | **3.00E-10** | **1.60E-07** | | | | | | | |
| **meyeni** | **7.70E-04** | 1.00E+00 | 1.00E-01 | **1.80E-08** | | | | | | |
| **montanus NORTH** | **2.30E-04** | 1.00E+00 | **4.30E-05** | **2.20E-06** | **9.70E-03** | | | | | |
| **montanus SOUTH** | **6.80E-06** | 1.00E+00 | **1.60E-03** | **1.30E-12** | 1.00E+00 | 1.50E-01 | | | | |
| **nigrourm** | **1.10E-07** | **3.40E-04** | **7.80E-08** | **7.70E-11** | **4.30E-05** | 1.00E+00 | **2.00E-06** | | | |
| **nigrorum CS** | 1.00E+00 | **1.00E-03** | 1.00E+00 | **9.20E-03** | **2.50E-02** | **5.40E-04** | **4.00E-03** | **2.40E-05** | | |
| **palpebrosus** | **4.10E-06** | **3.40E-03** | **2.40E-07** | **1.10E-08** | **3.00E-05** | 1.00E+00 | **9.30E-05** | 1.00E+00 | **5.40E-05** | |
| **simplex** | **1.40E-05** | 1.00E+00 | **5.00E-05** | **1.40E-09** | 6.30E-01 | 1.00E+00 | 1.00E+00 | **2.10E-02** | **3.20E-04** | 7.30E-02 |
| **TAIL LENGTH** | | | | | | | | | | |
| **everetti** | 1.00E+00 | | | | | | | | | |
| **japonicus CONTINENTAL** | **8.50E-03** | **5.60E-03** | | | | | | | | |
| **japonicus OCEANIC** | **1.90E-06** | **1.30E-07** | 1.00E+00 | | | | | | | |
| **meyeni** | 1.00E+00 | 1.00E+00 | **3.90E-02** | **9.30E-06** | | | | | | |
| **montanus NORTH** | **3.20E-02** | **7.80E-03** | **5.60E-05** | **2.50E-06** | **6.10E-03** | | | | | |
| **montanus SOUTH** | 3.20E-01 | **4.60E-03** | **2.20E-08** | **2.50E-13** | **3.20E-02** | 1.00E+00 | | | | |
| **nigrourm** | 1.00E+00 | 4.40E-01 | **7.80E-05** | **3.30E-08** | 3.90E-01 | 1.00E+00 | 1.00E+00 | | | |
| **nigrorum CS** | 1.00E+00 | 1.00E+00 | 1.00E+00 | **5.30E-03** | 1.00E+00 | **3.10E-02** | 8.50E-02 | 6.30E-01 | | |
| **palpebrosus** | 1.00E+00 | 1.00E+00 | **1.30E-03** | **4.30E-07** | 1.00E+00 | 2.70E-01 | 1.00E+00 | 1.00E+00 | 1.00E+00 | |
| **simplex** | 1.00E+00 | 1.00E+00 | **1.20E-02** | **3.90E-06** | 1.00E+00 | **6.20E-02** | 1.70E-01 | 1.00E+00 | 1.00E+00 | 1.00E+00 |

Supplementary table 8: P-values from Bonferroni corrected (p.adjust.method="bonferroni") pairwise Wilcoxon comparisons of seven linear morphological measures across 354 specimens. Statistically significant comparisons (<0.05) highlighted in bold.
