## Supplementary figures and images for "The not-so-great speciator: Systematics and species limits in a rapid radiation, the Asiatic white-eye complex (*Zosterops spp.*)"

### Supplemental figure 1

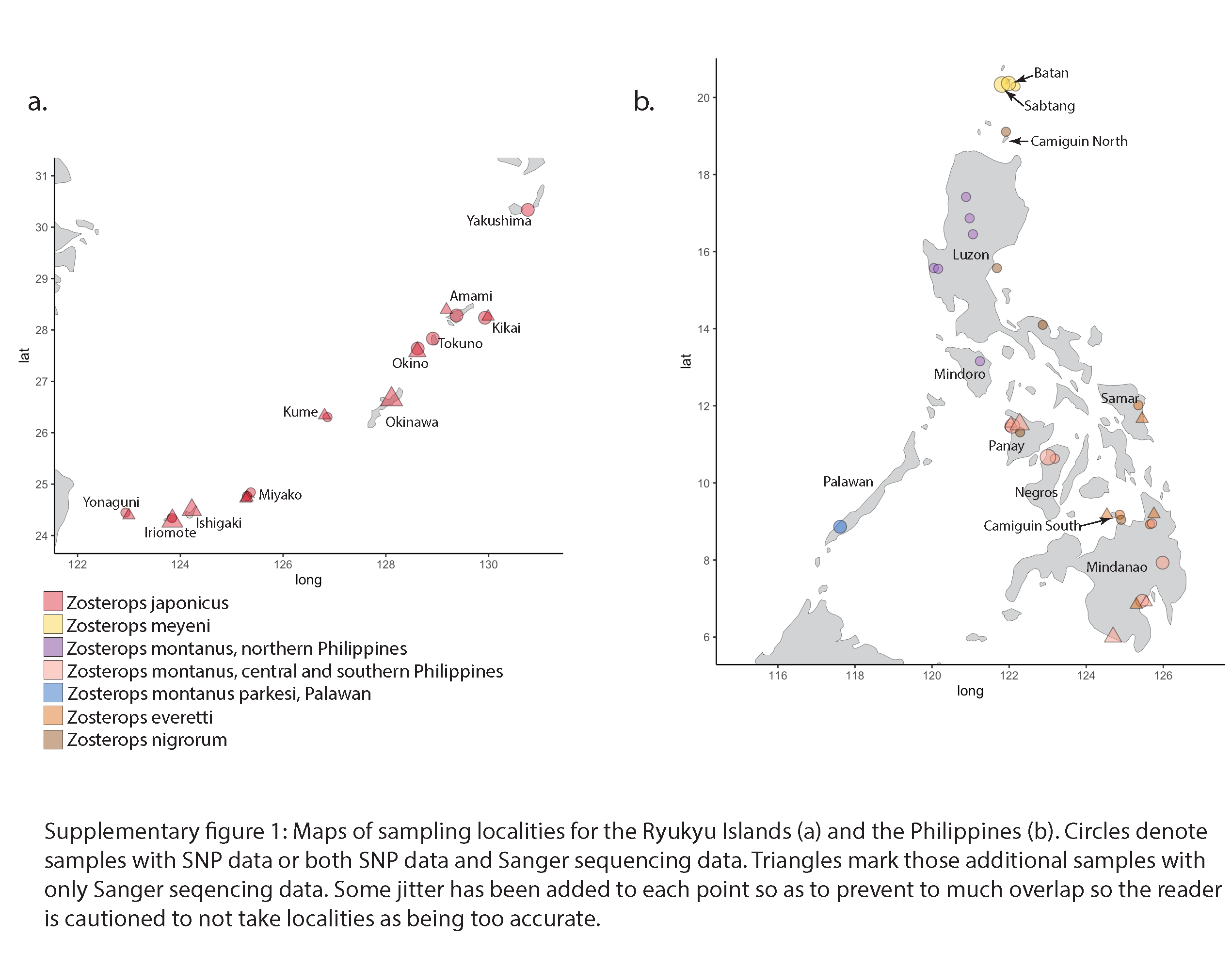

### Supplemental figure 2

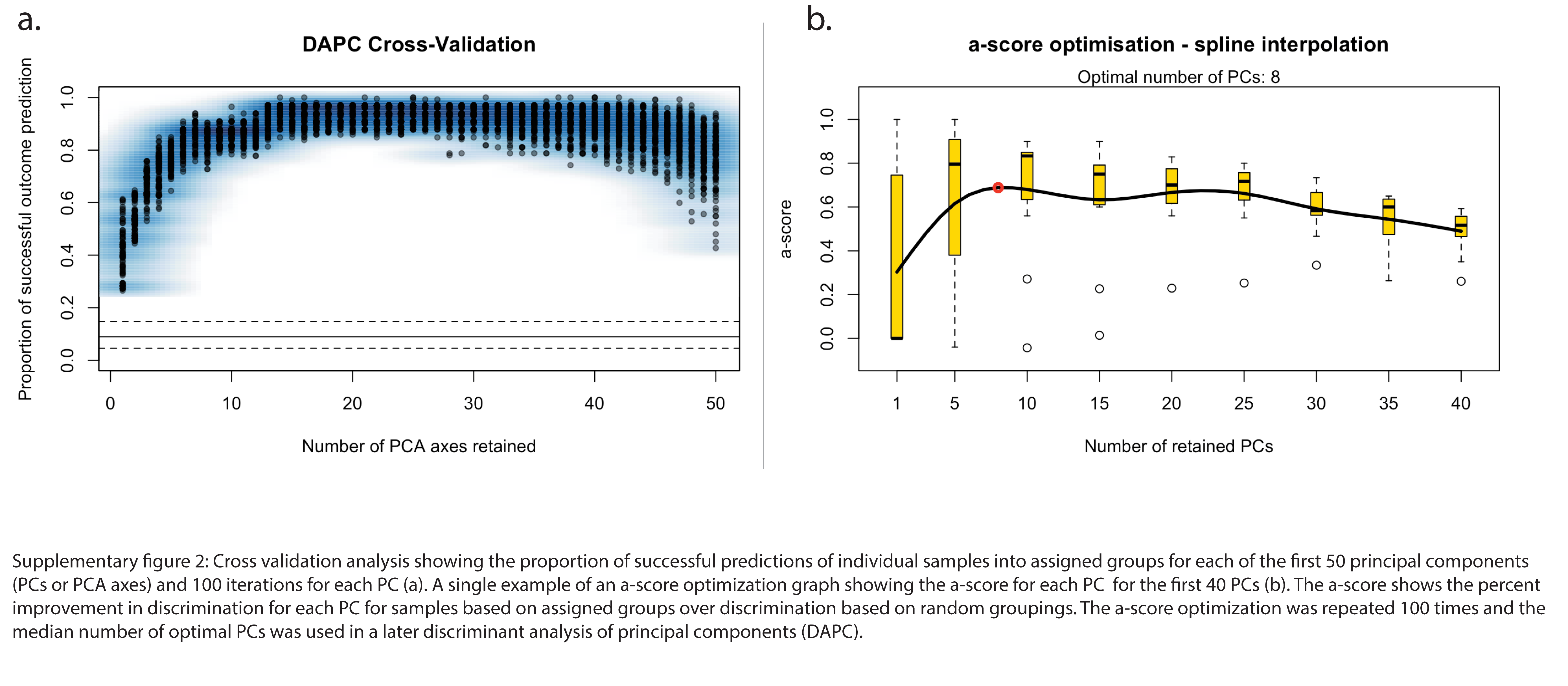

### Supplemental figure 3

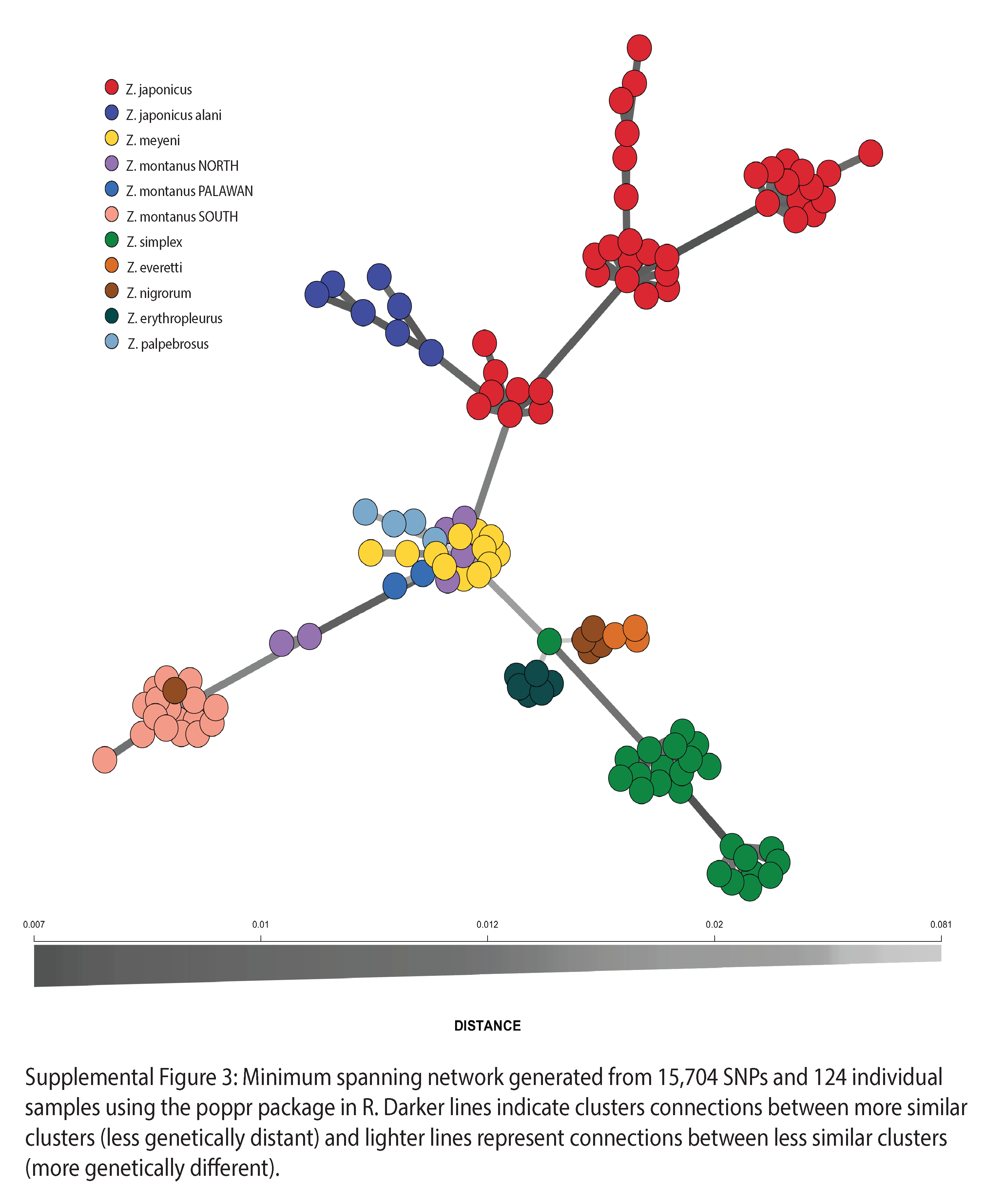

### Supplemental figure 4

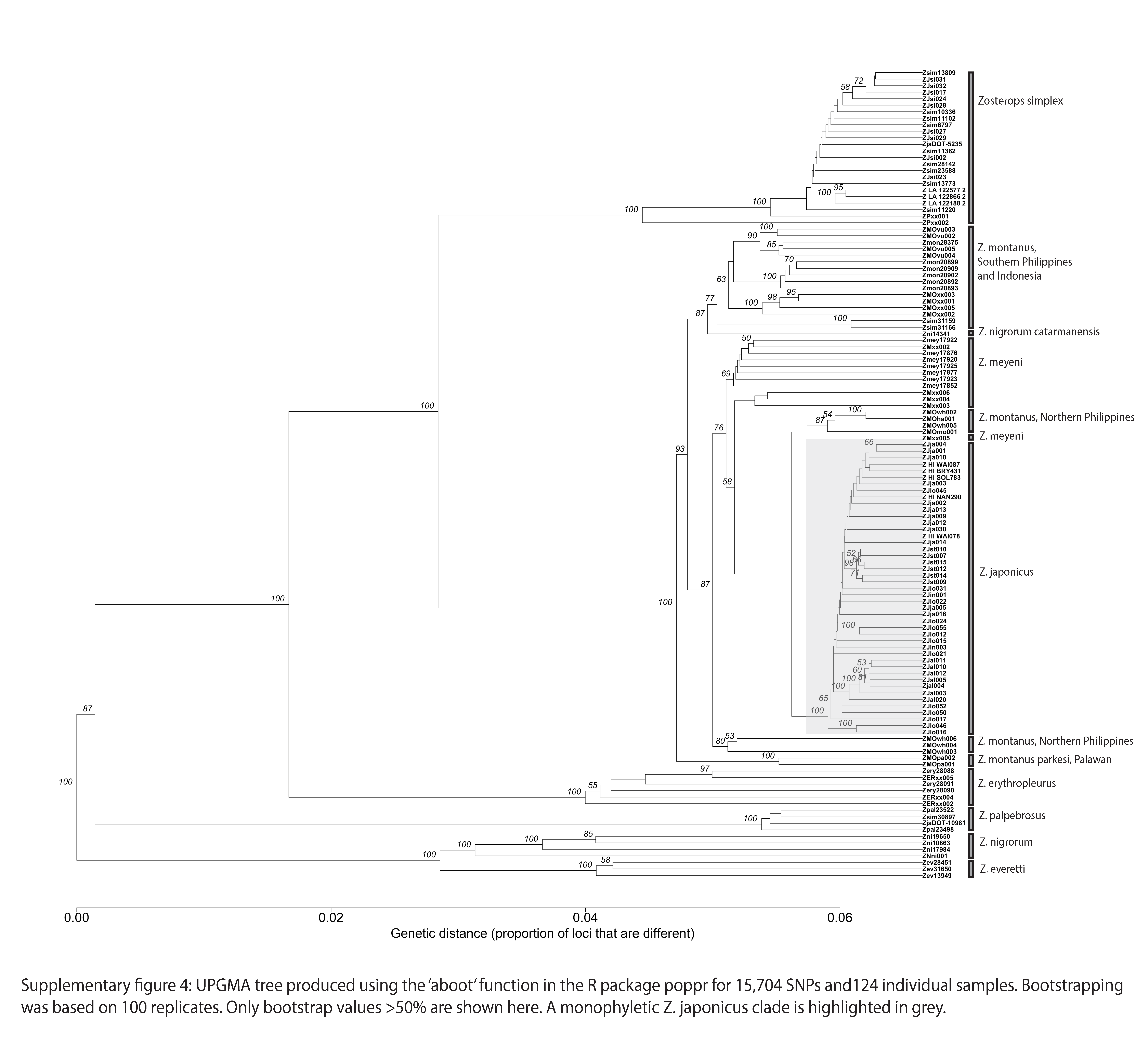

### Supplemental figure 5

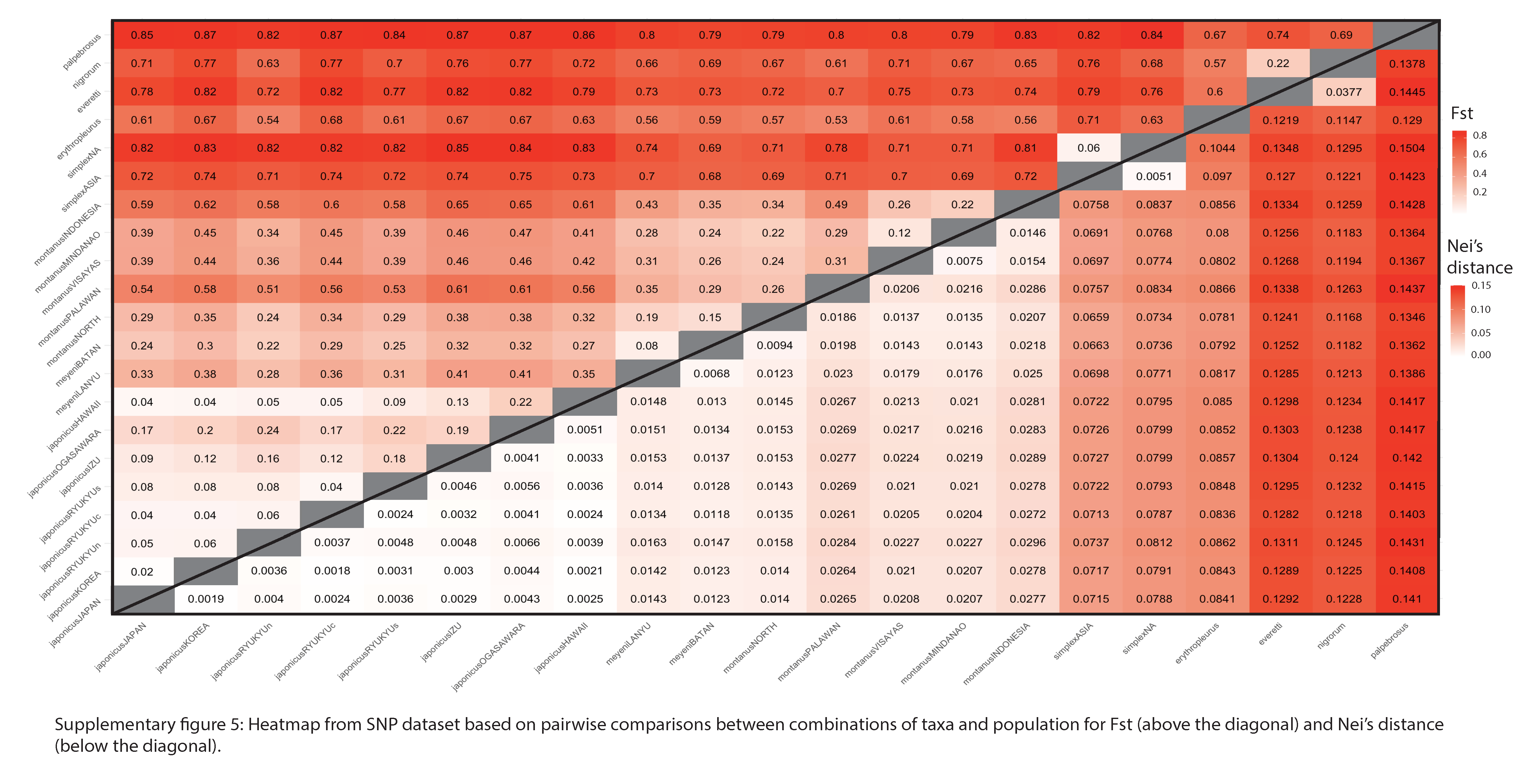

### Supplemental figure 6

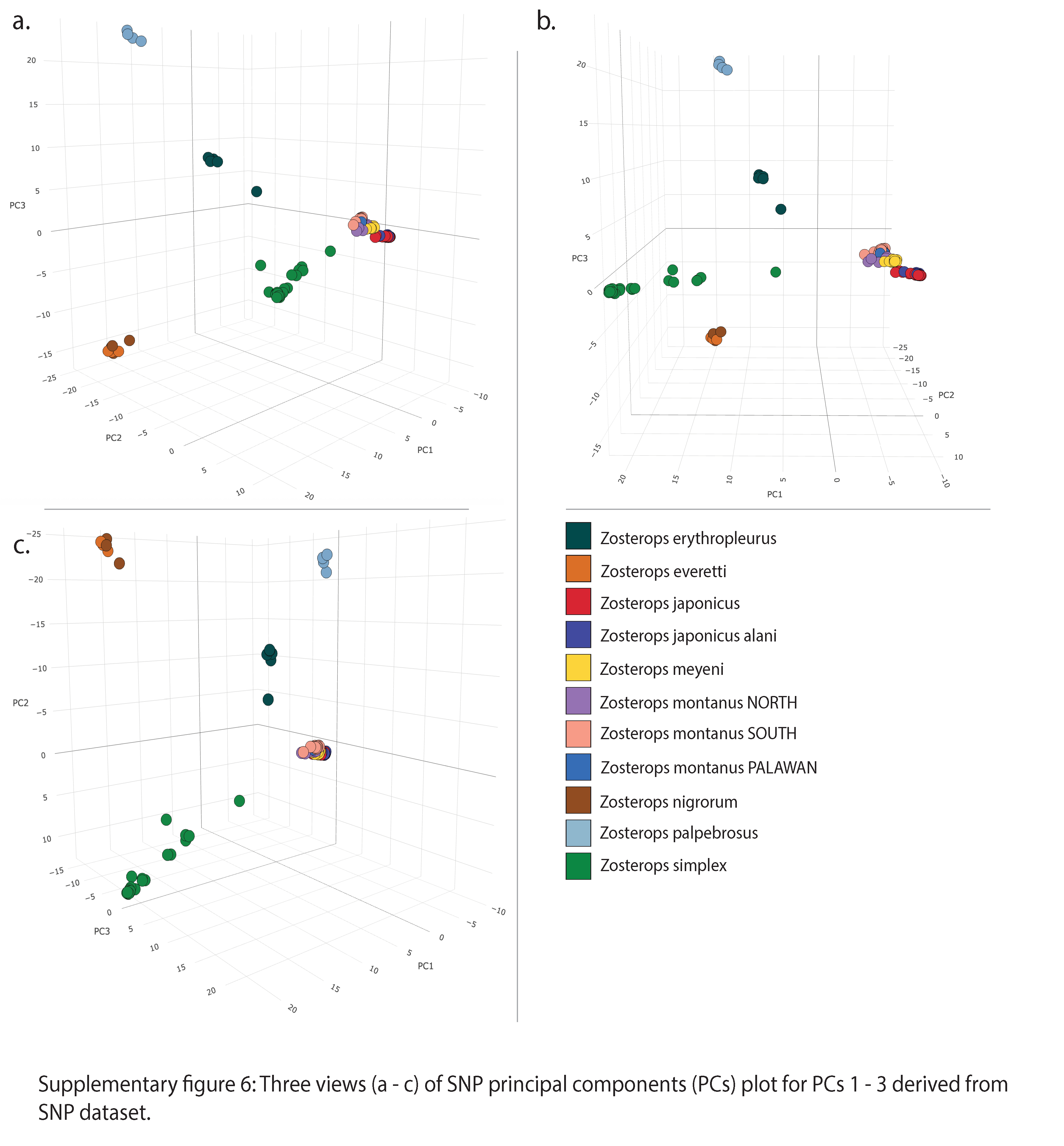

### Supplemental figure 7

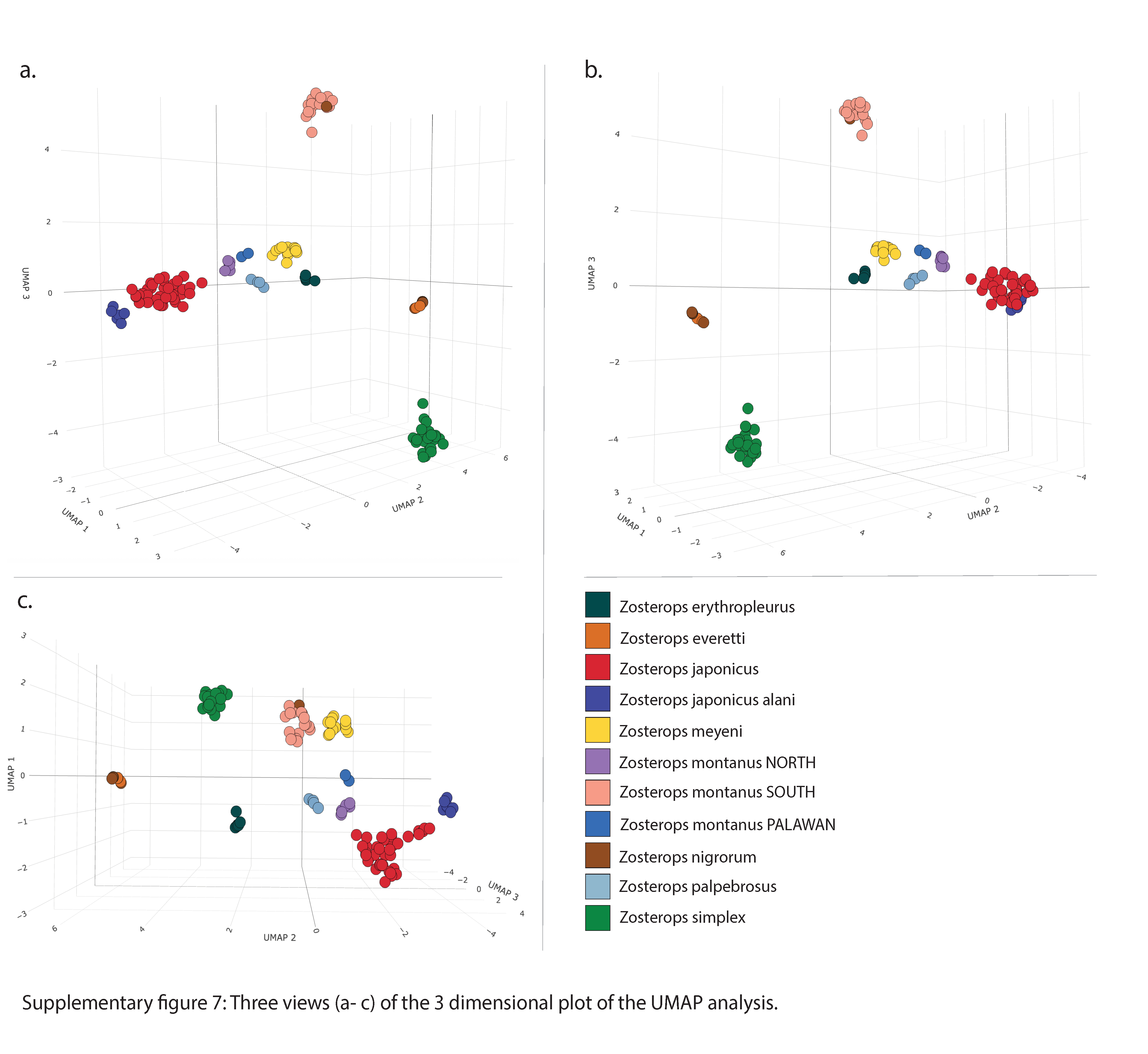

### Supplemental figure 8

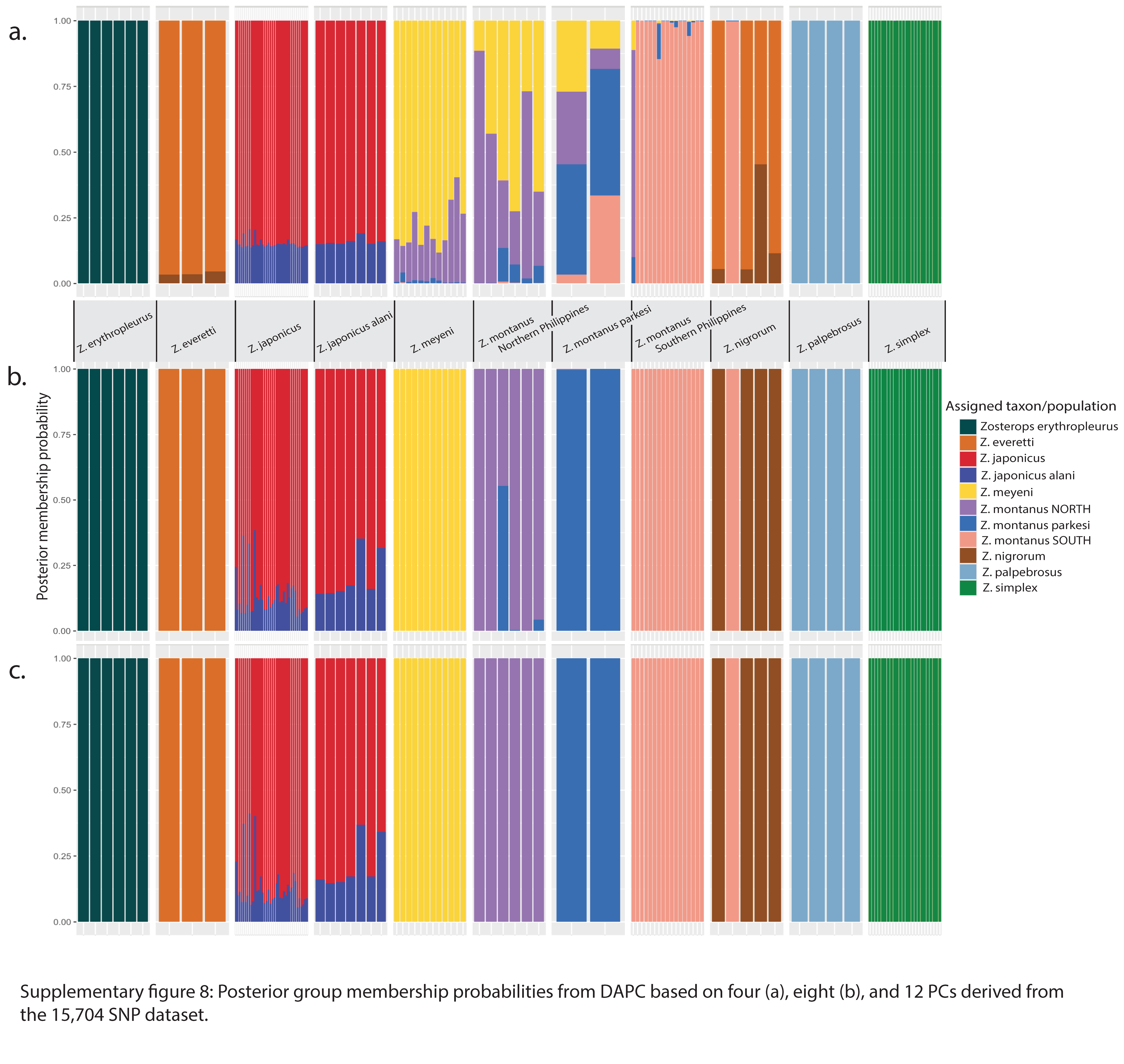

### Supplemental figure 9

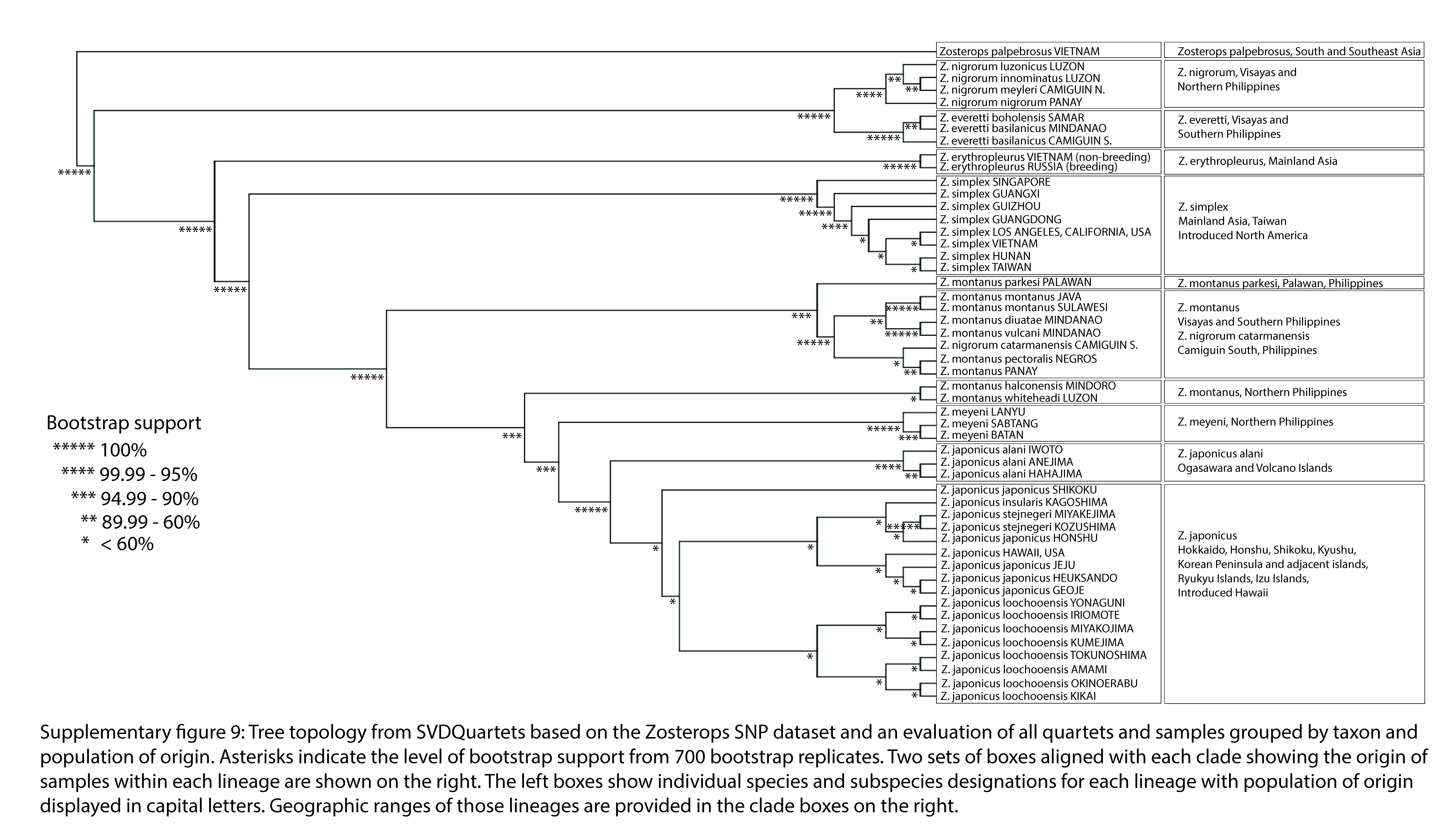

### Supplemental figure 10

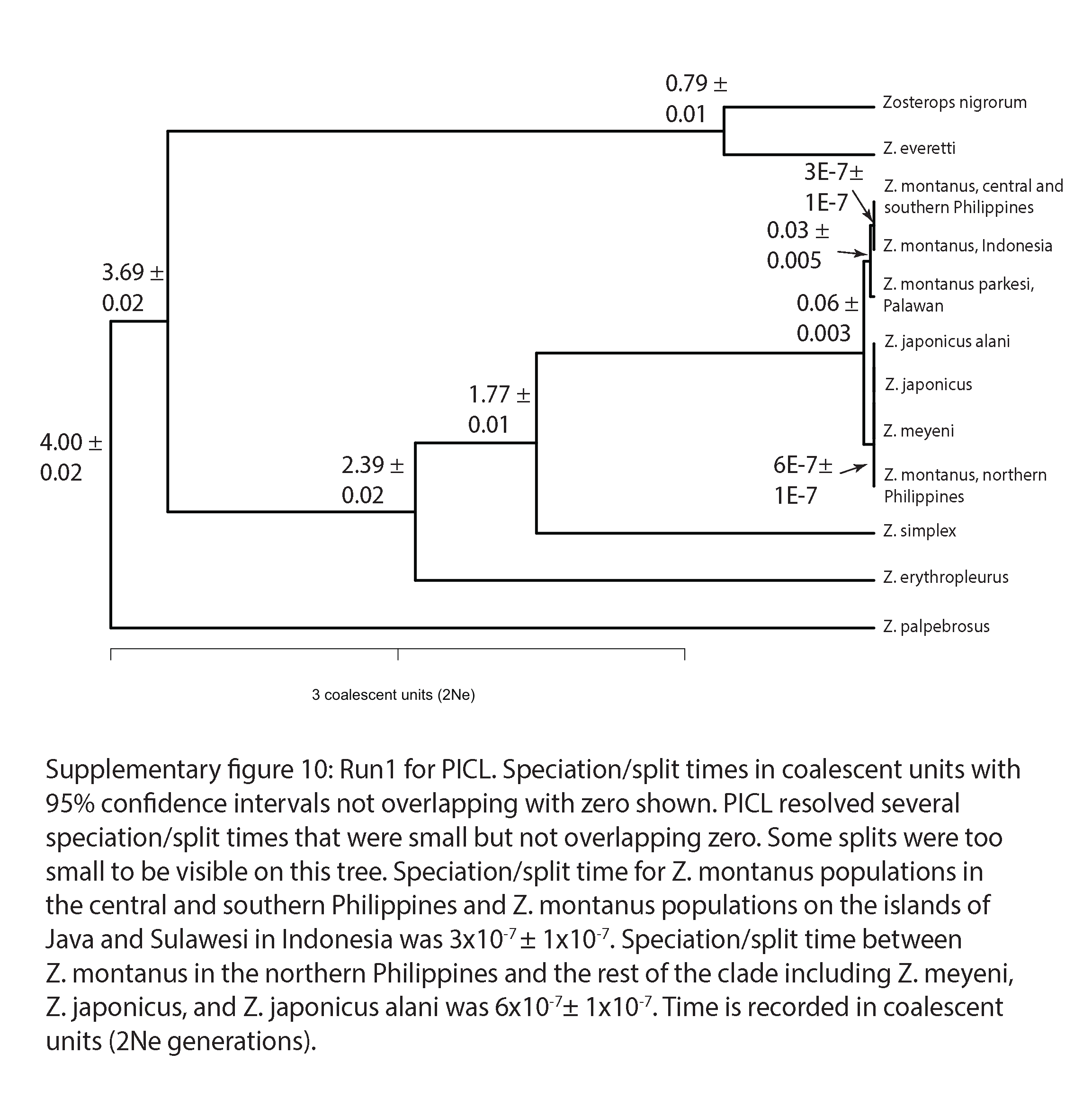
